# The plasmid-borne *hipBA* operon of *Klebsiella michiganensis* encodes a potent plasmid stabilization system

**DOI:** 10.1101/2024.07.25.605072

**Authors:** J Shutt-McCabe, KB Shaik, L Hoyles, G McVicker

**Affiliations:** Department of Biosciences, Nottingham Trent University, Clifton, Nottingham, NG11 8NS

**Keywords:** Plasmid, toxin-antitoxin, HipBA, *Klebsiella oxytoca* species complex, postsegregational killing

## Abstract

**Aims:** *Klebsiella michiganensis* is a medically-important bacterium that has been subject to relatively little attention in the literature. Interrogation of sequence data from *K. michiganensis* strains in our collection has revealed the presence of multiple large plasmids encoding type II toxin-antitoxin (TA) systems. Such TA systems are responsible for mediating a range of phenotypes including plasmid stability (“addiction”) and antibiotic persistence. In this work, we characterize the *hipBA* TA locus found within the *Klebsiella oxytoca* species complex (KoSC).

**Methods and Results:** The HipBA TA system is encoded on a plasmid carried by *K. michiganensis* PS_Koxy4, isolated from an infection outbreak. Employing viability and plasmid stability assays, we demonstrate that PS_Koxy4 HipA is a potent antibacterial toxin and that HipBA is a functional TA module contributing substantially to plasmid maintenance. Further, we provide *in silico* data comparing HipBA modules across the entire KoSC.

**Conclusions:** We provide the first evidence of the role of a plasmid-encoded HipBA system in stability of mobile genetic elements and analyze the presence of HipBA across the KoSC. These results expand our knowledge of both a common enterobacterial TA system and a highly medically-relevant group of bacteria.

**Impact Statement:** The HipBA TA system is typically encoded on bacterial chromosomes where it contributes to antimicrobial tolerance by interfering with translation during cellular stress. Here, we show that plasmid-encoded HipBA from a disease isolate of *Klebsiella michiganensis* is responsible for highly effective plasmid addiction; the first such evidence of a HipBA module contributing to plasmid stability. This has important implications for enteric pathogen evolution and horizontal gene transfer in the era of multidrug resistance.

## Introduction

*Klebsiella michiganensis* is a medically-important bacterium within the *Klebsiella oxytoca* species complex (KoSC) that is under-represented in the literature (Seiffert, et al. 2019, Chen, et al. 2020) and has recently been shown to encode the *til* biosynthetic gene cluster linked to production of cytotoxic metabolites that contribute to antibiotic-associated haemorrhagic colitis (Shibu, et al. 2021). Interrogation of Bakta-annotated sequence data for our in-house collection of KoSC strains revealed *K. michiganensis* PS_Koxy4 (Newberry, et al. 2023, Shibu, et al. 2021) to encode the type II toxin-antitoxin (TA) system HipBA on one of its plasmids, pPSKoxy4_2.

Type II TA systems are commonly found encoded on the mobile genetic elements (MGEs) and chromosomes of both Gram-positive and Gram-negative bacteria. They serve numerous roles within the bacterial cell and were originally characterized as postsegregational killing (PSK, otherwise known as “addiction”) systems (reviewed by Harms, et al. 2018). Such systems function through the interaction of a stable toxic protein, which causes bacterial cell lysis or cessation of growth, and a labile antitoxin protein that directly sequesters and abrogates the effect of the toxin but is itself actively and specifically degraded by cellular proteases. During PSK, loss of the MGE encoding a type II TA system allows the antitoxin pool to be rapidly depleted whilst the toxin remains stable, preventing the growth of any daughter cells that lack the MGE.

HipBA is a well-characterized type II TA system in *Escherichia coli* where it is commonly encoded in the chromosome of the organism. Chromosomal TA modules, rather than typically being considered PSK systems, are instead thought to contribute to persister cell formation (Harms, Maisonneuve and Gerdes 2016) and abortive infection as an anti-phage defence (Dy, et al. 2014). The *hipBA* operon is structured as a classical type II TA system, in which the antitoxin gene *hipB* precedes the toxin gene *hipA* within the operon. The HipA protein in *E. coli* is a kinase that may phosphorylate multiple members of the cellular translation machinery, e.g. elongation factor Tu (EF-Tu) and glutamyl-tRNA-synthetase, causing cessation of translation that leads to persistence (Kaspy, et al. 2013, Schumacher, et al. 2009, Bokinsky, et al. 2013). Interestingly, unlike the majority of type II TA families, there has been no prior experimental characterisation of plasmid-encoded HipBA systems and hence prior to this work it was unknown whether HipBA could serve as an effective PSK module to stabilize extrachromosomal MGEs.

Herein, we show that the HipA protein of *K. michiganensis* PS_Koxy4 is a potent antibacterial toxin that is effectively neutralized by its cognate partner, HipB. We provide evidence that HipA activity is dramatically increased in KoSC bacteria relative to *E. coli*, such that induced expression of *hipA* results in the rapid accumulation of toxin-inactivating mutations. We use a laboratory-constructed vector containing the natural IncA/C replicon of pPSKoxy4_2 to demonstrate that HipBA is a highly effective plasmid addiction system in both *E. coli* and *Klebsiella*. We also provide *in silico* data showing diverse plasmid-encoded HipBA variants are found across the KoSC, including a novel *Klebsiella* sp. (OXY-12; Ma, et al. 2021).

## Materials and Methods

### Bacterial strains and growth conditions

Bacteria were grown in lysogeny broth (LB, Lennox formulation; Sigma-Aldrich, UK) or plated onto solid lysogeny agar (LA; consisting of LB + 1.5 % (w/v) agar (Difco, UK)). Sucrose agar for plasmid stability testing contained 10 g/L tryptone (Sigma-Aldrich, UK), 5 g/L yeast extract (Sigma-Aldrich, UK), 10 % (w/v) sucrose (Sigma-Aldrich, UK) and 1.5 % (w/v) agar. Antibiotics (Sigma-Aldrich, UK) were added where required to the following final concentrations: ampicillin 100 μg/ml; chloramphenicol 30 μg/ml; neomycin 50 μg/ml. Unless otherwise stated, liquid cultures were grown with aeration at 37 °C and plates were incubated stationary at 37 °C. Bacteria were stored at −80 °C in 15 % (v/v) glycerol.

### DNA manipulation and cloning

Whole bacterial DNA was purified using the Wizard Genomic DNA Purification Kit (Promega, UK). Laboratory vectors and plasmid constructs (**Supplementary Table 1**) were purified using the Monarch Plasmid Miniprep Kit (New England Biolabs, UK). DNA fragments for cloning were amplified using Q5 DNA polymerase (New England Biolabs, UK), with oligonucleotide primers shown in **Supplementary Table 2**. If necessary to reduce cloning background growth, PCRs were digested overnight at 37 °C with *Dpn*I (New England Biolabs, UK). Amplicons or digest reactions were purified using the Wizard SV Gel and PCR Clean-Up System (Promega, UK). Isothermal DNA assembly was carried out via NEBuilder HiFi DNA Assembly (New England Biolabs, UK). Constructs and PCRs were analysed via agarose gel electrophoresis and Sanger (Source BioScience, UK) or Oxford Nanopore (plasmidsNG, UK) sequencing as appropriate.

### Toxicity assays

Toxicity and antitoxicity of TA system gene products were tested according to previous work (McVicker and Tang 2016). Cells carrying toxin and antitoxin constructs or empty vector controls were grown in LB + 0.2 % (w/v) glucose (Sigma-Aldrich, UK) overnight. These were subcultured to OD_600_ = 0.01 in fresh LB + 0.2 % (w/v) glucose and regrown to OD = 0.1-0.2, at which point cells were pelleted via centrifugation for 10 min at 4,500 ***g*** and the growth medium discarded. Cells were resuspended in prewarmed LB + 1 % (w/v) arabinose (Sigma-Aldrich, UK) to induce toxin expression. Immediately following induction, and again at each timepoint, a ten-fold dilution series was carried out in phosphate-buffered saline (PBS; Sigma-Aldrich, UK) and plated onto LA + 0.2 % (w/v) glucose to determine viable counts. At all points of the assay including plating, appropriate antibiotics were included to maintain the plasmid constructs.

### Plasmid stability assays

Stability of pSTAB vectors was tested according to previous work (McVicker, et al. 2019). Briefly, bactefria were streaked from freezer stocks onto LA containing no antibiotics. After 18-20 h growth, colonies were individually picked and suspended in 100 μl PBS. A ten-fold serial dilution was then carried out and plated separately onto both LA + neomycin and sucrose agar to determine the number of cells in the original colony that contain or lack the *sacB-neoR* marker, respectively.

### Genome sequences used in analyses

Details for RefSeq KoSC genome sequences (*n* = 861) used in this study are given in **Supplementary Table 3**. Quality of the genomes was assessed using CheckM2 (Chklovski, et al. 2023). Affiliation of the genome sequences with members of the KoSC was confirmed by average nucleotide analysis (ANI) (**Supplementary Table 3**) using FastANI v.1.3.3 (Jain, et al. 2018) against the genomes of reference sequences (**Table 1**), and phylogenetic analysis using PhyloPhlAn v3.1.68 (Asnicar, et al. 2020). The phylogenetic tree was visualized and annotated using iToL v6 (Letunic and Bork 2024). Multilocus sequence type (MLST) data were generated using the KoSC scheme, which covers *K. oxytoca*, *K. michiganensis*, *K. grimontii* and *K. pasteurii* (Herzog, et al. 2014, Jolley, Bray and Maiden 2018).

**Table 1.** Reference KoSC genomes used in this study.

| Taxon | Phylogroup (*bla<sub>oxy</sub>* group) | Type/representative strain | RefSeq accession |
| --- | --- | --- | --- |
| *Klebsiella grimontii* | Ko6 (OXY-6) | 06D021<sup>T</sup> | GCF_900200035.1 |
| *Klebsiella huaxiensis* | Ko8 (OXY-8) | WCHKI090001<sup>T</sup> | GCF_003261575.2 |
| *Klebsiella michiganensis* | Ko1 (OXY-1, OXY-5) | W14<sup>T</sup> | GCF_901556995.1 |
| *Klebsiella oxytoca* | Ko2 (OXY-2) | ATCC 13182<sup>T</sup> | GCF_900977765.1 |
| *Klebsiella pasteurii* | Ko4 (OXY-4) | SB6412<sup>T</sup> | GCF_902158725.1 |
| *Klebsiella spallanzanii* | Ko3 (OXY-3, OXY-9) | SB6411<sup>T</sup> | GCF_902158555.1 |
| "*Klebsiella* taxon 1" | (OXY-10) | 67 | GCF_003201885.1 |
| "*Klebsiella* taxon 2" | (OXY-11) | P620 | GCF_009707385.1 |
| "*Klebsiella* taxon 3" | (OXY-12) | RHBSTW-00484 | GCF_013705725.1 |

### Identification of plasmid and chromosomal contigs in genome data

Kraken2 2.0.7-beta was run against 861 genome sequences with the plasmid database described by Gomi, et al. (2021). For each genome sequence, contigs identified as unclassified (NCBI taxonomy ID 0) or root (NCBI taxonomy ID 1) by Kraken2 were discarded from further analyses as their taxonomic affiliation was uncertain; contigs identified as plasmid (NCBI taxonomy ID 36549) by the Kraken2 database were written to a new fasta file using Biostrings v2.72.1 (Pagès, et al. 2024); all other contigs (i.e. the chromosomal contigs) were written to a new fasta file using Biostrings v2.72.1.

### Identification of HipA and its partner sequences in plasmid and chromosomal sequence data

Both the plasmid and chromosomal sequences for each genome were run through SLING v2.0.1 using the default setting (Horesh, et al. 2018). As we were only interested in the HipBA-encoding sequences within the KoSC genomes in this study, data were filtered from the plasmid- and chromosomal-specific sequence files to retain only those genomes encoding one or more complete SLING hits for HipA and its partner sequence(s).

### Phylogenetic analysis of HipA (toxin) and HipB (antitoxin) sequences

The nucleotide sequences (i.e. HipA, upstream partner HipB sequence and/or downstream partner HipB sequence) of the complete hits from SLING were translated to protein sequences using Biostrings v2.72.1. Reference protein sequences of experimentally validated HipB and HipA (Guan, et al. 2024) were downloaded from UniProt or NCBI Protein (**Table 2**). All fasta files were imported into Geneious Prime v2022-07-07 and used to generate multiple-sequence alignments (MAFFT v7.490 (Katoh and Standley 2013), BLOSUM62 matrix). RAxML (v8 (Stamatakis 2014)) phylogenetic trees (-f a -x 1; 100 bootstraps) were generated from the multiple-sequence alignments. Newick format files were exported from Geneious and visualized using iToL v6 (Letunic and Bork 2024).

**Table 2.** Reference, experimentally validated HipBA sequences included in this study (all chromosomal)

| TADB3 ID | Strain | Protein accession |  | Family/domain | Reference(s) |
| --- | --- | --- | --- | --- | --- |
|  |  | HipB | HipA |  |  |
| TA000658 | *Shewanella oneidensis* MR-1 | Q8EIX4 | Q8EIX3 | hipBA/Xre-HipA | (Wen, et al. 2014) |
| TA001040 | *Escherichia coli* K-12 (MG1655) | P23873 | P23874 | hipBA/Xre-HipA | (Rotem, et al. 2010, Hansen, et al. 2012) |
| TA006108 | *Burkholderia pseudomallei* K96243 | Q63JY1 | Q63JY0 | hipBA/Xre-HipA | (Butt, et al. 2013, Ross, et al. 2019) |
| TA010038 | *Shewanella putrefaciens* CN-32 | ABP76632.1* | ABP76633.1* | hipBA/- | (Zhao, et al. 2022) |
| TA010075 | *Caulobacter crescentus* NA1000 | A0A0H3CAT0 | A0A0H3CA27 | hipBA/- | (Zhou, Eckart and Shapiro 2021, Huang, et al. 2020) |
| TA010076 | *Caulobacter crescentus* NA1000 | A0A0H3CAW1 | A0A0H3CA56 | hipBA/- | (Zhou, Eckart and Shapiro 2021, Huang, et al. 2020) |
| TA010098 | *Escherichia coli* Nissle 1917 | V0VAH2 | WP_038427096.1* | hipBA/- | (Xu, et al. 2020) |
| TA010104 | *Caulobacter crescentus* NA1000 | A0A0H3C5I2 | A0A0H3C5Z1 | hipBA/- | (Huang, et al. 2020) |
| TA010106 | *Xenorhabdus nematophila* ATCC 19061 | N1NPE2 | D3VDL4 | hipBA/- | (Yadav and Rathore 2020) |
| TA010195 | *Escherichia coli* BL21 (TaKaRa) | WP_001301023.1* | S1ELM3 | hipBA/- | (Maeda, et al. 2017) |

\*Downloaded from NCBI Protein; all other protein sequences downloaded from UniProt.

### Structural comparisons

Structural predictions were made from protein sequences using ColabFold (Mirdita, et al. 2022) and AlphaFold-multimer (Richard Evans, et al. 2021) via ChimeraX v1.8 (Meng, et al. 2023). Protein structures were aligned using the matchmaker command in ChimeraX.

## Results

### PS_Koxy4 carries a large IncA/C plasmid

The clinical isolate *K. michiganensis* PS_Koxy4 carries multiple plasmids (Newberry, et al. 2023). The 110 kb IncA/C plasmid pPSKoxy4_2 (**Figure 1**) possesses conjugation- and mobility-related genes, and putatively encodes the broad host range plasmid stabilising factor KfrA, a ParAB-like plasmid partitioning system and a range of DNA-modifying enzymes such as adenine/cytosine methyltransferases. These genes suggest that pPSKoxy4_2 may be a “helper plasmid” that aids in the conjugation and stable transfer and maintenance of both itself and the strain’s other plasmids, including perhaps the strain’s IncFII resistance plasmid, pPSKoxy4_1.

**Figure 1:**
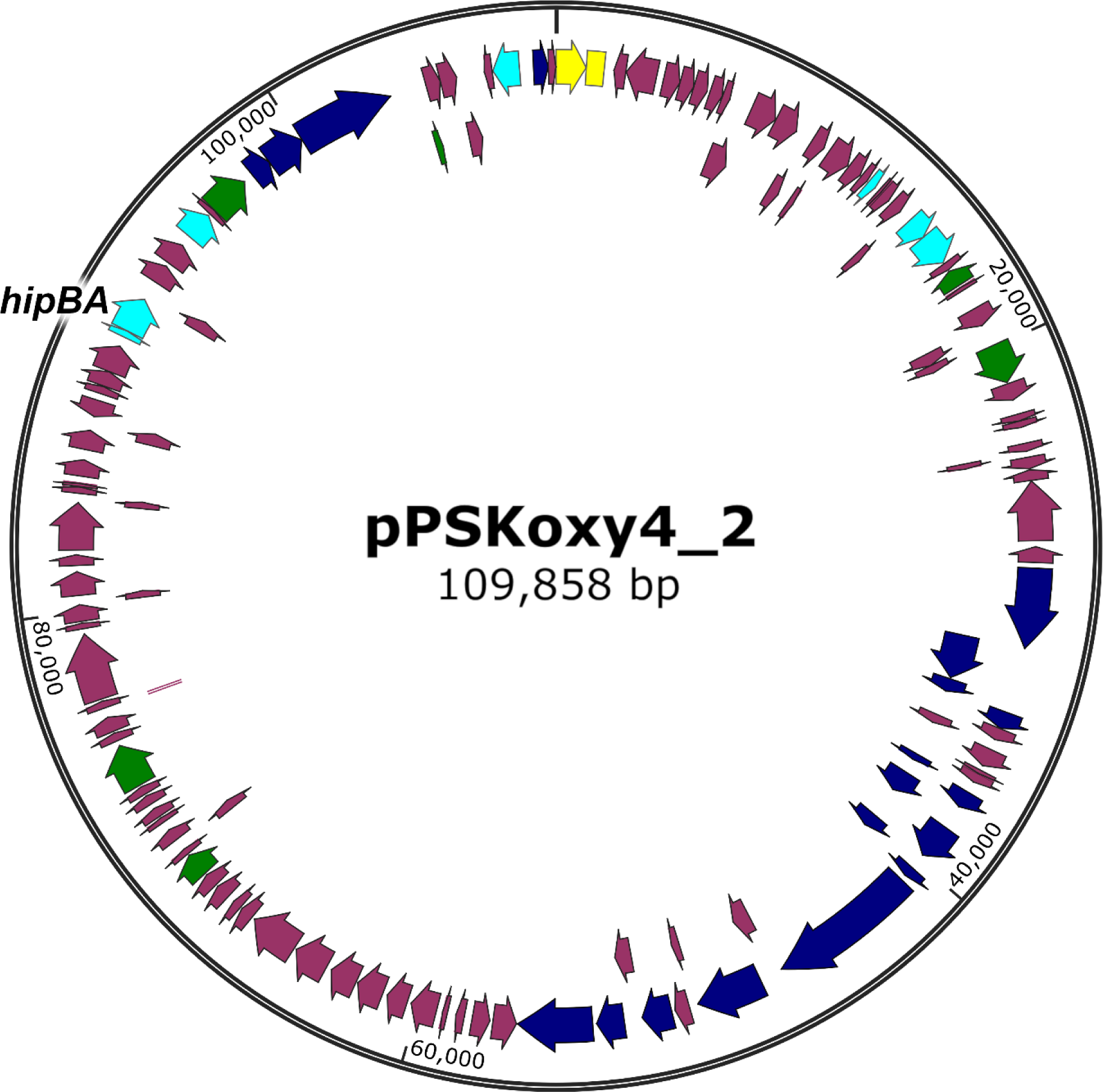
Map of the *K. michiganensis* IncA/C plasmid, pPSKoxy4_2. Yellow: plasmid replication. Dark blue: conjugation/transfer. Cyan: plasmid stability. Green: DNA modification/repair. The *hipBA* locus is indicated.

### pPSKoxy4_2 encodes a highly toxic HipBA system

HipBA is a TA system that is widespread in bacterial chromosomes and has been implicated in antibiotic persistence in multiple species (Semanjski, et al. 2018, Korch, Henderson and Hill 2003, Falla and Chopra 1998) and stability of a genomic island (Zhao, et al. 2022); however, it has not been characterized as a stabilising factor on bacterial plasmids. pPSKoxy4_2 encodes a HipBA variant with 43.93 % identity (54.46 % similarity) to the toxin protein sequence and 38.55 % identity (51.8 % similarity) to the antitoxin protein sequence from *E. coli* K-12 chromosomal HipBA (UniProt: P23873 and P23874). Other TA systems found in PSKoxy_4 include two CcdAB modules, encoded as distinct variants on pPSKoxy4_1 and the strain’s chromosome (Whelan 2022), and VapBC and ParDE, both encoded on the IncR plasmid pPSKoxy4_3. Notably, the ParE sequence seems to be disrupted by a premature stop codon, so only the first 32 amino acids (approximately one third of a canonical ParE protein depending upon the specific variant) are intact.

To establish whether *hipA*, *vapC* or *parE* from PSKoxy_4 encode functional toxins, we cloned the genes individually into the arabinose-inducible expression vector pBAD33 (Guzman, et al. 1995) and induced expression in *E. coli* NEB 5-alpha (**Figure 2A**). *ccdB* from pPSKoxy4_1 was also included as a known toxic control (Whelan 2022). Results demonstrated significant toxicity of CcdB as expected (*P* < 0.0001, viability 0.01 % of empty vector control at 180 min post induction), and also mild toxicity of VapC (*P* < 0.0001, viability 1.22 % at 180 min post induction). Interestingly, HipA showed extremely fast and effective toxicity versus the empty vector control (*P* < 0.0001, viability 0.40 % after only 15 min and lowest at 0.02 % after 60 min). The culture also showed slight recovery by the final timepoint (viability 0.08 % at 180 min; *P* < 0.0001 versus control), which has been observed previously in the case of highly effective toxins (McVicker and Tang 2016) and is likely due to mutation of the expression vector in surviving cells. Expressing *parE* did not give a significantly different result to the empty vector control (*P* > 0.98 throughout experiment), as expected from the gene’s truncation.

**Figure 2:**
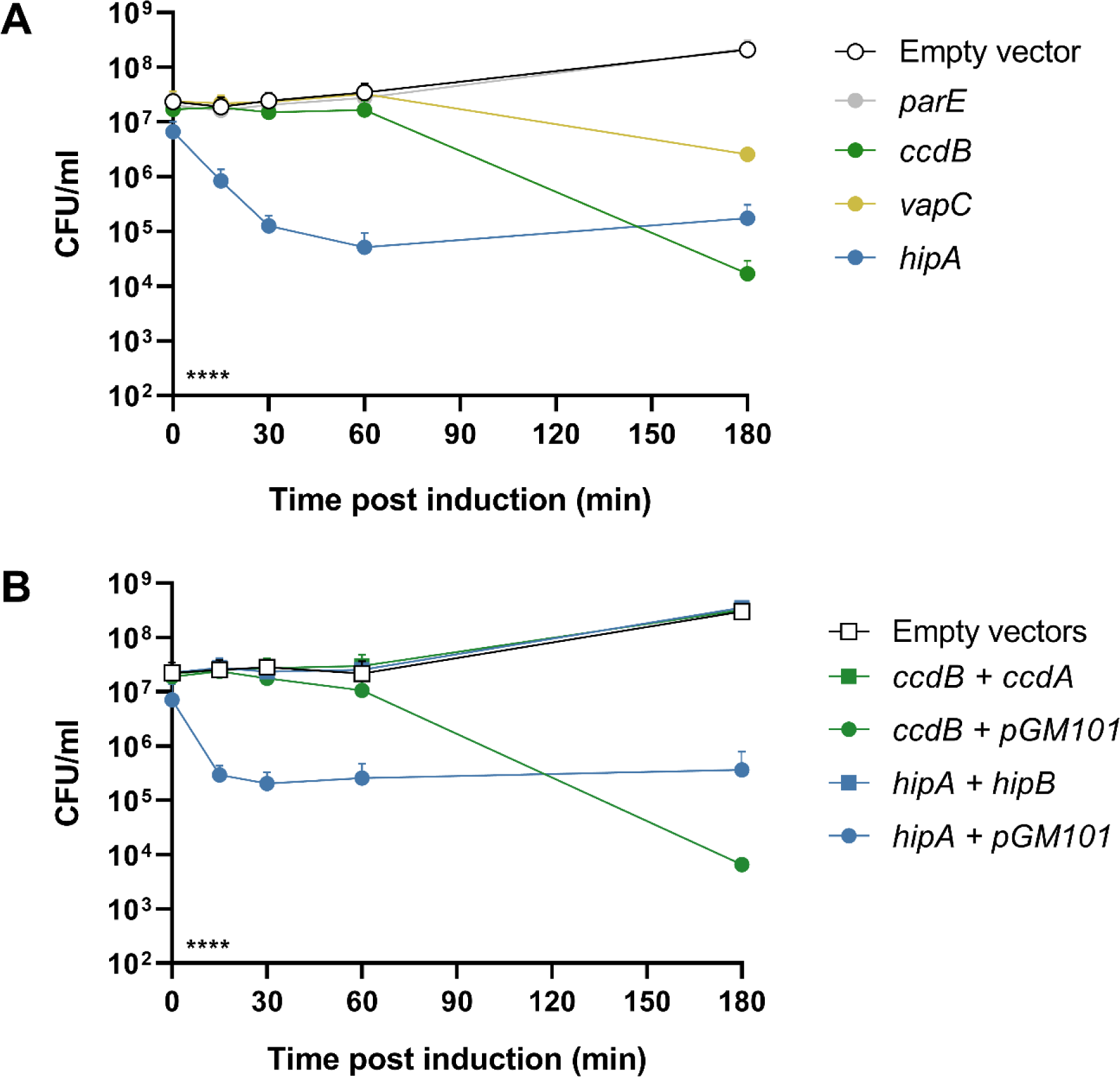
Characterisation of plasmid-borne TA systems in PS_Koxy4. Mean bacterial viability after induction of the indicated putative toxin genes from pBAD33. **A: Gene expression induced alone in *E. coli* NEB 5-alpha. B: Toxin gene expression induced in *E. coli* NEB 5-alpha in the presence (gene indicated) or absence (“pGM101”; empty vector) of cognate antitoxin genes.** Bars show standard error of the mean. Statistical analysis by two-way ANOVA: indicator in lower left shows the overall effect of strain in each experiment; **** *P* < 0.0001 (from *n* ≥ 3 biological replicates). Multiplicity-adjusted statistical analysis from Tukey’s post tests given in the main text.

To investigate the effects of the PSKoxy_4 HipBA TA system further, we cloned the *hipB* antitoxin gene onto the pBAD33-compatible vector pGM101 (McVicker and Tang 2016). The antitoxin’s native promoter was included upstream to take advantage of conditional cooperativity, i.e. the ability of the promoter to express a proportionate amount of antitoxin to the level of toxin in the cell (Cataudella, et al. 2012, Overgaard, et al. 2008). Results (**Figure 2B**) showed that the HipB antitoxin was able to abrogate HipA toxicity in *E. coli* for the entire duration of the experiment (*P* ≥ 0.2677 versus no-toxin control throughout), whilst HipA remained significantly toxic in the presence of the empty pGM101 (P ≤ 0.0179 versus no-toxin control, at all timepoints after induction).

### Induced expression of hipA in Klebsiella oxytoca generates nontoxic mutants

Ideally, toxicity assays should be conducted in a strain more similar to the native host of the TA system than *E. coli* NEB 5-alpha. To this end, we transformed plasmids into *K. oxytoca* NCTC 13727^T^ (originally isolated from human pharyngeal tonsil; Flügge 1886, Lautrop 1956), a strain of *Klebsiella* that does not carry plasmids or encode a HipBA TA module, and induced toxin expression to observe the effect on viability of KoSC bacteria (**Supplementary Figure 1**). Surprisingly, HipA did not significantly reduce viability relative to the control (*P* ≥ 0.0792 throughout experiment), whilst CcdB remained toxic (*P* < 0.0001 at 180 min post induction). Given the extremely high toxicity observed in *E. coli*, we suspected that a mutation may have arisen in the *hipA* expression vector whilst in *Klebsiella*. To confirm this, we extracted the expression plasmid from surviving *K. oxytoca* and sequenced the *hipA* ORF, which revealed a threonine to proline substitution at amino acid 175. We transformed this “*Klebsiella*-passaged” plasmid back into *E. coli* NEB 5-alpha, and also generated an *E. coli* strain carrying a plasmid with an identical mutation created *de novo* through site-directed mutagenesis of the original vector. In each case, toxicity was no longer observed relative to the empty vector control (**Figure 3**; *P* ≥ 0.1440 throughout experiment), whereas the original expression vector remained toxic upon induction (*P* < 0.0001), suggesting that the T175P substitution is sufficient to prevent HipA toxicity.

**Figure 3:**
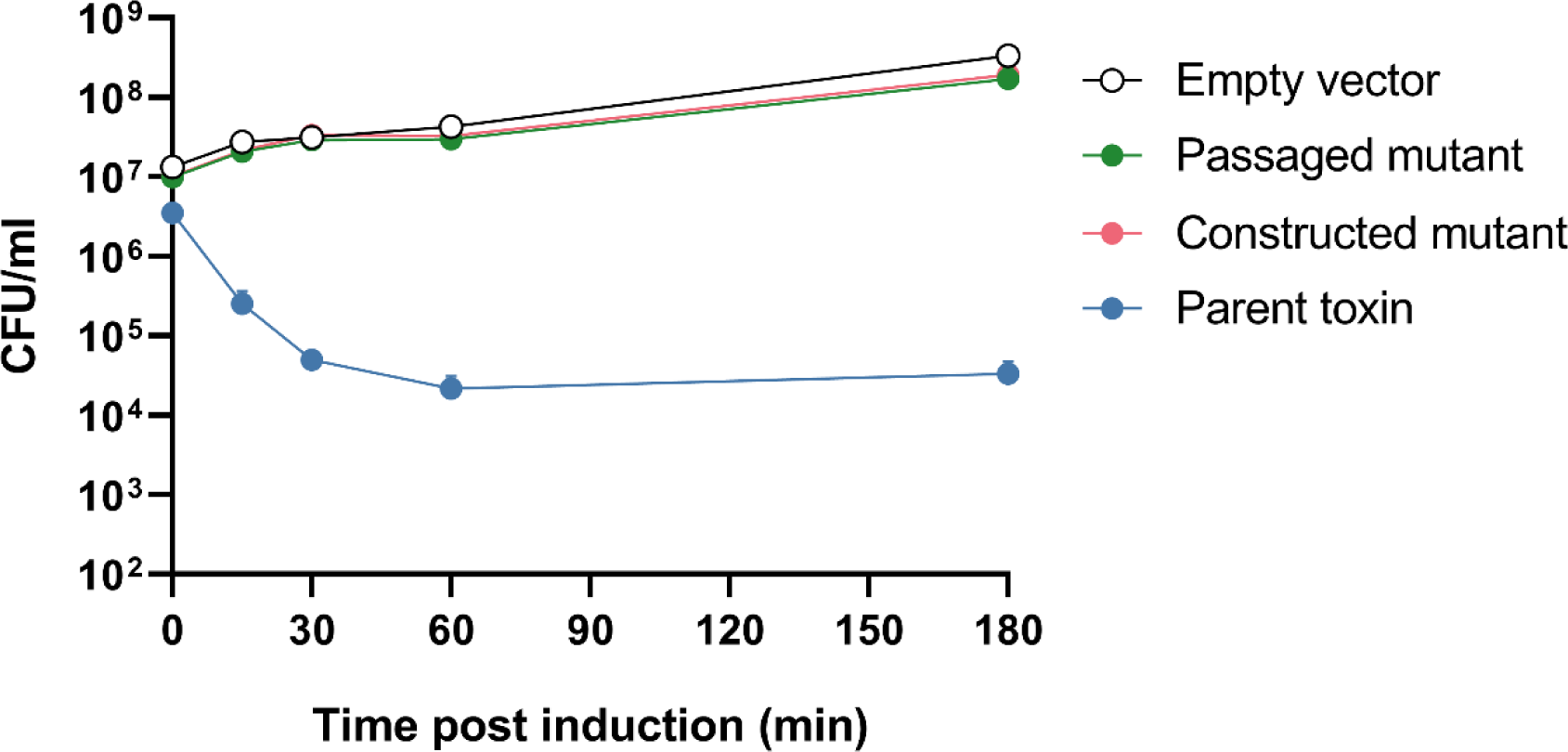
Generation of a non-toxic *hipA* mutant by passage of expression vector through *K. oxytoca*. Mean bacterial viability after induction of expression from pBAD33: “passaged mutant” is T175P variant extracted from *K. oxytoca* after a toxicity experiment; “constructed mutant” is T175P variant created through site-directed mutagenesis *in vitro*; “parent toxin” is the wild type variant. Gene expression induced in *E. coli* NEB 5-alpha. Bars show standard error of the mean. Statistical analysis by two-way ANOVA: indicator in lower left shows the overall effect of strain; **** *P* < 0.0001 (from *n* = 3 biological replicates). Multiplicity-adjusted statistical analysis from Tukey’s post tests given in the main text.

We reasoned that if this mutation occurred due to extremely high toxicity of HipA in *Klebsiella*, similar disabling mutations would be generated by further attempts to induce the *hipA* gene in this genus. To test this, we once again transformed the vector encoding wild-type HipA into *K. oxytoca* NCTC 13727^T^ and subjected several separate lineages of transformants to the same expression conditions under which the first mutant was produced. All three lineages produced a high number of surviving colonies (data not shown). Of the three plasmids generated by this experiment, one contained a threonine-proline substitution at a different site to the original mutation (T327P), one contained a frameshift mutation downstream of codon 234 due to a single-base deletion and one contained no identifiable mutations within the ORF, suggesting a mutation elsewhere on the vector or in the chromosome of the host.

Structural analyses of the wild-type and mutated (T175P or T327P) toxins demonstrated both mutations introduced similar changes in the structure of HipA, leaving the mutated structures more similar to one another than to the wild-type HipA (**Supplementary Figure 2**). That is, the root mean square deviation between wild-type HipA and the T175P and T327P mutants was 0.297 Å and 0.316 Å (across all 437 pruned atom pairs: 5.144 and 5.111), respectively, while the T175P and T327P mutants shared a root mean square deviation of 0.148 Å (across all 437 pruned atom pairs: 0.148).

### The PS_Koxy4 IncA/C plasmid is readily lost upon tellurite treatment

It is possible to disrupt TA system function in the original host strain by supplying the relevant antitoxin from an exogenous vector (Radnedge, et al. 1997 and our unpublished data), removing the need for the *in situ* antitoxin gene and hence “short-circuiting” the TA addiction mechanism. However, PS_Koxy4 is resistant to many commonly-used selection markers (Shibu, et al. 2021), so to successfully transform the strain, we constructed a pGM101 variant conferring tellurite resistance via the *kilA-telAB* cassette from plasmid pMo130-TelR (Amin, et al. 2013). We produced this vector (pGM101_tel_) both with and without the *hipB* gene under control of its native promoter. After construction in *E. coli* NEB 5-alpha, transformation of each vector into PS_Koxy4 was successful, conferring resistance to 10 μg/ml potassium tellurite. Transformants were grown overnight in LB containing 10 μg/ml tellurite, stocked and then plated after serial dilution to obtain single colonies. Colony PCR targeting the *hipA* gene was then used to detect the presence or absence of pPSKoxy4_2 after this short period of growth. However, we were surprised to find that the cultures had lost the IncA/C plasmid in up to 95 % of tested colonies regardless of whether they carried the control or HipB-encoding vector.

Given this result, we grew the PS_Koxy4 parent strain without tellurite and tested for plasmid presence in the same way, using primers targeting both *hipA* and the IncA/C replication origin, to assess the natural rate of plasmid loss. We observed 100 % plasmid retention in the tested colonies within the timeframe of the original experiment, therefore we reasoned that pPSKoxy4_2 is naturally stable and had been lost in our initial testing either due to tellurite-induced stress or due to incompatibility with the pGM101 replicon that was being selected for. pGM101 relies upon the pBR322 origin of replication (Balbás, et al. 1986, McVicker and Tang 2016) and is hence theoretically compatible with IncA/C plasmids, so we assumed the former to be most likely. To reduce tellurite stress, we re-transformed the pGM101 variants into PS_Koxy4 at a lower tellurite concentration of 2.5 μg/ml, as this seemed to be the minimum inhibitory concentration under our growth conditions (**Supplementary Figure 3**), and verified by PCR that the stocked transformants retained both pPSKoxy4_2 and the appropriate vector.

### PSKoxy_4 HipBA is a functional plasmid maintenance system

The HipB-encoding derivative of pGM101 did not decrease pPSKoxy4_2 plasmid stability relative to the empty vector control under 2.5 μg/ml tellurite selection, though this was likely due to the poor detection limit offered by colony PCR (≥1 % plasmid loss; data not shown). We therefore decided to employ an alternative method of plasmid stability testing. Plasmids encoding appropriate replication machinery and a counter-selectable *sacB-neoR* cassette (“pSTAB” vectors) can be used to test stability functions of plasmid maintenance elements, with plasmid loss detection limit as low as approximately 0.0001 % of a bacterial population (McVicker, et al. 2019, Martyn, et al. 2022). pSTAB vectors also have the advantage of isolating a given stability element from any other stability, resistance and virulence factors on the original host plasmid, therefore elucidating the role of that element alone.

We constructed a pSTAB vector combining the IncA/C replication origin from pPSKoxy4_2 with the *sacB-neoR* cassette from pIB279 (Blomfield, et al. 1991), with or without the *hipBA* locus from pPSKoxy4_2 (**Figure 4A**). These plasmids were designated pSTAB_incAC_ and pSTAB_incAC_::*hipBA*,respectively, and were transformed independently into either *E. coli* NEB 5-alpha or *K. oxytoca* NCTC 13727^T^ to observe plasmid stability in both *E. coli* and *Klebsiella* lacking native plasmids and any conflicting antibiotic resistance (**Figure 4B**). Growth of bacteria containing pSTAB_incAC_ for approximately 20-25 generations at 37 °C on LA resulted in a mean plasmid loss of approximately 1-10 % and a non-significant difference between the two host species (two-way ANOVA with Sidak’s multiple comparisons tests, *P* = 0.9712). The plasmid encoding HipBA was dramatically and significantly more stable than the empty vector in both species (*P* < 0.0001), with the TA system having a statistically greater effect in *K. oxytoca* than in *E. coli* (*P* < 0.0001), reaching a mean plasmid loss of less than 0.0001 %.

**Figure 4:**
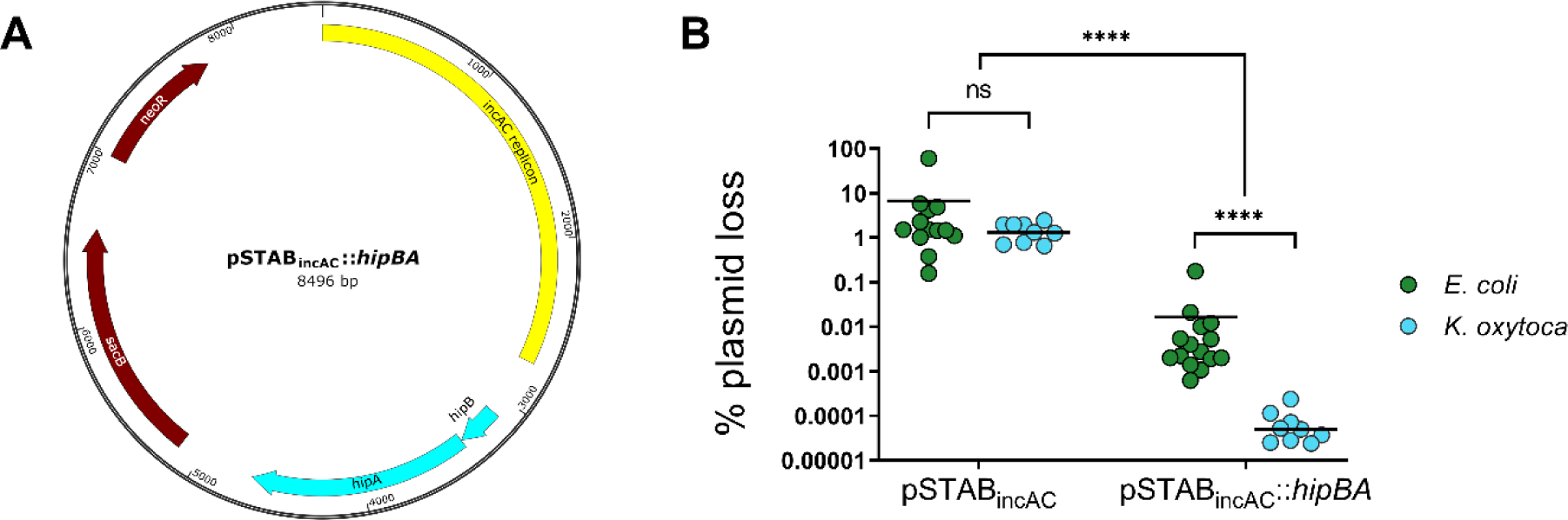
Impact of HipBA on stability of test vector pSTAB_incAC_. A: Map of pSTAB_incAC_::*hipBA*. Yellow: replication region from pPSKoxy4_2; red: selection markers; cyan: *hipBA*. **B: Plasmid loss of pSTAB_incAC_ constructs in *E. coli* (green) and *K. oxytoca* (blue).** Plasmid loss determined by plating onto neomycin (plasmid^+^) or sucrose (plasmid^−^) in accordance with previous work (McVicker, et al. 2019). Each point shows plasmid loss as a percentage of the total viable cell population within a single biological replicate (one independent colony grown on LA for 20-25 generations at 37°C). Solid lines show the mean. Data combined from at least two transformant clones per strain/plasmid combination and multiple independent experiments. Statistical analysis by two-way ANOVA with Sidak’s post test: ns, not significant; **** *P* < 0.0001.

### In silico analyses of HipBA in members of the KoSC

The identification of a functional plasmid-encoded HipBA in *K. michiganensis* PS_Koxy4 led us to investigate the prevalence of chromosomal- and plasmid-encoded versions of this TA system in the KoSC. A collection (*n* = 861) of high-quality KoSC genomes were included in this study: *K. michiganensis*, *n* = 393 (including PS_Koxy4); *K. oxytoca*, *n* = 235; *K. grimontii*, *n* = 185; *K. pasteurii*, *n* = 39; *Klebsiella* taxon 3, *n* = 4; *K. huaxiensis*, *n* = 3; *Klebsiella* taxon 1, *n* = 1; *Klebsiella* taxon 2, *n* = 1 (full details in **Supplementary Table 3**). The collection represented 234 different sequence types of the KoSC, with 104 genomes not assigned to known sequence types. Among the KoSC genomes, 761/861 were predicted to encode plasmids using the Kraken2-based detection method of Gomi et al. (2021) (**Supplementary Table 3**).

A SLING-based analysis predicted 292 chromosomal copies of HipA across 288 genomes: 236 genomes were predicted to encode chromosomal HipBA (two *K. oxytoca* genomes encoded two copies each), while 54 genomes encoded what appeared to be a chromosomal tripartite HipBAB system (**Figure 4**). Except for two *K. grimontii* ST559 genomes, encoding both HipBA and HipBAB on their chromosomes, the tripartite system was confined to 52 *K. michiganensis* genomes representing sequence types 85 (*n* = 23/23), 213 (*n* = 10/10), 11 (*n* = 9/9), 324 (*n* = 4/4) and 28 (*n* = 2/2), and unknown sequence types (n = 4/41) (**Supplementary Table 3**, **Supplementary Table 4**). Nine different variants of HipBAB were represented in the dataset: one in *K. grimontii* and eight in *K. michiganensis*. The *K. grimontii* and *K. michiganensis* variants shared <20 % amino-acid identity, while the *K. michiganensis* variants shared 97.69-99.83 % amino-acid identity. Detailed functional and structural analyses of the *K. grimontii* and *K. michiganensis* HipBAB variants would be required to determine whether they represent TA systems akin to the HipBST TA system of *E. coli* O127:H6 (Bærentsen, et al. 2023); however, these would have to be the subject of complex studies beyond the scope of the current report.

Chromosome-encoded HipBA was rare among *K. michiganensis* genomes (*n*=3/393, 0.76 %), instead found predominantly in *K. grimontii* (*n*=149/185, 80.54 %), *K. oxytoca* (*n*=83/235, 35.32 %) and *Klebsiella* taxon 3 (*n*=1/4, 25.00 %) (**Figure 5**).

**Figure 5:**
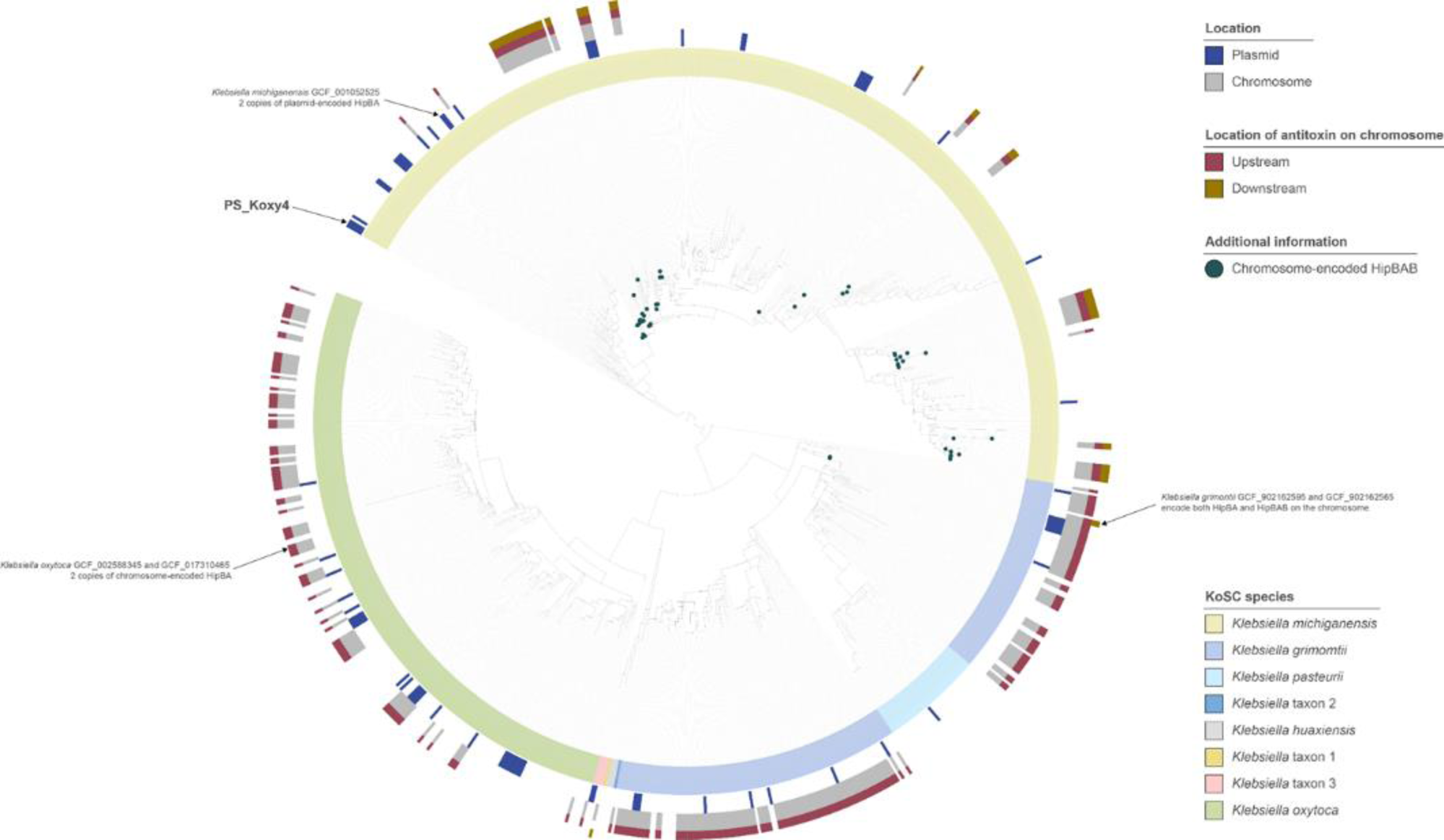
PhyloPhlAn-based analysis of proteome data for the 871 KoSC genomes included in this study. The tree (rooted at the mid-point) was created using RAxML implemented within PhyloPhlAn3 from MAFFT-generated multiple-sequence alignments, and visualized using iToL. The tree has been annotated to highlight genomes encoding chromosome- and/or plasmid-encoded variants of HipBA. Genomes encoding the antitoxin HipB both up- and down-stream of the toxin HipA are predicted to encode tripartite HipBAB (shown by green dots at tree tips).

All HipB-encoding (antitoxin) genes were predicted to be encoded upstream of the HipA-encoding (toxin) genes. However, there was sequence diversity among the plasmid-encoded HipBA pairs, with five distinct clusters observed in our phylogenetic analyses (**Figure 5**). Across the five clusters, with the exceptions of Cluster 1 and Cluster 2 sequences (83.94-84.62 % identity), plasmid-encoded HipA sequences shared low sequence identity (**Supplementary Table 5**). The same phenomenon was observed across the plasmid-encoded HipB sequences (Cluster 1 and Cluster 2, 71 % identity), with HipB sequences tending to share lower sequence identity than the HipA sequences (**Supplementary Table 6**). HipBA sequences associated with Clusters 1 and 2 shared highest identity (HipB – 35.82-36.57 % and 38.06 %, respectively; HipA – 57.53-57.75 % and 56.63 %, respectively) with HipBA (*hipBA*/Xre-HipA) of *Burkholderia pseudomallei* K96243. HipA sequences of Cluster 3, with which *K. michiganensis* PS_Koxy4 was affiliated, shared highest identity (53.96-67.82 %) with HipA (*hipBA*/−) of *Xenorhabdus nematophila* ATCC 19061. Except for one truncated Cluster 3 *K. michiganensis* (genome accession GCF_014050535) HipA sequence sharing 75.66 % identity with cluster members, all Cluster 3 HipA sequences were identical, as were their HipB sequences. Cluster 4 HipBA sequences shared highest identity (HipB – 21.49-22.31 %; HipA – 28.19-28.63 %) with HipBA (TA010076; *hipBA*/−) of *Caulobacter crescentus* NA1000. The singleton sequence associated with Cluster 5 shared 7.17-20.22 % identity with all HipB reference sequences and those of Clusters 1-4, while its HipA sequence shared 15.01-20.72 % identity with its reference and Clusters 1-4 counterparts.

Structurally, the HipBA system of PS_Koxy4 was almost identical to that of the experimentally validated HipBA system of *Xenorhabdus nematophila* ATCC 19061 (Yadav and Rathore 2020), with a sequence alignment score (matchmaker) of 1709.9 between the two HipA sequences (**Supplementary Figure 4**). The root mean square deviation between 421 pruned atom pairs was 0.488 Å, and across all 432 pairs was 0.816 Å.

**Figure 6:**
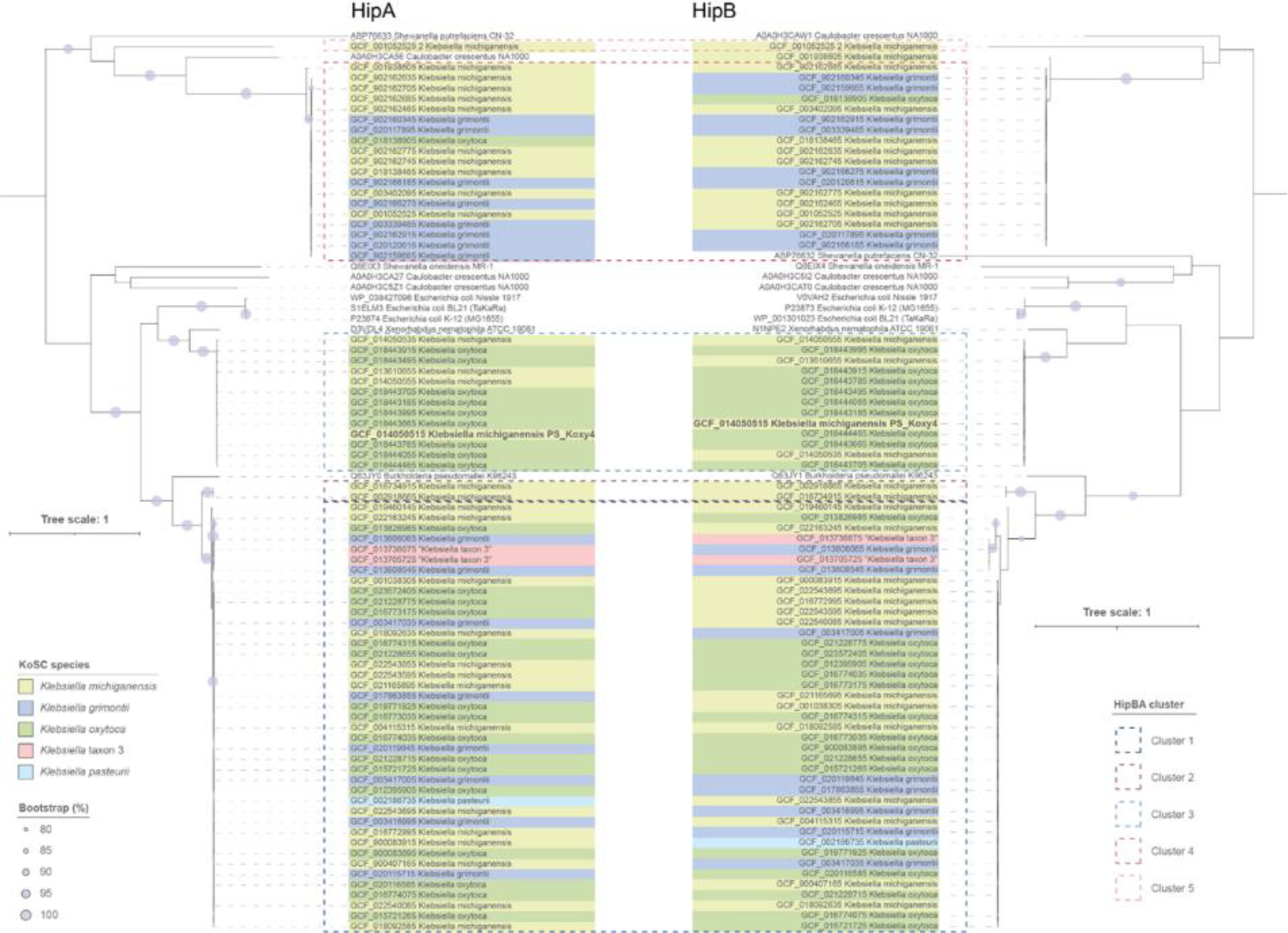
Phylogenetic relationships among plasmid-encoded HipA and HipB sequences of the KoSC. The trees (both rooted at the mid-point) were created using RAxML from MAFFT-generated multiple-sequence alignments, and visualized using iToL. Bootstrap values are presented as a percentage of 100 replications. Details of reference sequences included in the analyses can be found in **Table 2**.

## Discussion

TA systems of *Klebsiella* species have received little attention, with only four type II TA systems found in *K. pneumoniae* and two in *K. michiganensis* validated experimentally to date (**Table 3**). Using *in silico* analyses, Guan et al. (2024) recently identified 211697 TA systems (not dereplicated) in complete genomes representing 5980 different species of bacteria. Over one quarter (*n* = 59175, 28.0 %) of these TA systems were found in *E. coli*. Although TA systems predicted to be encoded by *K. pneumoniae* were second most abundant (*n* = 22173, 10.5 %) in the analyses undertaken by Guan and colleagues, other *Klebsiella* spp. were poorly represented in the dataset (**Supplementary Table 7**). Among the 52 complete *K. michiganensis* genomes included in their study, Guan et al. (2024) predicted 531 TA systems, including the CcdB, VapC, ParE and HipA toxins of PS_Koxy4 that we have characterized. We had previously identified these TA systems in PS_Koxy4 through manually checking Bakta-generated annotations and SLING-based analysis. Notably, we surmised that the ParE protein encoded by PS_Koxy4 would likely be inactive due to the gene’s truncation; a prediction that was borne out experimentally. This highlights an important and underappreciated discrepancy that can exist between *in silico* TA predictions and their biological activity.

**Table 3:**
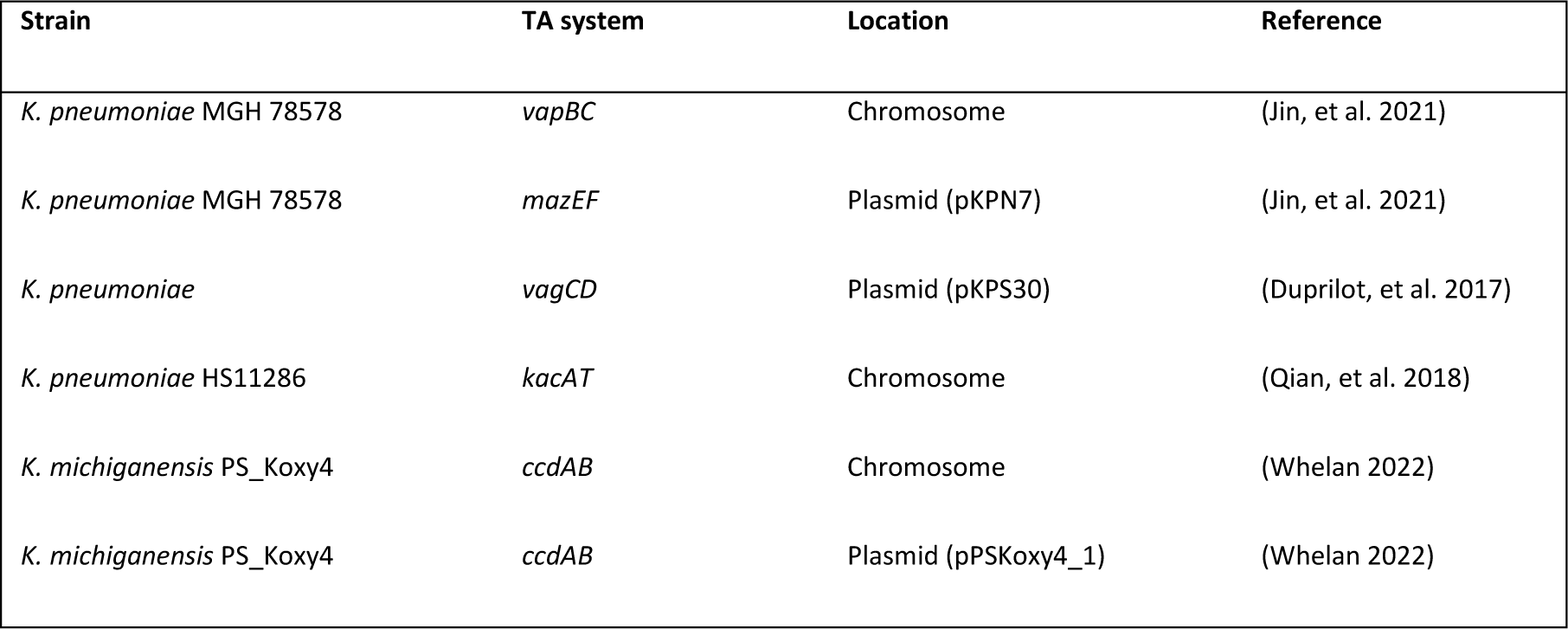
Details of experimentally validated *Klebsiella* type II TA systems.

The *hipBA* TA module is found in the chromosome of enteric bacteria such as *E. coli* and is widely understood to contribute towards persister formation and antimicrobial tolerance, whether or not the HipA protein is actively toxic to the host bacterium (Falla and Chopra 1998, Korch, Henderson and Hill 2003, Semanjski, et al. 2018). We have demonstrated that HipA from *K. michiganensis* pPSKoxy4_2 is highly toxic when expressed in both *E. coli* and *K. oxytoca* and that its associated antitoxin, HipB, is able to abrogate this toxicity as expected from a cognate TA pair. Commonly for type II TA systems, *hipBA* gene expression is autoregulated via conditional cooperativity, i.e. binding of the antitoxin molecule to the TA operon’s promoter (Black, Irwin and Moyed 1994). This is supported by our experiments, in which the native *hipBA* promoter drives sufficient *hipB* expression to counteract HipA production from induced pBAD33.

Non-toxic HipA mutants spontaneously generated by our experiments in *K. oxytoca* highlight not only the potency of this toxin variant in the KoSC (i.e. lethal enough to generate escape mutants prior to toxin induction), but also provide clues as to the role of specific amino acid residues within the protein. Individual experimental lineages resulted in different mutations. One such lineage possessed a frameshift mutation downstream of *hipA* codon 234, which unsurprisingly abolished toxin activity, most likely due to the removal of crucial kinase residues such as D308 (Huse and Kuriyan 2002, Correia, et al. 2006, Yadav and Rathore 2020) in addition to leaving only the first 54 % of the protein intact for folding and target binding. In other lineages, threonine-to-proline substitutions T175P and T327P were identified, the former of which we reconstructed *de novo* to verify lack of toxicity. Both of these threonine residues are conserved in the *E. coli* K-12 chromosomal HipA sequence but have not been identified as important for toxin function previously. That both mutations introduced similar structural changes to the toxin, based on our *in silico* analyses (**Supplementary Figure 2**), suggests they inactivate the toxin in the same way and that threonine residues at positions 175 and 327 are essential for toxicity of plasmid-encoded HipA in PS_Koxy4.

Genes encoding HipA and related antitoxins have been identified on MGEs from a wide range of species, including an *Acinetobacter* resistance plasmid harbouring the *tet*(X) and *bla*_OXA-58_ genes (Wang, et al. 2020) and multiple virulence plasmids in *Cronobacter sakazakii* (Finkelstein, et al. 2019). HipBA has even been proposed to contribute to *Rhizobium* symbiosis plasmid stability (Falla and Chopra 1999), though this was based solely upon its nature as a TA system and no experimental evidence was provided for the claim. Prior to this work, HipBA was shown to stabilize a chromosomal genomic island in *Shewanella putrefaciens* (Zhao, et al. 2022), but no extrachromosomal MGEs. We therefore present herein the first characterization of HipBA as a potent plasmid addiction system, showing that a pPSKoxy4_2 IncA/C-replicon vector can be stabilized approximately 10,000-fold in *K. oxytoca* by the inclusion of *hipBA*; notably, this is far more effective than many other characterized PSK systems, including those tested in *K. pneumoniae* (Wu, Kamruzzaman and Iredell 2020, Wilbaux, et al. 2007, McVicker, et al. 2019). The *hipBA* module also increases stability of the same plasmid in *E. coli*, but to a significantly lesser extent. Such reduced functionality may be due to the decreased toxicity of pPSKoxy4_2 HipA in the latter species, as evidenced by the generation of toxin-inactivating mutations in *K. oxytoca* but not in *E. coli*. If so, this may reflect a difference in the cellular target of the kinase, thought to be part of the bacterial translation machinery (Kaspy, et al. 2013, Schumacher, et al. 2009, Bokinsky, et al. 2013). Alternatively, it may represent a disparity in how efficiently the pPSKoxy4_2 replicon functions in the two species, though this seems less likely since pSTAB_incAC_ was lost to the same degree in both organisms when it lacked *hipBA*.

In summary, we have provided the first experimental evidence that a plasmid-borne HipBA TA system can provide a formidable plasmid-stabilising effect in both *E. coli* and *K. oxytoca*, most likely through the action of PSK. Furthermore, we have identified similar plasmid-borne *hipBA* loci across the entirety of the KoSC. This adds substantially both to our knowledge of multiple medically-relevant *Klebsiella* species and to the mechanistic understanding of an important and widespread TA system in enteric bacteria.

## Data availability

The data underlying this article will be shared on reasonable request to the corresponding author. Additional supplementary materials associated with this manuscript are available from figshare: https://figshare.com/projects/The_plasmid-borne_hipBA_operon_of_Klebsiella_michiganensis_encodes_a_potent_plasmid_stabilisation_system/213793.

## Author contributions

Shutt-McCabe – Investigation, Formal analysis, Writing – review & editing. Shaik – Investigation. Hoyles – Conceptualization, Data curation, Formal analysis, Investigation, Methodology, Visualization, Writing – original draft, Writing – review & editing. McVicker – Conceptualization, Data curation, Formal analysis, Funding acquisition, Investigation, Methodology, Project administration, Supervision, Visualization, Writing – original draft, Writing – review & editing.

## Abbreviations

ANI: average nucleotide identity
KoSC: *Klebsiella oxytoca* species complex
LA: lysogeny agar
LB: lysogeny broth
MGE: mobile genetic element
PSK: post-segregational killing
TA: toxin–antitoxin.

---

**Supplementary Figure 1:**
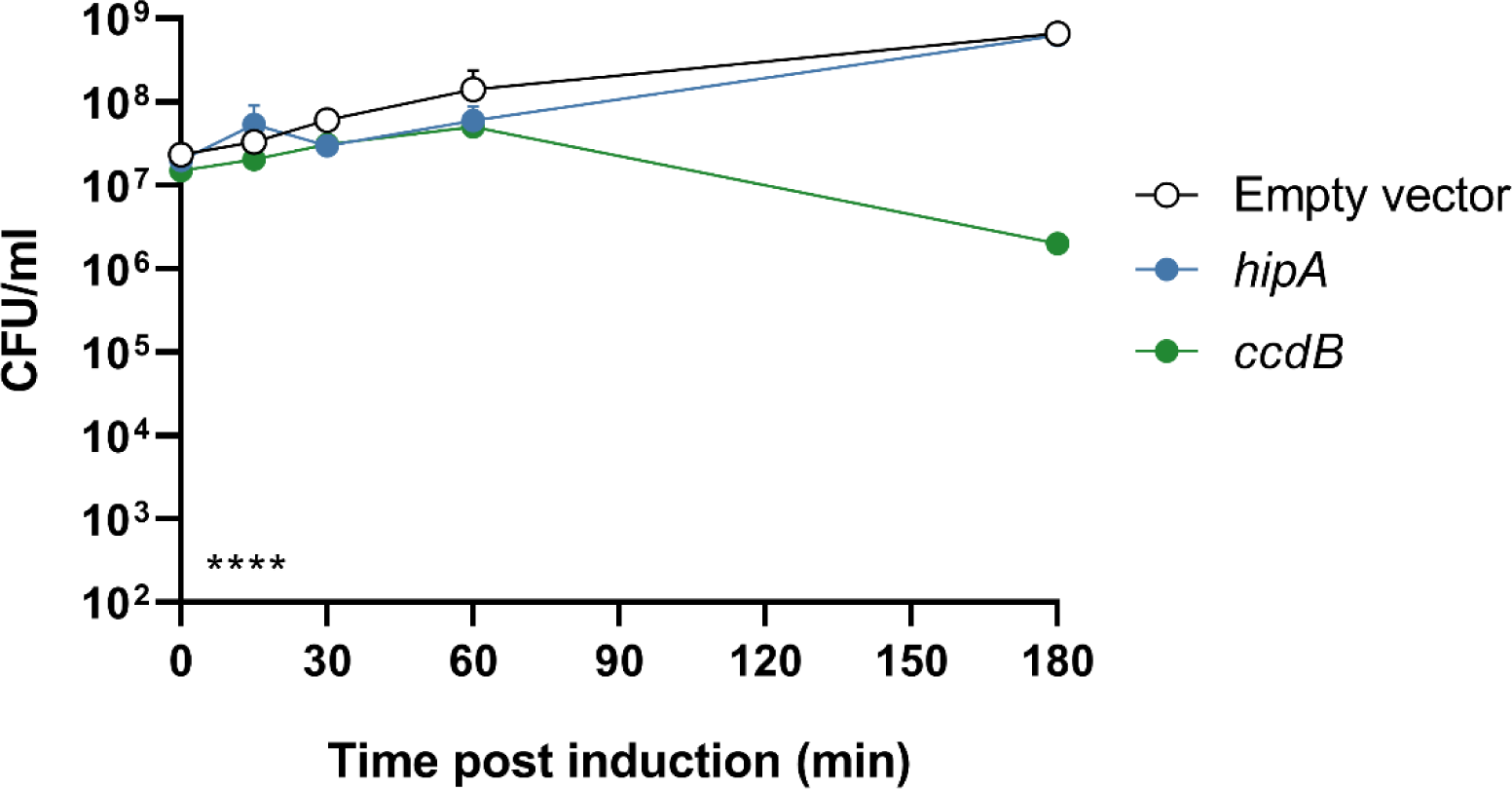
*K. oxytoca* survives induced *hipA* expression. Mean bacterial viability after induction of production of the indicated genes from pBAD33. Cells also contain empty pGM101. Gene expression induced in *K. oxytoca* NCTC 13727^T^. Bars show standard error of the mean. Statistical analysis by two-way ANOVA: indicator in lower left shows the overall effect of strain; **** *P* < 0.0001 (from *n* = 2 biological replicates). Multiplicity-adjusted statistical analysis from Tukey’s post tests given in the main text.

**Supplementary Figure 2:**
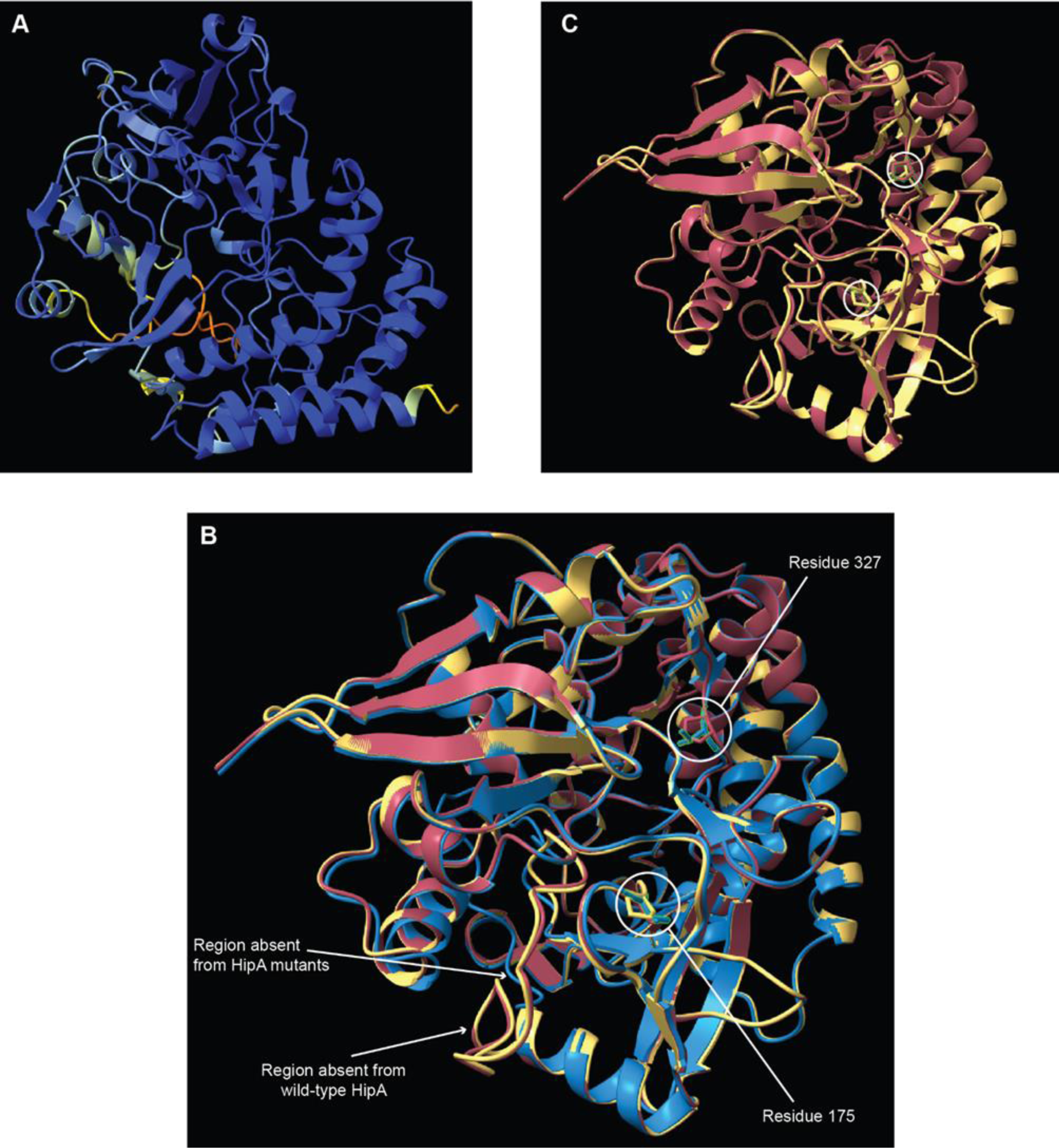
Structural comparisons of wild-type and mutated HipA. A: AlphaFold-predicted structure of wild-type HipA of PS_Koxy4. The more blue elements of the structure are, the more confident AlphaFold is in the predicted structure. **B: Alignment of the AlphaFold-predicted variant HipA structures - HipA T175P (yellow) and HipA T327P (red) - against the wild-type HipA (blue).** The residues that have been mutated are shown in green (also indicated by circles). The atoms (threonine and proline) are shown at the respective mutated residues. Although the T->P mutations are in different regions of the HipA molecule, both are predicted to introduce a similar structural change in the shape of HipA. **C: The mutated HipA structures are more similar to one another than they are to wild type HipA.**

**Supplementary Figure 3:**
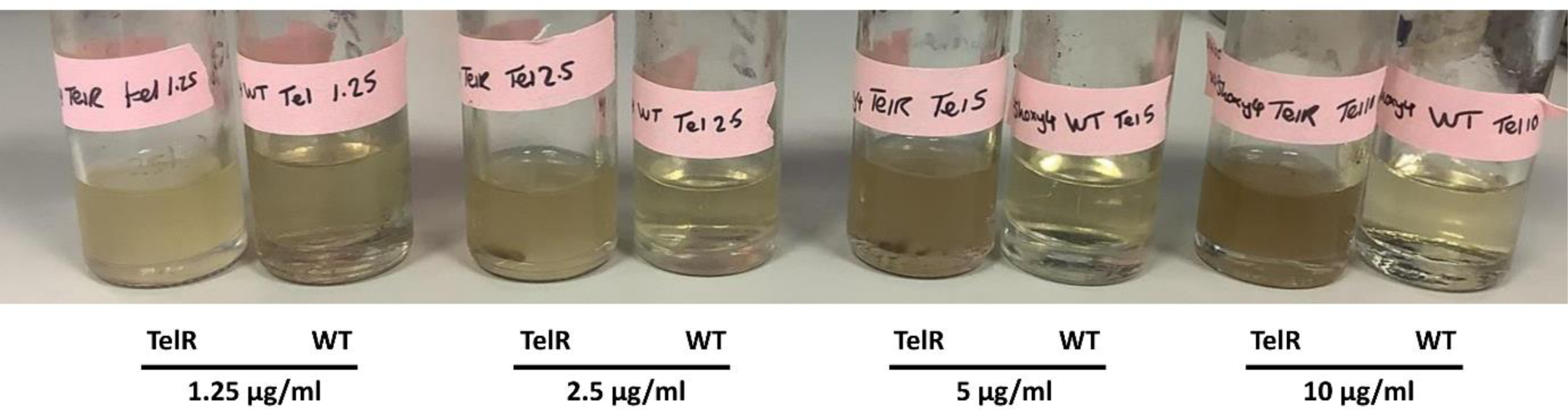
Survival of *K. michiganensis* PS_Koxy4 in tellurite. Overnight cultures grown in 5 ml LB with the indicated concentration of potassium tellurite. “TelR” cultures contain empty pGM101-TelR. “WT” cultures contain no vector.

**Supplementary Figure 4:**
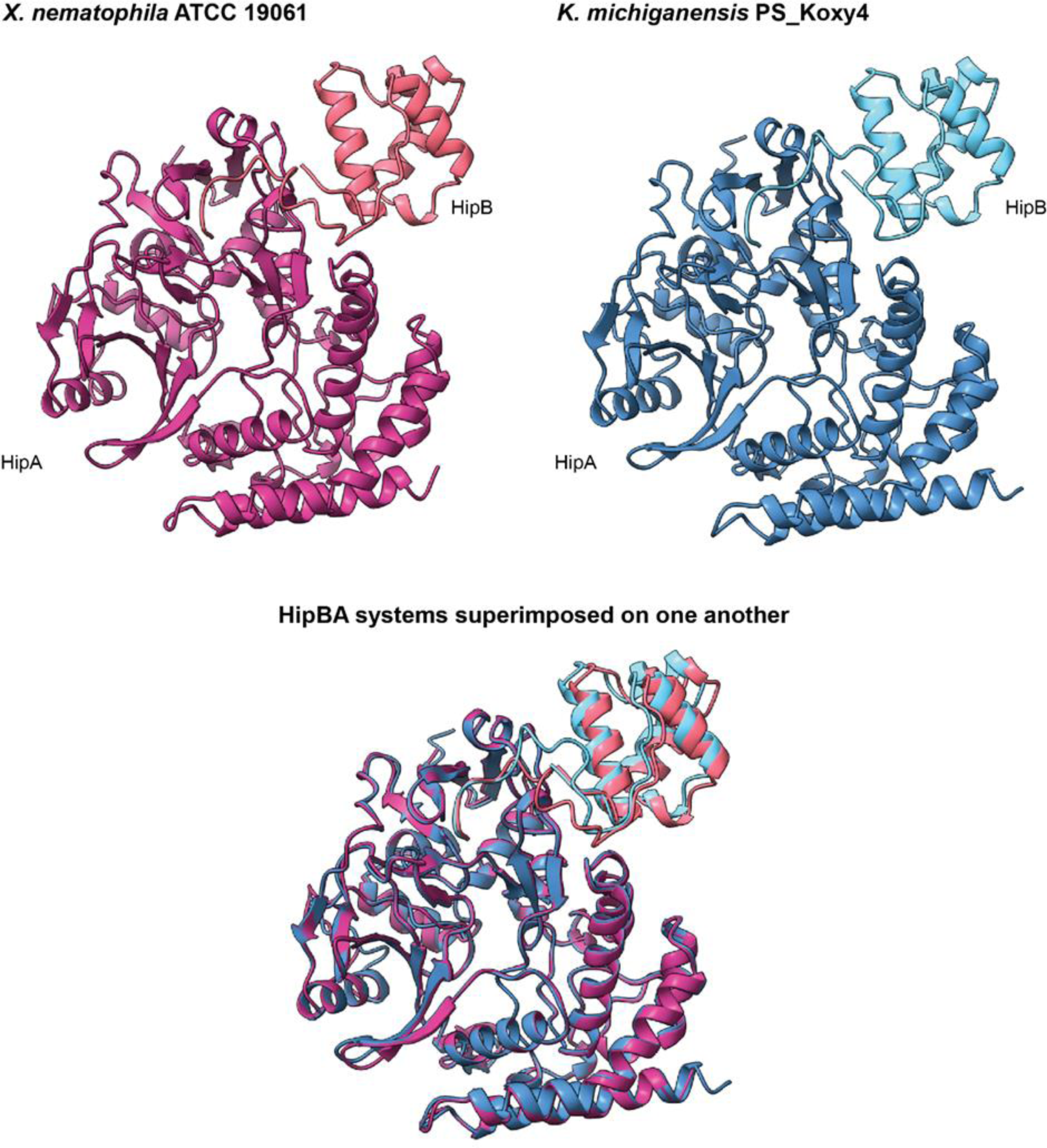
Structural similarity of the plasmid-encoded HipBA system of *K. michiganensis* PS_Koxy4 with its closest experimentally validated TA system (*X. nematophila* ATCC 19061). All structural predictions were made using AlphaFold-multimer, via ChimeraX v1.8; the matchmaker command of ChimeraX was used to align the structures to create the superimposed image. The analysis demonstrated the two HipBA TA systems share high structural similarity.

