## Supplementary Table 7 for "The plasmid-borne *hipBA* operon of *Klebsiella michiganensis* encodes a potent plasmid stabilization system"

**Supplementary Table 7. Summary of *Klebsiella* TA system information provided by Guan *et al.* (2024)**

| **Species** | **Number of:** | | |
| --- | --- | --- | --- |
|  | **Predicted loci** | **Experimentally validated TA systems** | **Genomes included in analyses** |
| *Klebsiella pneumoniae* | 22177 | 4 | 1562 |
| *Klebsiella variicola* | 983 | 0 | 75 |
| *Klebsiella quasipneumoniae* | 845 | 0 | 103 |
| *Klebsiella michiganensis* | 531 | 0 | 52 |
| *Klebsiella aerogenes* | 335 | 0 | 45 |
| *Klebsiella oxytoca* | 302 | 0 | 36 |
| *Klebsiella grimontii* | 144 | 0 | 14 |
