## Supplementary Table 1 for "The plasmid-borne *hipBA* operon of *Klebsiella michiganensis* encodes a potent plasmid stabilization system"

**Supplementary Table 1. Plasmids used in this study.**

| **Plasmid ID** | **Construction/purpose** | **Source** |
| --- | --- | --- |
| pBAD33 | Arabinose-inducible vector for toxicity assays | Guzman, et al. 1995 |
| pBAD33::*ccdB* | pBAD33 encoding CcdB (pPSKoxy4_1) | This study |
| pBAD33::*hipA* | pBAD33 encoding HipA (pPSKoxy4_2) | This study |
| pBAD33::*parE* | pBAD33 encoding ParE (pPSKoxy4_3) | This study |
| pBAD33::*vapC* | pBAD33 encoding VapC (pPSKoxy4_3) | This study |
| pIB279 | Source of *sacB*-NeoR cassette for pSTAB vectors | Blomfield, et al. 1991 |
| pGM101_neo_ | Promoterless pBAD33-compatible cloning vector | Whelan 2022 |
| pGM101_neo_::*ccdA* | pGM101_neo_ containing *ccdA* (pPSKoxy4_1) plus promoter | This study |
| pGM101_neo_::*hipB* | pGM101_neo_ containing *hipB* (pPSKoxy4_2) plus promoter | This study |
| pGM101_tel_ | Promoterless pBAD33-compatible cloning vector | This study |
| pGM101_tel_::*hipB* | pGM101_tel_ containing *hipB* (pPSKoxy4_2) plus promoter | This study |
| pMo130-TelR | Source of tellurite resistance (*kilA-telAB*) for pGM101_tel_ | Amin, et al. 2013 |
| pSTAB_incAC_ | pSTAB with IncA/C replicon (pPSKoxy4_2) | This study |
| pSTAB_incAC_::*hipBA* | pSTAB_incAC_ containing entire *hipBA* locus (pPSKoxy4_2) | This study |
