## Supplementary Table 2 for "The plasmid-borne *hipBA* operon of *Klebsiella michiganensis* encodes a potent plasmid stabilization system"

**Supplementary Table 2. Oligonucleotide primers used in this study.**

| **Primer ID** | **Sequence (5’-3’)** | **Target** |
| --- | --- | --- |
| JSMntu001 | GGATCCTCTAGAGTCGAC | pBAD33 |
| JSMntu002 | CCGGGTACCGAGCTCGAATTC | pBAD33 |
| JSMntu003 | aattcgagctcggtacccggGAAAACAGGGACTGGTGAG | *ccdB* |
| JSMntu004 | aggtcgactctagaggatccTCAAATTCCCCAGAACATC | *ccdB* |
| JSMntu005 | aattcgagctcggtacccggGGGATAGGGAGTGGTAAG | *hipA* |
| JSMntu006 | aggtcgactctagaggatccTCATTTGGCTACTCCCTTC | *hipA* |
| JSMntu007 | aattcgagctcggtacccggGAAGGACGCTTTGAGCCATG | *vapC* |
| JSMntu008 | aggtcgactctagaggatccTTATTTCACCCAGTCTTCCAGC | *vapC* |
| JSMntu011 | aattcgagctcggtacccggGTCAGGGCCAGGGTGGCA | *parE* |
| JSMntu012 | aggtcgactctagaggatccCTATGCATCCGCCTGAACAACCC | *parE* |
| JSMntu013 | CGAAGCGGCATGCATTTACG | pGM101_neo_ |
| JSMntu014 | CCTTCGCGCGCGAATTGATC | pGM101_neo_ |
| JSMntu015 | gatcaattcgcgcgcgaaggTGGTACACTTCCGGAAAC | *ccdA* |
| JSMntu016 | cgtaaatgcatgccgcttcgTCACCAGTCCCTGTTTTC | *ccdA* |
| JSMntu017 | gatcaattcgcgcgcgaaggGCAAGCTTCGGGGCGCTC | *hipB* |
| JSMntu018 | cgtaaatgcatgccgcttcgTTACCACTCCCTATCCCAGACTTTACTATCG | *hipB* |
| JSMntu023 | cgagataaccgttggcct | *hipA* |
| JSMntu024 | gttcggcaagaagaaggtg | *hipA* |
| JSMntu025 | gaagatggagaagttcttcc | *hipA* |
| JSMntu026 | gcagtgcagttttctcttgg | *hipA* |
| JSMntu027 | cttgttgggggGCTGCCCAGGGGGCGTTG | *hipA* (mutant) |
| JSMntu028 | ctgggcagccCCCCAACAAGCCATATTTTTAAG | *hipA* (mutant) |
| JSMntu029 | CTGTCAGACCAAGTTTACTC | pGM101_neo_ |
| JSMntu030 | AGAAACGCAAAAAGGCCATC | pGM101_neo_ |
| JSMntu031 | gatggcctttttgcgtttctGATTAATGGTCAACAGCTCAAGC | pMo130-TelR |
| JSMntu032 | gagtaaacttggtctgacagCTGATATCAGGGCCCCGC | pMo130-TelR |
| JSMntu035 | gaaaggcagagtaagggtag | pPSKoxy4_2 IncAC |
| JSMntu036 | gtgcagcatccccatatatt | pPSKoxy4_2 IncAC |
| JSMntu037 | taccgagataacaaaggggc | *hipA* |
| JSMntu038 | ctttgcaatgcgcaaacaga | *hipA* |
| GMntu084 | gaggcttgcaacaacatctgtagcagctac | pPSKoxy4_2 IncAC |
| GMntu085 | aaaaaggatctctagagctgacatggttc | pPSKoxy4_2 IncAC |
| GMntu086 | cagctctagagatcctttttaacccatcacatatacctgc | pIB279 |
| GMntu087 | cagatgttgttgcaagcctcgtcgtcctg | pIB279 |
| GMntu088 | gaggcttgcaacaacatctgtagcagctac | pPSKoxy4_2 IncAC |
| GMntu089 | tgcggataattctagagctgacatggttc | pPSKoxy4_2 IncAC |
| GMntu090 | cagctctagaattatccgcacatgccttaatc | *hipBA* |
| GMntu091 | aaaaaggatctcatttggctactcccttc | *hipBA* |
| GMntu092 | agccaaatgagatcctttttaacccatcacatatacctgc | pIB279 |
| GMntu093 | cagatgttgttgcaagcctcgtcgtcctg | pIB279 |
| MS083 | aattcgagctcggtacccggagtaaagtctgggatagg | *hipA* |
