## Supplementary Table 3 for "The plasmid-borne *hipBA* operon of *Klebsiella michiganensis* encodes a potent plasmid stabilization system"

**Supplementary Table 3. Seq genomes included in this study**

| **Assembly** | **Completeness** | **Contamination** | **ANI reference genome** | **Plasmid(s) detected?** | **Plasmid HipBA** | **ST** | ***gapA*** | ***infB*** | ***mdh*** | ***pgi*** | ***phoE*** | ***rpoB*** | ***tonB*** |
| --- | --- | --- | --- | --- | --- | --- | --- | --- | --- | --- | --- | --- | --- |
| GCF_003201885.1 | 100 | 3 | "*Klebsiella* taxon 1" 100 % | Yes | No | - | ~31 | - | - | ~81 | - | 28? | - |
| GCF_009707385.1 | 100 | 0.16 | "*Klebsiella* taxon 2" 100 % | No | No | - | ~40 | ~25 | - | - | - | ~55 | - |
| GCF_013705725.1 | 100 | 3.04 | "*Klebsiella* taxon 3" 100 % | Yes | Yes | - | ~31 | - | - | ~81 | - | 24? | - |
| GCF_014103785.1 | 100 | 2.36 | "*Klebsiella* taxon 3" 96.8993 % | Yes | No | - | ~31 | - | - | ~81 | - | 24? | - |
| GCF_014103505.1 | 100 | 2.31 | "*Klebsiella* taxon 3" 99.5736 % | Yes | No | - | ~31 | - | - | ~81 | - | 24? | - |
| GCF_013736875.1 | 100 | 3.04 | "*Klebsiella* taxon 3" 99.9966 % | Yes | Yes | - | ~31 | - | - | ~81 | - | 24? | - |
| GCF_900200035.1 | 100 | 0.93 | *Klebsiella grimontii* 100 % | Yes | No | 319 | 4 | 6 | 14 | 11 | 46 | 11 | 9 |
| GCF_020116755.1 | 100 | 0.55 | *Klebsiella grimontii* 97.022 % | Yes | No | - | 17 | 35 | 85 | ~31 | ~118 | 55 | ~107 |
| GCF_020120615.1 | 100 | 0.25 | *Klebsiella grimontii* 98.8944 % | Yes | Yes | 408 | 21 | 6 | 19 | 10 | 46 | 7 | 9 |
| GCF_001065765.1 | 100 | 0.97 | *Klebsiella grimontii* 98.9063 % | Yes | No | 563 | 5 | 6 | 18 | 10 | 46 | 8 | 31 |
| GCF_902159715.1 | 100 | 0.23 | *Klebsiella grimontii* 98.9171 % | Yes | No | 216 | 4 | 6 | 19 | 10 | 46 | 24 | 31 |
| GCF_902162915.1 | 100 | 0.48 | *Klebsiella grimontii* 98.9442 % | Yes | Yes | 263 | 6 | 6 | 26 | 10 | 46 | 24 | 33 |
| GCF_016656825.1 | 100 | 1.25 | *Klebsiella grimontii* 98.9501 % | Yes | No | 350 | 4 | 6 | 11 | 10 | 46 | 8 | 9 |
| GCF_013707575.1 | 100 | 0.62 | *Klebsiella grimontii* 98.9952 % | Yes | No | 184 | 4 | 6 | 26 | 10 | 46 | 5 | 6 |
| GCF_902166185.1 | 100 | 0.46 | *Klebsiella grimontii* 98.9964 % | Yes | Yes | 263 | 6 | 6 | 26 | 10 | 46 | 24 | 33 |
| GCF_902160275.1 | 100 | 1.34 | *Klebsiella grimontii* 98.9992 % | Yes | No | 577 | 15 | 6 | 19 | 11 | 46 | 5 | 31 |
| GCF_902166275.1 | 100 | 0.48 | *Klebsiella grimontii* 99.0034 % | Yes | Yes | 263 | 6 | 6 | 26 | 10 | 46 | 24 | 33 |
| GCF_902162565.1 | 100 | 0.24 | *Klebsiella grimontii* 99.0042 % | Yes | No | 559 | 5 | 6 | 2 | 10 | 46 | 5 | 63 |
| GCF_015208375.1 | 100 | 1.92 | *Klebsiella grimontii* 99.0046 % | Yes | No | 350 | 4 | 6 | 11 | 10 | 46 | 8 | 9 |
| GCF_902162595.1 | 100 | 0.24 | *Klebsiella grimontii* 99.0063 % | Yes | No | 559 | 5 | 6 | 2 | 10 | 46 | 5 | 63 |
| GCF_902165695.1 | 100 | 0.47 | *Klebsiella grimontii* 99.0088 % | Yes | No | 566 | 5 | 6 | 26 | 10 | 46 | 7 | 9 |
| GCF_902159625.1 | 100 | 0.74 | *Klebsiella grimontii* 99.0189 % | Yes | No | 186 | 21 | 6 | 14 | 10 | 62 | 7 | 9 |
| GCF_014103875.1 | 100 | 0.61 | *Klebsiella grimontii* 99.0213 % | Yes | No | 263 | 6 | 6 | 26 | 10 | 46 | 24 | 33 |
| GCF_902160335.1 | 100 | 1.28 | *Klebsiella grimontii* 99.0241 % | Yes | No | 577 | 15 | 6 | 19 | 11 | 46 | 5 | 31 |
| GCF_902160225.1 | 100 | 0.63 | *Klebsiella grimontii* 99.0266 % | Yes | No | 361 | 6 | 6 | 14 | 10 | 46 | 8 | 33 |
| GCF_902160605.1 | 100 | 0.43 | *Klebsiella grimontii* 99.0387 % | Yes | No | 517 | 4 | 6 | 26 | 10 | 46 | 7 | 81 |
| GCF_902160355.1 | 100 | 1.26 | *Klebsiella grimontii* 99.0536 % | Yes | No | 577 | 15 | 6 | 19 | 11 | 46 | 5 | 31 |
| GCF_902160265.1 | 100 | 1.27 | *Klebsiella grimontii* 99.0543 % | Yes | No | 577 | 15 | 6 | 19 | 11 | 46 | 5 | 31 |
| GCF_902160345.1 | 100 | 0.24 | *Klebsiella grimontii* 99.0544 % | Yes | Yes | 168 | 15 | 6 | 39 | 10 | 46 | 24 | 6 |
| GCF_902162835.1 | 100 | 0.82 | *Klebsiella grimontii* 99.0548 % | Yes | No | 411 | 5 | 6 | 19 | 10 | 109 | 5 | 6 |
| GCF_902160515.1 | 100 | 0.45 | *Klebsiella grimontii* 99.0581 % | Yes | No | 186 | 21 | 6 | 14 | 10 | 62 | 7 | 9 |
| GCF_902164155.1 | 100 | 0.88 | *Klebsiella grimontii* 99.0586 % | Yes | No | 246 | 5 | 10 | 59 | 10 | 46 | 7 | 9 |
| GCF_902160245.1 | 100 | 0.64 | *Klebsiella grimontii* 99.0592 % | Yes | No | 361 | 6 | 6 | 14 | 10 | 46 | 8 | 33 |
| GCF_013606065.1 | 100 | 0.46 | *Klebsiella grimontii* 99.0601 % | Yes | Yes | 225 | 4 | 6 | 26 | 10 | 80 | 7 | 63 |
| GCF_902160365.1 | 100 | 1.75 | *Klebsiella grimontii* 99.0713 % | Yes | No | 577 | 15 | 6 | 19 | 11 | 46 | 5 | 31 |
| GCF_902164205.1 | 100 | 0.8 | *Klebsiella grimontii* 99.0773 % | Yes | No | 246 | 5 | 10 | 59 | 10 | 46 | 7 | 9 |
| GCF_902160405.1 | 100 | 3.31 | *Klebsiella grimontii* 99.0822 % | Yes | No | 577 | 15 | 6 | 19 | 11 | 46 | 5 | 31 |
| GCF_001052825.1 | 100 | 0.24 | *Klebsiella grimontii* 99.0848 % | Yes | No | 408 | 21 | 6 | 19 | 10 | 46 | 7 | 9 |
| GCF_902160585.1 | 100 | 0.52 | *Klebsiella grimontii* 99.086 % | Yes | No | 517 | 4 | 6 | 26 | 10 | 46 | 7 | 81 |
| GCF_013734595.1 | 100 | 1.59 | *Klebsiella grimontii* 99.0867 % | Yes | No | 350 | 4 | 6 | 11 | 10 | 46 | 8 | 9 |
| GCF_902160175.1 | 100 | 0.63 | *Klebsiella grimontii* 99.0938 % | Yes | No | 361 | 6 | 6 | 14 | 10 | 46 | 8 | 33 |
| GCF_902160195.1 | 100 | 0.62 | *Klebsiella grimontii* 99.0961 % | Yes | No | 361 | 6 | 6 | 14 | 10 | 46 | 8 | 33 |
| GCF_013624225.1 | 100 | 0.16 | *Klebsiella grimontii* 99.1008 % | Yes | No | 263 | 6 | 6 | 26 | 10 | 46 | 24 | 33 |
| GCF_902160375.1 | 100 | 3.1 | *Klebsiella grimontii* 99.1011 % | Yes | No | 577 | 15 | 6 | 19 | 11 | 46 | 5 | 31 |
| GCF_902160675.1 | 100 | 2.19 | *Klebsiella grimontii* 99.1064 % | Yes | No | - | 5 | 10 | 59 | 10 | 46 | 7 | ~100 |
| GCF_014104195.1 | 100 | 0.66 | *Klebsiella grimontii* 99.1136 % | Yes | No | 263 | 6 | 6 | 26 | 10 | 46 | 24 | 33 |
| GCF_022540165.1 | 100 | 0.6 | *Klebsiella grimontii* 99.1138 % | Yes | No | 371 | 15 | 6 | 19 | 10 | 46 | 24 | 6 |
| GCF_020889225.1 | 100 | 1.13 | *Klebsiella grimontii* 99.1164 % | Yes | No | 215 | 21 | 6 | 14 | 10 | 46 | 11 | 6 |
| GCF_013730235.1 | 100 | 0.33 | *Klebsiella grimontii* 99.1174 % | Yes | No | 411 | 5 | 6 | 19 | 10 | 109 | 5 | 6 |
| GCF_002080105.1 | 100 | 0.58 | *Klebsiella grimontii* 99.1189 % | Yes | No | 236 | 27 | 6 | 14 | 11 | 46 | 5 | 9 |
| GCF_902161065.1 | 100 | 0.6 | *Klebsiella grimontii* 99.1189 % | Yes | No | 411 | 5 | 6 | 19 | 10 | 109 | 5 | 6 |
| GCF_902159595.1 | 100 | 0.68 | *Klebsiella grimontii* 99.1239 % | Yes | No | 517 | 4 | 6 | 26 | 10 | 46 | 7 | 81 |
| GCF_022559855.1 | 100 | 0.73 | *Klebsiella grimontii* 99.1252 % | Yes | No | 391 | 5 | 54 | 14 | 11 | 46 | 5 | 6 |
| GCF_014901955.1 | 100 | 3.29 | *Klebsiella grimontii* 99.126 % | Yes | No | - | 4 | 6 | 39 | 10 | 46 | 5 | 81 |
| GCF_001052235.1 | 100 | 0.65 | *Klebsiella grimontii* 99.1266 % | Yes | No | 186 | 21 | 6 | 14 | 10 | 62 | 7 | 9 |
| GCF_902161195.1 | 100 | 0.6 | *Klebsiella grimontii* 99.1268 % | Yes | No | 411 | 5 | 6 | 19 | 10 | 109 | 5 | 6 |
| GCF_017863855.1 | 100 | 0.48 | *Klebsiella grimontii* 99.128 % | Yes | Yes | 172 | 6 | 6 | 19 | 11 | 46 | 7 | 9 |
| GCF_902165435.1 | 100 | 1.15 | *Klebsiella grimontii* 99.1289 % | Yes | No | 556 | 4 | 6 | 19 | 10 | 46 | 24 | 32 |
| GCF_013608545.1 | 100 | 0.72 | *Klebsiella grimontii* 99.1342 % | Yes | Yes | - | 5 | 6 | 19 | ~10 | 46 | 11 | ~113 |
| GCF_001072835.1 | 100 | 1.5 | *Klebsiella grimontii* 99.1366 % | Yes | No | 215 | 21 | 6 | 14 | 10 | 46 | 11 | 6 |
| GCF_902161205.1 | 100 | 0.51 | *Klebsiella grimontii* 99.1376 % | Yes | No | 411 | 5 | 6 | 19 | 10 | 109 | 5 | 6 |
| GCF_902165565.1 | 100 | 0.67 | *Klebsiella grimontii* 99.139 % | Yes | No | 556 | 4 | 6 | 19 | 10 | 46 | 24 | 32 |
| GCF_003261555.1 | 100 | 0.73 | *Klebsiella grimontii* 99.1392 % | Yes | No | 215 | 21 | 6 | 14 | 10 | 46 | 11 | 6 |
| GCF_902162695.1 | 100 | 0.65 | *Klebsiella grimontii* 99.1392 % | Yes | No | 556 | 4 | 6 | 19 | 10 | 46 | 24 | 32 |
| GCF_902166145.1 | 100 | 0.84 | *Klebsiella grimontii* 99.1405 % | Yes | No | 556 | 4 | 6 | 19 | 10 | 46 | 24 | 32 |
| GCF_902159485.1 | 100 | 0.68 | *Klebsiella grimontii* 99.1421 % | Yes | No | 517 | 4 | 6 | 26 | 10 | 46 | 7 | 81 |
| GCF_902165535.1 | 100 | 1.22 | *Klebsiella grimontii* 99.1449 % | Yes | No | 408 | 21 | 6 | 19 | 10 | 46 | 7 | 9 |
| GCF_902162375.1 | 100 | 0.74 | *Klebsiella grimontii* 99.1513 % | Yes | No | 186 | 21 | 6 | 14 | 10 | 62 | 7 | 9 |
| GCF_017863935.1 | 100 | 0.59 | *Klebsiella grimontii* 99.1535 % | Yes | No | 189 | 5 | 10 | 10 | 10 | 70 | 5 | 9 |
| GCF_902161245.1 | 100 | 0.08 | *Klebsiella grimontii* 99.1566 % | Yes | No | 408 | 21 | 6 | 19 | 10 | 46 | 7 | 9 |
| GCF_020118395.1 | 100 | 0.33 | *Klebsiella grimontii* 99.1587 % | Yes | No | 431 | 21 | 6 | 19 | 10 | 46 | 7 | 6 |
| GCF_001076805.1 | 100 | 0.63 | *Klebsiella grimontii* 99.1592 % | Yes | No | 215 | 21 | 6 | 14 | 10 | 46 | 11 | 6 |
| GCF_902159665.1 | 100 | 0.21 | *Klebsiella grimontii* 99.1602 % | Yes | Yes | 327 | 4 | 6 | 68 | 11 | 46 | 5 | 79 |
| GCF_902161625.1 | 100 | 0.25 | *Klebsiella grimontii* 99.161 % | Yes | No | 225 | 4 | 6 | 26 | 10 | 80 | 7 | 63 |
| GCF_013744615.1 | 100 | 0.16 | *Klebsiella grimontii* 99.1655 % | Yes | No | 263 | 6 | 6 | 26 | 10 | 46 | 24 | 33 |
| GCF_902160215.1 | 100 | 0.15 | *Klebsiella grimontii* 99.1671 % | Yes | No | 184 | 4 | 6 | 26 | 10 | 46 | 5 | 6 |
| GCF_001070955.1 | 100 | 1.19 | *Klebsiella grimontii* 99.1673 % | Yes | No | 215 | 21 | 6 | 14 | 10 | 46 | 11 | 6 |
| GCF_002856195.1 | 100 | 1.47 | *Klebsiella grimontii* 99.1678 % | Yes | No | 557 | 4 | 6 | 19 | 10 | 70 | 7 | 31 |
| GCF_001053665.1 | 100 | 0.47 | *Klebsiella grimontii* 99.1683 % | Yes | No | 408 | 21 | 6 | 19 | 10 | 46 | 7 | 9 |
| GCF_008120915.1 | 100 | 0.21 | *Klebsiella grimontii* 99.1774 % | Yes | No | 361 | 6 | 6 | 14 | 10 | 46 | 8 | 33 |
| GCF_902162205.1 | 100 | 0.69 | *Klebsiella grimontii* 99.179 % | Yes | No | 186 | 21 | 6 | 14 | 10 | 62 | 7 | 9 |
| GCF_902363155.1 | 100 | 1.05 | *Klebsiella grimontii* 99.1805 % | Yes | No | 216 | 4 | 6 | 19 | 10 | 46 | 24 | 31 |
| GCF_003416995.1 | 100 | 1.01 | *Klebsiella grimontii* 99.1813 % | Yes | Yes | 172 | 6 | 6 | 19 | 11 | 46 | 7 | 9 |
| GCF_001060405.1 | 100 | 0.59 | *Klebsiella grimontii* 99.1871 % | Yes | No | 408 | 21 | 6 | 19 | 10 | 46 | 7 | 9 |
| GCF_013590775.1 | 100 | 0.48 | *Klebsiella grimontii* 99.1877 % | Yes | No | 215 | 21 | 6 | 14 | 10 | 46 | 11 | 6 |
| GCF_020118015.1 | 100 | 0.22 | *Klebsiella grimontii* 99.1915 % | Yes | No | 411 | 5 | 6 | 19 | 10 | 109 | 5 | 6 |
| GCF_022543725.1 | 100 | 0.5 | *Klebsiella grimontii* 99.1925 % | Yes | No | 371 | 15 | 6 | 19 | 10 | 46 | 24 | 6 |
| GCF_008120465.1 | 100 | 0.17 | *Klebsiella grimontii* 99.1978 % | Yes | No | - | ~15 | 6 | 19 | 10 | 46 | 5 | ~113 |
| GCF_013735435.1 | 100 | 0.92 | *Klebsiella grimontii* 99.1987 % | Yes | No | - | 5 | 6 | 19 | ~10 | 46 | 11 | 9 |
| GCF_013926505.1 | 100 | 0.48 | *Klebsiella grimontii* 99.1992 % | Yes | No | 168 | 15 | 6 | 39 | 10 | 46 | 24 | 6 |
| GCF_004104525.2 | 100 | 0.48 | *Klebsiella grimontii* 99.2 % | Yes | No | - | 21 | 6 | 14 | 10 | ~46 | 11 | 6 |
| GCF_902161015.1 | 100 | 0.05 | *Klebsiella grimontii* 99.202 % | Yes | No | 408 | 21 | 6 | 19 | 10 | 46 | 7 | 9 |
| GCF_001054995.1 | 100 | 0.62 | *Klebsiella grimontii* 99.2046 % | Yes | No | 215 | 21 | 6 | 14 | 10 | 46 | 11 | 6 |
| GCF_020118875.1 | 100 | 0.3 | *Klebsiella grimontii* 99.2066 % | Yes | No | 411 | 5 | 6 | 19 | 10 | 109 | 5 | 6 |
| GCF_902161025.1 | 100 | 0.08 | *Klebsiella grimontii* 99.2087 % | Yes | No | 408 | 21 | 6 | 19 | 10 | 46 | 7 | 9 |
| GCF_902165685.1 | 100 | 0.51 | *Klebsiella grimontii* 99.2114 % | Yes | No | 227 | 21 | 6 | 56 | 10 | 46 | 11 | 6 |
| GCF_003417005.1 | 100 | 3.73 | *Klebsiella grimontii* 99.2132 % | Yes | Yes | 172 | 6 | 6 | 19 | 11 | 46 | 7 | 9 |
| GCF_020119895.1 | 100 | 0.57 | *Klebsiella grimontii* 99.2143 % | Yes | No | 517 | 4 | 6 | 26 | 10 | 46 | 7 | 81 |
| GCF_902160315.1 | 100 | 0.15 | *Klebsiella grimontii* 99.2152 % | Yes | No | 184 | 4 | 6 | 26 | 10 | 46 | 5 | 6 |
| GCF_902161135.1 | 100 | 0.06 | *Klebsiella grimontii* 99.2152 % | Yes | No | 408 | 21 | 6 | 19 | 10 | 46 | 7 | 9 |
| GCF_902159585.1 | 100 | 0.18 | *Klebsiella grimontii* 99.2176 % | Yes | No | 408 | 21 | 6 | 19 | 10 | 46 | 7 | 9 |
| GCF_902161235.1 | 100 | 0.06 | *Klebsiella grimontii* 99.2193 % | Yes | No | 408 | 21 | 6 | 19 | 10 | 46 | 7 | 9 |
| GCF_003339485.1 | 100 | 0.38 | *Klebsiella grimontii* 99.2194 % | Yes | Yes | 216 | 4 | 6 | 19 | 10 | 46 | 24 | 31 |
| GCF_009721385.1 | 100 | 0.19 | *Klebsiella grimontii* 99.2196 % | Yes | No | - | 4 | 6 | 19 | 10 | 46 | ~5 | 9 |
| GCF_020117965.1 | 100 | 0.16 | *Klebsiella grimontii* 99.2197 % | Yes | No | - | 4 | 6 | 19 | 10 | 46 | ~5 | 9 |
| GCF_020119845.1 | 100 | 0.9 | *Klebsiella grimontii* 99.223 % | Yes | Yes | 172 | 6 | 6 | 19 | 11 | 46 | 7 | 9 |
| GCF_902161285.1 | 100 | 0.2 | *Klebsiella grimontii* 99.2243 % | Yes | No | 408 | 21 | 6 | 19 | 10 | 46 | 7 | 9 |
| GCF_020117895.1 | 100 | 0.63 | *Klebsiella grimontii* 99.2253 % | Yes | Yes | 490 | 5 | 6 | 18 | 10 | 46 | 23 | 9 |
| GCF_022539995.1 | 100 | 0.61 | *Klebsiella grimontii* 99.2253 % | Yes | No | 371 | 15 | 6 | 19 | 10 | 46 | 24 | 6 |
| GCF_020117115.1 | 100 | 0.25 | *Klebsiella grimontii* 99.2258 % | Yes | No | 246 | 5 | 10 | 59 | 10 | 46 | 7 | 9 |
| GCF_902162255.1 | 100 | 0.28 | *Klebsiella grimontii* 99.2279 % | Yes | No | 291 | 5 | 6 | 66 | 10 | 95 | 5 | 9 |
| GCF_902164985.1 | 100 | 0.23 | *Klebsiella grimontii* 99.228 % | Yes | No | 411 | 5 | 6 | 19 | 10 | 109 | 5 | 6 |
| GCF_902160845.1 | 100 | 0.3 | *Klebsiella grimontii* 99.2283 % | Yes | No | 216 | 4 | 6 | 19 | 10 | 46 | 24 | 31 |
| GCF_019678365.1 | 100 | 0.27 | *Klebsiella grimontii* 99.2288 % | Yes | No | 440 | 5 | 6 | 19 | 10 | 70 | 5 | 6 |
| GCF_900083595.1 | 100 | 0.45 | *Klebsiella grimontii* 99.2289 % | Yes | No | 186 | 21 | 6 | 14 | 10 | 62 | 7 | 9 |
| GCF_008120425.1 | 100 | 0.33 | *Klebsiella grimontii* 99.2313 % | Yes | No | 216 | 4 | 6 | 19 | 10 | 46 | 24 | 31 |
| GCF_019428385.1 | 100 | 0.22 | *Klebsiella grimontii* 99.2357 % | Yes | No | 555 | 4 | 6 | 19 | 10 | 46 | 7 | 6 |
| GCF_020115715.1 | 100 | 0.34 | *Klebsiella grimontii* 99.2365 % | Yes | Yes | 172 | 6 | 6 | 19 | 11 | 46 | 7 | 9 |
| GCF_003417035.1 | 100 | 0.33 | *Klebsiella grimontii* 99.2382 % | Yes | Yes | 172 | 6 | 6 | 19 | 11 | 46 | 7 | 9 |
| GCF_022543655.1 | 100 | 0.59 | *Klebsiella grimontii* 99.2397 % | Yes | No | 371 | 15 | 6 | 19 | 10 | 46 | 24 | 6 |
| GCF_004343645.1 | 100 | 0.18 | *Klebsiella grimontii* 99.2419 % | Yes | No | 567 | 6 | 6 | 18 | 10 | 46 | 8 | 16 |
| GCF_902162845.1 | 100 | 0.31 | *Klebsiella grimontii* 99.2431 % | Yes | No | 168 | 15 | 6 | 39 | 10 | 46 | 24 | 6 |
| GCF_020120855.1 | 100 | 0.18 | *Klebsiella grimontii* 99.244 % | Yes | No | - | 4 | 6 | 19 | 10 | 46 | ~5 | 9 |
| GCF_022543715.1 | 100 | 0.59 | *Klebsiella grimontii* 99.2493 % | Yes | No | 371 | 15 | 6 | 19 | 10 | 46 | 24 | 6 |
| GCF_022539845.1 | 100 | 0.61 | *Klebsiella grimontii* 99.2539 % | Yes | No | 371 | 15 | 6 | 19 | 10 | 46 | 24 | 6 |
| GCF_902166675.1 | 99.72 | 0.03 | *Klebsiella grimontii* 99.2559 % | No | No | - | 21 | 6 | ~19 | 10 | 46 | 7 | 9 |
| GCF_902164185.1 | 100 | 0.11 | *Klebsiella grimontii* 99.2571 % | Yes | No | 408 | 21 | 6 | 19 | 10 | 46 | 7 | 9 |
| GCF_016811025.1 | 100 | 0.19 | *Klebsiella grimontii* 99.2572 % | No | No | 261 | 4 | 6 | 19 | 10 | 46 | 5 | 6 |
| GCF_902160775.1 | 100 | 0.3 | *Klebsiella grimontii* 99.2593 % | Yes | No | 216 | 4 | 6 | 19 | 10 | 46 | 24 | 31 |
| GCF_902160755.1 | 100 | 0.86 | *Klebsiella grimontii* 99.2644 % | Yes | No | 562 | 5 | 6 | 14 | 10 | 46 | 7 | 9 |
| GCF_013635015.1 | 100 | 0.35 | *Klebsiella grimontii* 99.2694 % | Yes | No | 215 | 21 | 6 | 14 | 10 | 46 | 11 | 6 |
| GCF_902162765.1 | 100 | 0.72 | *Klebsiella grimontii* 99.2738 % | Yes | No | 215 | 21 | 6 | 14 | 10 | 46 | 11 | 6 |
| GCF_019677835.1 | 100 | 0.13 | *Klebsiella grimontii* 99.2854 % | Yes | No | - | 4 | 6 | ~19 | 10 | 46 | 7 | 6 |
| GCF_019428125.1 | 100 | 0.19 | *Klebsiella grimontii* 99.2875 % | Yes | No | 261 | 4 | 6 | 19 | 10 | 46 | 5 | 6 |
| GCF_902161425.1 | 100 | 0.48 | *Klebsiella grimontii* 99.2912 % | Yes | No | 186 | 21 | 6 | 14 | 10 | 62 | 7 | 9 |
| GCF_902164135.1 | 100 | 0.1 | *Klebsiella grimontii* 99.2937 % | Yes | No | 408 | 21 | 6 | 19 | 10 | 46 | 7 | 9 |
| GCF_019428325.1 | 100 | 0.12 | *Klebsiella grimontii* 99.2946 % | Yes | No | 555 | 4 | 6 | 19 | 10 | 46 | 7 | 6 |
| GCF_019334465.1 | 100 | 1.58 | *Klebsiella grimontii* 99.2954 % | Yes | No | 488 | 4 | 6 | 19 | 10 | 46 | 5 | 9 |
| GCF_001066775.1 | 100 | 0.2 | *Klebsiella grimontii* 99.2963 % | Yes | No | 563 | 5 | 6 | 18 | 10 | 46 | 8 | 31 |
| GCF_902164055.1 | 100 | 0.11 | *Klebsiella grimontii* 99.2973 % | Yes | No | 408 | 21 | 6 | 19 | 10 | 46 | 7 | 9 |
| GCF_902164975.1 | 100 | 0.5 | *Klebsiella grimontii* 99.2995 % | Yes | No | 341 | 21 | 6 | 70 | 10 | 46 | 11 | 6 |
| GCF_900451335.1 | 99.99 | 1.58 | *Klebsiella grimontii* 99.3011 % | No | No | 517 | 4 | 6 | 26 | 10 | 46 | 7 | 81 |
| GCF_902166735.1 | 100 | 0.04 | *Klebsiella grimontii* 99.3015 % | No | No | - | 21 | 6 | ~19 | 10 | 46 | 7 | 9 |
| GCF_022014295.1 | 100 | 0.2 | *Klebsiella grimontii* 99.303 % | Yes | No | 283 | 5 | 6 | 19 | 11 | 46 | 7 | 32 |
| GCF_022556395.1 | 100 | 0.2 | *Klebsiella grimontii* 99.303 % | Yes | No | 283 | 5 | 6 | 19 | 11 | 46 | 7 | 32 |
| GCF_019428165.1 | 100 | 0.19 | *Klebsiella grimontii* 99.3032 % | Yes | No | 261 | 4 | 6 | 19 | 10 | 46 | 5 | 6 |
| GCF_020479545.1 | 100 | 0.22 | *Klebsiella grimontii* 99.3097 % | Yes | No | 440 | 5 | 6 | 19 | 10 | 70 | 5 | 6 |
| GCF_900083605.1 | 100 | 0.34 | *Klebsiella grimontii* 99.3105 % | Yes | No | 216 | 4 | 6 | 19 | 10 | 46 | 24 | 31 |
| GCF_019426485.1 | 100 | 0.34 | *Klebsiella grimontii* 99.3188 % | Yes | No | 290 | 5 | 6 | 14 | 11 | 70 | 7 | 6 |
| GCF_902166545.1 | 100 | 0.19 | *Klebsiella grimontii* 99.3201 % | No | No | 216 | 4 | 6 | 19 | 10 | 46 | 24 | 31 |
| GCF_000308735.2 | 99.97 | 3.81 | *Klebsiella grimontii* 99.3214 % | No | No | 104 | 4 | 6 | 11 | 10 | 46 | 27 | 9 |
| GCF_009905335.1 | 100 | 0.14 | *Klebsiella grimontii* 99.325 % | No | No | - | 15 | 6 | ~54 | 10 | 46 | 5 | 9 |
| GCF_020119035.1 | 100 | 0.35 | *Klebsiella grimontii* 99.3266 % | Yes | No | 216 | 4 | 6 | 19 | 10 | 46 | 24 | 31 |
| GCF_014655015.1 | 100 | 0.62 | *Klebsiella grimontii* 99.3291 % | Yes | No | 262 | 6 | 6 | 19 | 35 | 46 | 7 | 9 |
| GCF_000427015.1 | 100 | 0.17 | *Klebsiella grimontii* 99.3345 % | Yes | No | 445 | 5 | 6 | 19 | 10 | 46 | 5 | 98 |
| GCF_002556465.1 | 100 | 0.2 | *Klebsiella grimontii* 99.3347 % | Yes | No | - | 5 | 6 | 5 | 10 | 46 | 7 | 103 |
| GCF_008364425.1 | 100 | 0.2 | *Klebsiella grimontii* 99.3347 % | Yes | No | - | 5 | 6 | 5 | 10 | 46 | 7 | 103 |
| GCF_020116695.1 | 100 | 0.12 | *Klebsiella grimontii* 99.352 % | Yes | No | 561 | 5 | 6 | 14 | 10 | 46 | 5 | 9 |
| GCF_015183075.1 | 100 | 0.23 | *Klebsiella grimontii* 99.3535 % | Yes | No | 104 | 4 | 6 | 11 | 10 | 46 | 27 | 9 |
| GCF_002090195.1 | 100 | 0.22 | *Klebsiella grimontii* 99.3572 % | No | No | 104 | 4 | 6 | 11 | 10 | 46 | 27 | 9 |
| GCF_020117435.1 | 100 | 0.28 | *Klebsiella grimontii* 99.359 % | Yes | No | 576 | 15 | 6 | 10 | 11 | 46 | 8 | 16 |
| GCF_020117575.1 | 100 | 0.17 | *Klebsiella grimontii* 99.3679 % | Yes | No | 261 | 4 | 6 | 19 | 10 | 46 | 5 | 6 |
| GCF_015074825.1 | 100 | 0.14 | *Klebsiella grimontii* 99.3714 % | No | No | 104 | 4 | 6 | 11 | 10 | 46 | 27 | 9 |
| GCF_020118435.1 | 100 | 0.1 | *Klebsiella grimontii* 99.3808 % | No | No | - | 4 | 6 | ~54 | 35 | 46 | ~5 | ~90 |
| GCF_002559635.1 | 100 | 0.29 | *Klebsiella grimontii* 99.3829 % | Yes | No | - | 4 | 6 | 19 | 10 | 46 | 5 | ~9 |
| GCF_901542455.1 | 100 | 0.44 | *Klebsiella grimontii* 99.3926 % | No | No | - | 21 | 6 | 14 | 10 | ~62 | 7 | 9 |
| GCF_001633115.1 | 100 | 0.17 | *Klebsiella grimontii* 99.3968 % | No | No | 104 | 4 | 6 | 11 | 10 | 46 | 27 | 9 |
| GCF_020117935.1 | 100 | 0.15 | *Klebsiella grimontii* 99.4025 % | No | No | 261 | 4 | 6 | 19 | 10 | 46 | 5 | 6 |
| GCF_020117975.1 | 100 | 0.12 | *Klebsiella grimontii* 99.4033 % | No | No | - | 4 | 6 | ~54 | 10 | 46 | 5 | 33 |
| GCF_020117755.1 | 100 | 0.12 | *Klebsiella grimontii* 99.4077 % | Yes | No | - | 6 | 6 | - | 10 | 46 | 7 | 17 |
| GCF_020117855.1 | 100 | 0.17 | *Klebsiella grimontii* 99.412 % | No | No | 238 | 4 | 6 | 57 | 10 | 46 | 8 | 6 |
| GCF_020117835.1 | 100 | 0.1 | *Klebsiella grimontii* 99.4129 % | No | No | 558 | 4 | 6 | 39 | 10 | 46 | 7 | 16 |
| GCF_020117915.1 | 100 | 0.15 | *Klebsiella grimontii* 99.4148 % | No | No | - | 6 | 6 | ~54 | 10 | 46 | 8 | ~16 |
| GCF_019679055.1 | 100 | 0.11 | *Klebsiella grimontii* 99.4171 % | Yes | No | 186 | 21 | 6 | 14 | 10 | 62 | 7 | 9 |
| GCF_020117635.1 | 100 | 0.12 | *Klebsiella grimontii* 99.4174 % | No | No | 578 | 15 | 6 | 26 | 10 | 46 | 11 | 9 |
| GCF_020120845.1 | 100 | 0.12 | *Klebsiella grimontii* 99.4209 % | No | No | - | 4 | 6 | ~54 | 10 | 46 | 5 | 33 |
| GCF_020117615.1 | 100 | 0.17 | *Klebsiella grimontii* 99.4231 % | No | No | 326 | 4 | 6 | 14 | 35 | 96 | 5 | 6 |
| GCF_902164675.1 | 100 | 0.12 | *Klebsiella grimontii* 99.4233 % | No | No | 560 | 5 | 6 | 11 | 11 | 46 | 5 | 16 |
| GCF_019678325.1 | 100 | 0.09 | *Klebsiella grimontii* 99.4242 % | No | No | - | 5 | 6 | 18 | 10 | ~46 | 8 | 16 |
| GCF_019913035.1 | 100 | 0.15 | *Klebsiella grimontii* 99.4266 % | No | No | 104 | 4 | 6 | 11 | 10 | 46 | 27 | 9 |
| GCF_019428245.1 | 100 | 0.07 | *Klebsiella grimontii* 99.429 % | Yes | No | 555 | 4 | 6 | 19 | 10 | 46 | 7 | 6 |
| GCF_020120815.1 | 100 | 0.08 | *Klebsiella grimontii* 99.4294 % | No | No | 261 | 4 | 6 | 19 | 10 | 46 | 5 | 6 |
| GCF_016653045.1 | 100 | 2.79 | *Klebsiella grimontii* 99.4385 % | Yes | No | 567 | 6 | 6 | 18 | 10 | 46 | 8 | 16 |
| GCF_019679095.1 | 100 | 0.07 | *Klebsiella grimontii* 99.4423 % | No | No | - | 6 | 6 | 14 | 10 | 46 | 7 | 92 |
| GCF_020118535.1 | 100 | 0.16 | *Klebsiella grimontii* 99.4496 % | Yes | No | 480 | 4 | 6 | 14 | 10 | 46 | 7 | 31 |
| GCF_002880715.1 | 100 | 0.07 | *Klebsiella grimontii* 99.4546 % | No | No | 316 | 5 | 6 | 19 | 11 | 46 | 5 | 9 |
| GCF_902164895.1 | 100 | 0.08 | *Klebsiella grimontii* 99.4735 % | No | No | 168 | 15 | 6 | 39 | 10 | 46 | 24 | 6 |
| GCF_902164065.1 | 100 | 0.14 | *Klebsiella grimontii* 99.5166 % | Yes | No | 246 | 5 | 10 | 59 | 10 | 46 | 7 | 9 |
| GCF_902163185.1 | 100 | 0.4 | *Klebsiella grimontii* 99.5167 % | Yes | No | 565 | 5 | 6 | 26 | 10 | 46 | 5 | 6 |
| GCF_022543775.1 | 100 | 1.01 | *Klebsiella grimontii* 99.5702 % | Yes | No | 319 | 4 | 6 | 14 | 11 | 46 | 11 | 9 |
| GCF_022543755.1 | 100 | 0.69 | *Klebsiella grimontii* 99.832 % | Yes | No | 319 | 4 | 6 | 14 | 11 | 46 | 11 | 9 |
| GCF_003261575.2 | 100 | 1 | *Klebsiella huaxiensis* 100 % | Yes | No | - | ~31 | - | - | ~81 | - | 24? | - |
| GCF_902158605.1 | 100 | 2.26 | *Klebsiella huaxiensis* 99.1222 % | No | No | - | ~31 | - | - | ~81 | - | 24? | - |
| GCF_902158625.1 | 100 | 1.19 | *Klebsiella huaxiensis* 99.545 % | Yes | No | - | ~31 | - | - | ~81 | - | 24? | - |
| GCF_002925905.1 | 100 | 0.69 | *Klebsiella michiganensis* 100 % | Yes | No | 84 | 3 | 8 | 17 | 21 | 40 | 17 | 29 |
| GCF_901556995.1 | 100 | 0.69 | *Klebsiella michiganensis* 100 % | Yes | No | 84 | 3 | 8 | 17 | 21 | 40 | 17 | 29 |
| GCF_013732655.1 | 100 | 2.19 | *Klebsiella michiganensis* 97.3581 % | Yes | No | 127 | 14 | 4 | 15 | 4 | 41 | 3 | 4 |
| GCF_019460145.1 | 100 | 0.87 | *Klebsiella michiganensis* 97.3707 % | Yes | Yes | 91 | 3 | 4 | 15 | 8 | 12 | 3 | 35 |
| GCF_021391575.1 | 100 | 0.68 | *Klebsiella michiganensis* 97.3785 % | Yes | No | 91 | 3 | 4 | 15 | 8 | 12 | 3 | 35 |
| GCF_015356405.1 | 100 | 0.56 | *Klebsiella michiganensis* 97.3855 % | Yes | No | 95 | 3 | 24 | 15 | 4 | 43 | 6 | 4 |
| GCF_902160725.1 | 100 | 1.22 | *Klebsiella michiganensis* 97.3903 % | Yes | No | - | 3 | 4 | 15 | 8 | 33 | 3 | ~35 |
| GCF_002853055.1 | 100 | 1.24 | *Klebsiella michiganensis* 97.3913 % | Yes | No | 92 | 3 | 4 | 15 | 8 | 18 | 6 | 37 |
| GCF_902159535.1 | 100 | 1.29 | *Klebsiella michiganensis* 97.3966 % | Yes | No | - | 3 | 4 | 15 | 8 | 33 | 3 | ~35 |
| GCF_902754085.1 | 100 | 0.56 | *Klebsiella michiganensis* 97.3975 % | Yes | No | 95 | 3 | 24 | 15 | 4 | 43 | 6 | 4 |
| GCF_013610655.1 | 100 | 0.96 | *Klebsiella michiganensis* 97.4113 % | Yes | Yes | 95 | 3 | 24 | 15 | 4 | 43 | 6 | 4 |
| GCF_022540125.1 | 100 | 0.75 | *Klebsiella michiganensis* 97.4134 % | Yes | No | 183 | 3 | 4 | 47 | 8 | 43 | 3 | 4 |
| GCF_018257735.1 | 100 | 0.33 | *Klebsiella michiganensis* 97.4248 % | Yes | No | 579 | 30 | 4 | 30 | 8 | 25 | 6 | 4 |
| GCF_001052525.1 | 100 | 0.58 | *Klebsiella michiganensis* 97.437 % | Yes | Yes | 83 | 3 | 4 | 30 | 8 | 39 | 26 | 35 |
| GCF_014655075.1 | 100 | 0.91 | *Klebsiella michiganensis* 97.4484 % | Yes | No | 93 | 3 | 26 | 31 | 8 | 33 | 6 | 4 |
| GCF_015356395.1 | 100 | 0.44 | *Klebsiella michiganensis* 97.4516 % | Yes | No | 95 | 3 | 24 | 15 | 4 | 43 | 6 | 4 |
| GCF_000567805.1 | 100 | 0.86 | *Klebsiella michiganensis* 97.4677 % | Yes | No | 29 | 3 | 4 | 15 | 8 | 18 | 6 | 11 |
| GCF_902159805.1 | 100 | 0.33 | *Klebsiella michiganensis* 97.4693 % | Yes | No | 29 | 3 | 4 | 15 | 8 | 18 | 6 | 11 |
| GCF_014856365.1 | 100 | 0.43 | *Klebsiella michiganensis* 97.4761 % | Yes | No | 548 | 3 | 15 | 15 | 8 | 25 | 3 | 35 |
| GCF_014856335.1 | 100 | 0.44 | *Klebsiella michiganensis* 97.4862 % | Yes | No | 548 | 3 | 15 | 15 | 8 | 25 | 3 | 35 |
| GCF_004115315.1 | 100 | 1.06 | *Klebsiella michiganensis* 97.4917 % | Yes | Yes | 29 | 3 | 4 | 15 | 8 | 18 | 6 | 11 |
| GCF_902164635.1 | 100 | 0.37 | *Klebsiella michiganensis* 97.4921 % | No | No | - | 3 | 4 | 15 | 8 | 33 | 3 | ~35 |
| GCF_902163385.1 | 100 | 0.51 | *Klebsiella michiganensis* 97.4946 % | Yes | No | 29 | 3 | 4 | 15 | 8 | 18 | 6 | 11 |
| GCF_902159745.1 | 100 | 0.33 | *Klebsiella michiganensis* 97.4966 % | Yes | No | 29 | 3 | 4 | 15 | 8 | 18 | 6 | 11 |
| GCF_900407165.1 | 100 | 1.26 | *Klebsiella michiganensis* 97.5039 % | Yes | Yes | 533 | 3 | 4 | 42 | 4 | 43 | 6 | 4 |
| GCF_015999385.1 | 100 | 0.39 | *Klebsiella michiganensis* 97.5109 % | Yes | No | 330 | 3 | 4 | 15 | 4 | 33 | 45 | 4 |
| GCF_022539825.1 | 100 | 0.53 | *Klebsiella michiganensis* 97.5112 % | Yes | No | 183 | 3 | 4 | 47 | 8 | 43 | 3 | 4 |
| GCF_014856375.1 | 100 | 0.49 | *Klebsiella michiganensis* 97.5145 % | Yes | No | 548 | 3 | 15 | 15 | 8 | 25 | 3 | 35 |
| GCF_902166415.1 | 100 | 0.29 | *Klebsiella michiganensis* 97.5165 % | No | No | 29 | 3 | 4 | 15 | 8 | 18 | 6 | 11 |
| GCF_014856345.1 | 100 | 0.42 | *Klebsiella michiganensis* 97.5271 % | Yes | No | 548 | 3 | 15 | 15 | 8 | 25 | 3 | 35 |
| GCF_900083695.1 | 100 | 0.33 | *Klebsiella michiganensis* 97.5287 % | Yes | No | 92 | 3 | 4 | 15 | 8 | 18 | 6 | 37 |
| GCF_013926025.1 | 100 | 0.8 | *Klebsiella michiganensis* 97.5394 % | Yes | No | 554 | 3 | 57 | 79 | 8 | 25 | 6 | 4 |
| GCF_018138485.1 | 100 | 0.47 | *Klebsiella michiganensis* 97.5417 % | Yes | No | 349 | 3 | 50 | 15 | 8 | 41 | 22 | 4 |
| GCF_014856435.1 | 100 | 0.38 | *Klebsiella michiganensis* 97.5428 % | Yes | No | 548 | 3 | 15 | 15 | 8 | 25 | 3 | 35 |
| GCF_014856385.1 | 100 | 0.39 | *Klebsiella michiganensis* 97.5429 % | Yes | No | 548 | 3 | 15 | 15 | 8 | 25 | 3 | 35 |
| GCF_013392155.1 | 100 | 3.04 | *Klebsiella michiganensis* 97.5467 % | Yes | No | 183 | 3 | 4 | 47 | 8 | 43 | 3 | 4 |
| GCF_901563895.1 | 100 | 0.26 | *Klebsiella michiganensis* 97.5518 % | Yes | No | 183 | 3 | 4 | 47 | 8 | 43 | 3 | 4 |
| GCF_001056765.1 | 100 | 0.46 | *Klebsiella michiganensis* 97.5544 % | Yes | No | 92 | 3 | 4 | 15 | 8 | 18 | 6 | 37 |
| GCF_902159765.1 | 100 | 0.28 | *Klebsiella michiganensis* 97.5548 % | Yes | No | 29 | 3 | 4 | 15 | 8 | 18 | 6 | 11 |
| GCF_004024445.1 | 100 | 1.52 | *Klebsiella michiganensis* 97.5583 % | Yes | No | 127 | 14 | 4 | 15 | 4 | 41 | 3 | 4 |
| GCF_002152325.1 | 100 | 0.25 | *Klebsiella michiganensis* 97.5605 % | Yes | No | - | 3 | 50 | 15 | 22 | ~18 | 6 | 4 |
| GCF_902165095.1 | 100 | 0.23 | *Klebsiella michiganensis* 97.5647 % | Yes | No | 29 | 3 | 4 | 15 | 8 | 18 | 6 | 11 |
| GCF_014267425.1 | 100 | 0.24 | *Klebsiella michiganensis* 97.5671 % | Yes | No | 549 | 3 | 21 | 15 | 8 | 18 | 6 | 4 |
| GCF_020117795.1 | 100 | 0.12 | *Klebsiella michiganensis* 97.5687 % | Yes | No | 29 | 3 | 4 | 15 | 8 | 18 | 6 | 11 |
| GCF_016620455.1 | 100 | 0.83 | *Klebsiella michiganensis* 97.5753 % | Yes | No | 534 | 3 | 4 | 49 | 8 | 12 | 3 | 4 |
| GCF_018138395.1 | 100 | 0.47 | *Klebsiella michiganensis* 97.5817 % | Yes | No | 349 | 3 | 50 | 15 | 8 | 41 | 22 | 4 |
| GCF_010093005.1 | 100 | 1.65 | *Klebsiella michiganensis* 97.583 % | Yes | No | - | ~45 | 61 | ~82 | ~74 | 20 | ~52 | 102 |
| GCF_000247835.1 | 100 | 0.47 | *Klebsiella michiganensis* 97.5866 % | Yes | No | 29 | 3 | 4 | 15 | 8 | 18 | 6 | 11 |
| GCF_013624375.1 | 100 | 1.29 | *Klebsiella michiganensis* 97.5968 % | Yes | No | 551 | 3 | 24 | 15 | 8 | 43 | 6 | 4 |
| GCF_000293135.1 | 100 | 3.25 | *Klebsiella michiganensis* 97.6117 % | Yes | No | 82 | 14 | 24 | 15 | 8 | 18 | 6 | 4 |
| GCF_018138465.1 | 100 | 0.34 | *Klebsiella michiganensis* 97.6191 % | Yes | Yes | 82 | 14 | 24 | 15 | 8 | 18 | 6 | 4 |
| GCF_019678505.1 | 100 | 0.3 | *Klebsiella michiganensis* 97.6371 % | Yes | No | 29 | 3 | 4 | 15 | 8 | 18 | 6 | 11 |
| GCF_014856145.1 | 100 | 0.2 | *Klebsiella michiganensis* 97.6405 % | Yes | No | 29 | 3 | 4 | 15 | 8 | 18 | 6 | 11 |
| GCF_014856135.1 | 100 | 0.27 | *Klebsiella michiganensis* 97.6616 % | Yes | No | 29 | 3 | 4 | 15 | 8 | 18 | 6 | 11 |
| GCF_023093835.1 | 100 | 0.31 | *Klebsiella michiganensis* 97.6679 % | No | No | 330 | 3 | 4 | 15 | 4 | 33 | 45 | 4 |
| GCF_017815775.1 | 100 | 0.4 | *Klebsiella michiganensis* 97.6849 % | No | No | 86 | 3 | 4 | 15 | 32 | 41 | 3 | 35 |
| GCF_014129405.1 | 100 | 0.47 | *Klebsiella michiganensis* 97.696 % | Yes | No | 86 | 3 | 4 | 15 | 32 | 41 | 3 | 35 |
| GCF_020117475.1 | 100 | 0.35 | *Klebsiella michiganensis* 97.7043 % | Yes | No | 29 | 3 | 4 | 15 | 8 | 18 | 6 | 11 |
| GCF_009930855.1 | 100 | 1.34 | *Klebsiella michiganensis* 97.7079 % | Yes | No | 383 | 3 | 4 | 75 | 4 | 25 | 3 | 28 |
| GCF_009825595.1 | 100 | 0.92 | *Klebsiella michiganensis* 97.8046 % | No | No | - | 45 | 61 | ~82 | ~74 | 20 | 52 | 102 |
| GCF_902163225.1 | 100 | 2.11 | *Klebsiella michiganensis* 98.2971 % | Yes | No | 151 | 3 | 5 | 21 | 13 | 20 | 6 | 12 |
| GCF_000783895.2 | 100 | 1.24 | *Klebsiella michiganensis* 98.2984 % | Yes | No | 537 | 3 | 5 | 7 | 65 | 20 | 6 | 5 |
| GCF_021228695.1 | 100 | 1.32 | *Klebsiella michiganensis* 98.306 % | Yes | No | 309 | 3 | 5 | 21 | 13 | 24 | 33 | 20 |
| GCF_020117775.1 | 100 | 3.1 | *Klebsiella michiganensis* 98.3064 % | Yes | No | 43 | 3 | 5 | 21 | 20 | 11 | 6 | 20 |
| GCF_902163215.1 | 100 | 0.74 | *Klebsiella michiganensis* 98.3108 % | Yes | No | 151 | 3 | 5 | 21 | 13 | 20 | 6 | 12 |
| GCF_902163205.1 | 100 | 2.71 | *Klebsiella michiganensis* 98.3154 % | Yes | No | 151 | 3 | 5 | 21 | 13 | 20 | 6 | 12 |
| GCF_015721525.1 | 100 | 1.83 | *Klebsiella michiganensis* 98.3185 % | Yes | No | 43 | 3 | 5 | 21 | 20 | 11 | 6 | 20 |
| GCF_902164835.1 | 100 | 0.84 | *Klebsiella michiganensis* 98.3231 % | Yes | No | - | 3 | 5 | 21 | 13 | 74 | ~6 | 12 |
| GCF_022869885.1 | 100 | 2.78 | *Klebsiella michiganensis* 98.3315 % | Yes | No | 202 | 3 | 5 | 21 | 13 | 74 | 6 | 12 |
| GCF_016618215.1 | 100 | 0.49 | *Klebsiella michiganensis* 98.335 % | Yes | No | 342 | 3 | 5 | 40 | 64 | 24 | 6 | 82 |
| GCF_001028875.1 | 100 | 0.86 | *Klebsiella michiganensis* 98.3421 % | Yes | No | 202 | 3 | 5 | 21 | 13 | 74 | 6 | 12 |
| GCF_020120175.1 | 100 | 2.73 | *Klebsiella michiganensis* 98.3454 % | Yes | No | 43 | 3 | 5 | 21 | 20 | 11 | 6 | 20 |
| GCF_002906395.1 | 100 | 2.39 | *Klebsiella michiganensis* 98.3478 % | Yes | No | 202 | 3 | 5 | 21 | 13 | 74 | 6 | 12 |
| GCF_014655105.1 | 100 | 1.45 | *Klebsiella michiganensis* 98.3528 % | Yes | No | 43 | 3 | 5 | 21 | 20 | 11 | 6 | 20 |
| GCF_002290285.1 | 100 | 0.65 | *Klebsiella michiganensis* 98.3537 % | Yes | No | 27 | 3 | 5 | 7 | 13 | 17 | 4 | 5 |
| GCF_002918655.1 | 100 | 1.04 | *Klebsiella michiganensis* 98.3546 % | Yes | No | 85 | 3 | 5 | 21 | 13 | 24 | 6 | 19 |
| GCF_902162645.1 | 100 | 1.05 | *Klebsiella michiganensis* 98.3548 % | Yes | No | 108 | 3 | 5 | 21 | 20 | 24 | 6 | 30 |
| GCF_001028885.1 | 100 | 0.86 | *Klebsiella michiganensis* 98.3551 % | Yes | No | 202 | 3 | 5 | 21 | 13 | 74 | 6 | 12 |
| GCF_002216835.1 | 100 | 0.61 | *Klebsiella michiganensis* 98.3571 % | Yes | No | 27 | 3 | 5 | 7 | 13 | 17 | 4 | 5 |
| GCF_015721545.1 | 100 | 1.22 | *Klebsiella michiganensis* 98.3601 % | Yes | No | 43 | 3 | 5 | 21 | 20 | 11 | 6 | 20 |
| GCF_014655115.1 | 100 | 1.9 | *Klebsiella michiganensis* 98.3614 % | Yes | No | 43 | 3 | 5 | 21 | 20 | 11 | 6 | 20 |
| GCF_002906435.1 | 100 | 1.92 | *Klebsiella michiganensis* 98.3617 % | Yes | No | 85 | 3 | 5 | 21 | 13 | 24 | 6 | 19 |
| GCF_002919605.1 | 100 | 1.82 | *Klebsiella michiganensis* 98.3663 % | Yes | No | 202 | 3 | 5 | 21 | 13 | 74 | 6 | 12 |
| GCF_002918695.1 | 100 | 1.04 | *Klebsiella michiganensis* 98.3693 % | Yes | No | 85 | 3 | 5 | 21 | 13 | 24 | 6 | 19 |
| GCF_015721745.1 | 100 | 2.4 | *Klebsiella michiganensis* 98.371 % | Yes | No | 85 | 3 | 5 | 21 | 13 | 24 | 6 | 19 |
| GCF_902164935.1 | 100 | 0.84 | *Klebsiella michiganensis* 98.373 % | Yes | No | - | 3 | 5 | 21 | 13 | 74 | ~6 | 12 |
| GCF_000276705.2 | 100 | 1.12 | *Klebsiella michiganensis* 98.3739 % | Yes | No | 27 | 3 | 5 | 7 | 13 | 17 | 4 | 5 |
| GCF_902166305.1 | 100 | 2.86 | *Klebsiella michiganensis* 98.3767 % | Yes | No | 85 | 3 | 5 | 21 | 13 | 24 | 6 | 19 |
| GCF_002919625.1 | 100 | 1.03 | *Klebsiella michiganensis* 98.3775 % | Yes | No | 85 | 3 | 5 | 21 | 13 | 24 | 6 | 19 |
| GCF_020117735.1 | 100 | 0.55 | *Klebsiella michiganensis* 98.3783 % | Yes | No | 213 | 3 | 5 | 21 | 13 | 24 | 6 | 61 |
| GCF_022343245.1 | 100 | 0.54 | *Klebsiella michiganensis* 98.3802 % | Yes | No | 27 | 3 | 5 | 7 | 13 | 17 | 4 | 5 |
| GCF_902162735.1 | 100 | 2.05 | *Klebsiella michiganensis* 98.3802 % | Yes | No | 108 | 3 | 5 | 21 | 20 | 24 | 6 | 30 |
| GCF_014654995.1 | 100 | 1.13 | *Klebsiella michiganensis* 98.3827 % | Yes | No | 85 | 3 | 5 | 21 | 13 | 24 | 6 | 19 |
| GCF_902166295.1 | 100 | 0.75 | *Klebsiella michiganensis* 98.383 % | Yes | No | 85 | 3 | 5 | 21 | 13 | 24 | 6 | 19 |
| GCF_020117865.1 | 100 | 0.62 | *Klebsiella michiganensis* 98.3852 % | Yes | No | 27 | 3 | 5 | 7 | 13 | 17 | 4 | 5 |
| GCF_902386125.1 | 100 | 0.98 | *Klebsiella michiganensis* 98.3854 % | Yes | No | 202 | 3 | 5 | 21 | 13 | 74 | 6 | 12 |
| GCF_000714655.1 | 100 | 0.98 | *Klebsiella michiganensis* 98.3859 % | Yes | No | 202 | 3 | 5 | 21 | 13 | 74 | 6 | 12 |
| GCF_002918635.1 | 100 | 2.02 | *Klebsiella michiganensis* 98.3929 % | Yes | No | 202 | 3 | 5 | 21 | 13 | 74 | 6 | 12 |
| GCF_902161955.1 | 100 | 1.16 | *Klebsiella michiganensis* 98.3934 % | Yes | No | 85 | 3 | 5 | 21 | 13 | 24 | 6 | 19 |
| GCF_015721605.1 | 100 | 0.72 | *Klebsiella michiganensis* 98.3937 % | Yes | No | 27 | 3 | 5 | 7 | 13 | 17 | 4 | 5 |
| GCF_014655005.1 | 100 | 1.08 | *Klebsiella michiganensis* 98.3972 % | Yes | No | - | 3 | 5 | ~21 | 13 | 20 | 6 | 12 |
| GCF_003590255.1 | 100 | 2.81 | *Klebsiella michiganensis* 98.4085 % | Yes | No | 43 | 3 | 5 | 21 | 20 | 11 | 6 | 20 |
| GCF_902161885.1 | 100 | 1.21 | *Klebsiella michiganensis* 98.4154 % | Yes | No | 85 | 3 | 5 | 21 | 13 | 24 | 6 | 19 |
| GCF_020116215.1 | 100 | 2.68 | *Klebsiella michiganensis* 98.4162 % | Yes | No | 108 | 3 | 5 | 21 | 20 | 24 | 6 | 30 |
| GCF_001970835.1 | 100 | 2.15 | *Klebsiella michiganensis* 98.4164 % | Yes | No | 144 | 3 | 5 | 21 | 20 | 24 | 6 | 20 |
| GCF_020117715.1 | 100 | 0.72 | *Klebsiella michiganensis* 98.4172 % | Yes | No | 85 | 3 | 5 | 21 | 13 | 24 | 6 | 19 |
| GCF_902166115.1 | 100 | 1.45 | *Klebsiella michiganensis* 98.423 % | Yes | No | 108 | 3 | 5 | 21 | 20 | 24 | 6 | 30 |
| GCF_001077175.1 | 100 | 0.91 | *Klebsiella michiganensis* 98.4248 % | Yes | No | 11 | 3 | 5 | 7 | 8 | 11 | 6 | 5 |
| GCF_010590065.1 | 100 | 0.9 | *Klebsiella michiganensis* 98.4268 % | Yes | No | 539 | 3 | 5 | 21 | 58 | 108 | 6 | 20 |
| GCF_003598595.1 | 100 | 0.71 | *Klebsiella michiganensis* 98.4324 % | Yes | No | 27 | 3 | 5 | 7 | 13 | 17 | 4 | 5 |
| GCF_008931605.1 | 100 | 2.19 | *Klebsiella michiganensis* 98.434 % | Yes | No | 108 | 3 | 5 | 21 | 20 | 24 | 6 | 30 |
| GCF_019428225.1 | 100 | 1.76 | *Klebsiella michiganensis* 98.4347 % | Yes | No | 43 | 3 | 5 | 21 | 20 | 11 | 6 | 20 |
| GCF_016905825.1 | 100 | 1.64 | *Klebsiella michiganensis* 98.4379 % | Yes | No | 144 | 3 | 5 | 21 | 20 | 24 | 6 | 20 |
| GCF_902162365.1 | 100 | 0.4 | *Klebsiella michiganensis* 98.4457 % | Yes | No | 85 | 3 | 5 | 21 | 13 | 24 | 6 | 19 |
| GCF_022343265.1 | 100 | 0.41 | *Klebsiella michiganensis* 98.4479 % | Yes | No | 27 | 3 | 5 | 7 | 13 | 17 | 4 | 5 |
| GCF_000724525.1 | 100 | 2.23 | *Klebsiella michiganensis* 98.4498 % | Yes | No | 108 | 3 | 5 | 21 | 20 | 24 | 6 | 30 |
| GCF_020116995.1 | 100 | 0.68 | *Klebsiella michiganensis* 98.4514 % | Yes | No | 43 | 3 | 5 | 21 | 20 | 11 | 6 | 20 |
| GCF_010592905.1 | 100 | 1.67 | *Klebsiella michiganensis* 98.4519 % | Yes | No | 103 | 3 | 5 | 34 | 13 | 17 | 4 | 5 |
| GCF_004102625.1 | 100 | 4.28 | *Klebsiella michiganensis* 98.453 % | Yes | No | 315 | 3 | 41 | 21 | 13 | 24 | 6 | 20 |
| GCF_002192755.1 | 100 | 0.66 | *Klebsiella michiganensis* 98.4538 % | Yes | No | 213 | 3 | 5 | 21 | 13 | 24 | 6 | 61 |
| GCF_020121095.1 | 100 | 0.4 | *Klebsiella michiganensis* 98.4599 % | Yes | No | 27 | 3 | 5 | 7 | 13 | 17 | 4 | 5 |
| GCF_902163235.1 | 100 | 0.45 | *Klebsiella michiganensis* 98.464 % | Yes | No | 213 | 3 | 5 | 21 | 13 | 24 | 6 | 61 |
| GCF_902807095.1 | 100 | 0.64 | *Klebsiella michiganensis* 98.4655 % | Yes | No | 253 | 3 | 5 | 21 | 58 | 11 | 6 | 12 |
| GCF_022540085.1 | 100 | 1.03 | *Klebsiella michiganensis* 98.4656 % | Yes | Yes | 143 | 3 | 8 | 43 | 7 | 20 | 33 | 23 |
| GCF_000735215.1 | 100 | 0.51 | *Klebsiella michiganensis* 98.467 % | Yes | No | 538 | 3 | 5 | 21 | 58 | 24 | 4 | 12 |
| GCF_900083835.1 | 100 | 0.52 | *Klebsiella michiganensis* 98.4715 % | Yes | No | 11 | 3 | 5 | 7 | 8 | 11 | 6 | 5 |
| GCF_020117015.1 | 100 | 0.78 | *Klebsiella michiganensis* 98.4746 % | Yes | No | 253 | 3 | 5 | 21 | 58 | 11 | 6 | 12 |
| GCF_022551435.1 | 100 | 0.54 | *Klebsiella michiganensis* 98.4749 % | Yes | No | 536 | 3 | 5 | 7 | 61 | 24 | 6 | 78 |
| GCF_002856965.1 | 100 | 0.56 | *Klebsiella michiganensis* 98.4774 % | Yes | No | 27 | 3 | 5 | 7 | 13 | 17 | 4 | 5 |
| GCF_019050695.1 | 100 | 0.73 | *Klebsiella michiganensis* 98.4797 % | Yes | No | 108 | 3 | 5 | 21 | 20 | 24 | 6 | 30 |
| GCF_000524315.1 | 100 | 0.27 | *Klebsiella michiganensis* 98.4799 % | Yes | No | 85 | 3 | 5 | 21 | 13 | 24 | 6 | 19 |
| GCF_016735005.1 | 100 | 0.51 | *Klebsiella michiganensis* 98.4836 % | Yes | No | 43 | 3 | 5 | 21 | 20 | 11 | 6 | 20 |
| GCF_015721435.1 | 100 | 1.03 | *Klebsiella michiganensis* 98.4866 % | Yes | No | 239 | 3 | 29 | 32 | 29 | 10 | 6 | 67 |
| GCF_020117665.1 | 100 | 0.66 | *Klebsiella michiganensis* 98.4878 % | Yes | No | 85 | 3 | 5 | 21 | 13 | 24 | 6 | 19 |
| GCF_020118455.1 | 100 | 0.37 | *Klebsiella michiganensis* 98.4879 % | Yes | No | 85 | 3 | 5 | 21 | 13 | 24 | 6 | 19 |
| GCF_019428145.1 | 100 | 0.78 | *Klebsiella michiganensis* 98.4895 % | Yes | No | 27 | 3 | 5 | 7 | 13 | 17 | 4 | 5 |
| GCF_010590245.1 | 100 | 0.67 | *Klebsiella michiganensis* 98.4918 % | Yes | No | 43 | 3 | 5 | 21 | 20 | 11 | 6 | 20 |
| GCF_020118655.1 | 100 | 0.82 | *Klebsiella michiganensis* 98.4929 % | Yes | No | 11 | 3 | 5 | 7 | 8 | 11 | 6 | 5 |
| GCF_018439305.1 | 100 | 0.5 | *Klebsiella michiganensis* 98.4935 % | Yes | No | 213 | 3 | 5 | 21 | 13 | 24 | 6 | 61 |
| GCF_902162465.1 | 100 | 0.63 | *Klebsiella michiganensis* 98.494 % | Yes | Yes | 324 | 32 | 5 | 7 | 61 | 24 | 6 | 78 |
| GCF_902162635.1 | 100 | 0.63 | *Klebsiella michiganensis* 98.4966 % | Yes | Yes | 324 | 32 | 5 | 7 | 61 | 24 | 6 | 78 |
| GCF_902162705.1 | 100 | 0.66 | *Klebsiella michiganensis* 98.4977 % | Yes | Yes | 324 | 32 | 5 | 7 | 61 | 24 | 6 | 78 |
| GCF_018439315.1 | 100 | 0.41 | *Klebsiella michiganensis* 98.4986 % | Yes | No | 213 | 3 | 5 | 21 | 13 | 24 | 6 | 61 |
| GCF_016653015.1 | 100 | 1.62 | *Klebsiella michiganensis* 98.5022 % | Yes | No | 213 | 3 | 5 | 21 | 13 | 24 | 6 | 61 |
| GCF_020118035.1 | 100 | 0.64 | *Klebsiella michiganensis* 98.5022 % | Yes | No | 146 | 3 | 5 | 44 | 13 | 24 | 6 | 50 |
| GCF_001753185.1 | 100 | 3.24 | *Klebsiella michiganensis* 98.5045 % | Yes | No | 85 | 3 | 5 | 21 | 13 | 24 | 6 | 19 |
| GCF_010590505.1 | 100 | 3.18 | *Klebsiella michiganensis* 98.5054 % | Yes | No | 11 | 3 | 5 | 7 | 8 | 11 | 6 | 5 |
| GCF_002947505.1 | 100 | 0.74 | *Klebsiella michiganensis* 98.5059 % | Yes | No | 146 | 3 | 5 | 44 | 13 | 24 | 6 | 50 |
| GCF_022543695.1 | 100 | 1.16 | *Klebsiella michiganensis* 98.5072 % | Yes | Yes | 143 | 3 | 8 | 43 | 7 | 20 | 33 | 23 |
| GCF_000567705.1 | 100 | 0.68 | *Klebsiella michiganensis* 98.5093 % | Yes | No | 143 | 3 | 8 | 43 | 7 | 20 | 33 | 23 |
| GCF_015721205.1 | 100 | 0.55 | *Klebsiella michiganensis* 98.5149 % | Yes | No | 11 | 3 | 5 | 7 | 8 | 11 | 6 | 5 |
| GCF_015721445.1 | 100 | 0.57 | *Klebsiella michiganensis* 98.5164 % | Yes | No | 239 | 3 | 29 | 32 | 29 | 10 | 6 | 67 |
| GCF_018442325.1 | 100 | 0.48 | *Klebsiella michiganensis* 98.5219 % | Yes | No | 213 | 3 | 5 | 21 | 13 | 24 | 6 | 61 |
| GCF_015721185.1 | 100 | 0.21 | *Klebsiella michiganensis* 98.5288 % | Yes | No | 85 | 3 | 5 | 21 | 13 | 24 | 6 | 19 |
| GCF_900083635.1 | 100 | 0.43 | *Klebsiella michiganensis* 98.5296 % | Yes | No | 85 | 3 | 5 | 21 | 13 | 24 | 6 | 19 |
| GCF_022543855.1 | 100 | 1.27 | *Klebsiella michiganensis* 98.5311 % | Yes | Yes | 143 | 3 | 8 | 43 | 7 | 20 | 33 | 23 |
| GCF_900083805.1 | 100 | 0.52 | *Klebsiella michiganensis* 98.5321 % | Yes | No | 11 | 3 | 5 | 7 | 8 | 11 | 6 | 5 |
| GCF_902162685.1 | 100 | 0.62 | *Klebsiella michiganensis* 98.5325 % | Yes | Yes | 324 | 32 | 5 | 7 | 61 | 24 | 6 | 78 |
| GCF_020117595.1 | 100 | 0.8 | *Klebsiella michiganensis* 98.5346 % | Yes | No | 85 | 3 | 5 | 21 | 13 | 24 | 6 | 19 |
| GCF_018439335.1 | 100 | 0.49 | *Klebsiella michiganensis* 98.5359 % | Yes | No | 213 | 3 | 5 | 21 | 13 | 24 | 6 | 61 |
| GCF_016652965.1 | 100 | 1.5 | *Klebsiella michiganensis* 98.5375 % | Yes | No | 27 | 3 | 5 | 7 | 13 | 17 | 4 | 5 |
| GCF_022543595.1 | 100 | 1.31 | *Klebsiella michiganensis* 98.5377 % | Yes | Yes | 143 | 3 | 8 | 43 | 7 | 20 | 33 | 23 |
| GCF_001030755.1 | 100 | 0.68 | *Klebsiella michiganensis* 98.5378 % | Yes | No | 143 | 3 | 8 | 43 | 7 | 20 | 33 | 23 |
| GCF_002152315.1 | 100 | 0.59 | *Klebsiella michiganensis* 98.5403 % | Yes | No | 143 | 3 | 8 | 43 | 7 | 20 | 33 | 23 |
| GCF_009025755.1 | 100 | 0.82 | *Klebsiella michiganensis* 98.5406 % | Yes | No | 27 | 3 | 5 | 7 | 13 | 17 | 4 | 5 |
| GCF_017810085.1 | 100 | 0.61 | *Klebsiella michiganensis* 98.542 % | Yes | No | 27 | 3 | 5 | 7 | 13 | 17 | 4 | 5 |
| GCF_015550995.1 | 100 | 0.84 | *Klebsiella michiganensis* 98.5465 % | No | No | - | ~28 | 5 | 60 | 13 | 24 | 33 | 12 |
| GCF_018442305.1 | 100 | 0.49 | *Klebsiella michiganensis* 98.5499 % | Yes | No | 213 | 3 | 5 | 21 | 13 | 24 | 6 | 61 |
| GCF_021228855.1 | 100 | 0.48 | *Klebsiella michiganensis* 98.5525 % | No | No | 27 | 3 | 5 | 7 | 13 | 17 | 4 | 5 |
| GCF_020118665.1 | 100 | 0.7 | *Klebsiella michiganensis* 98.5526 % | Yes | No | 43 | 3 | 5 | 21 | 20 | 11 | 6 | 20 |
| GCF_020119055.1 | 100 | 0.88 | *Klebsiella michiganensis* 98.5561 % | Yes | No | 27 | 3 | 5 | 7 | 13 | 17 | 4 | 5 |
| GCF_008693565.1 | 100 | 1.84 | *Klebsiella michiganensis* 98.5588 % | Yes | No | 180 | 3 | 8 | 22 | 42 | 20 | 33 | 23 |
| GCF_902162775.1 | 100 | 0.4 | *Klebsiella michiganensis* 98.5628 % | Yes | Yes | 88 | 3 | 8 | 24 | 33 | 20 | 6 | 23 |
| GCF_018422165.1 | 100 | 1.3 | *Klebsiella michiganensis* 98.5631 % | Yes | No | - | 3 | 8 | 24 | 46 | 53 | 32 | 114? |
| GCF_019378695.1 | 100 | 0.14 | *Klebsiella michiganensis* 98.5679 % | No | No | 550 | 3 | 23 | 7 | 19 | 75 | 6 | 12 |
| GCF_022163245.2 | 100 | 0.9 | *Klebsiella michiganensis* 98.5747 % | Yes | Yes | 265 | 28 | 5 | 60 | 13 | 24 | 33 | 12 |
| GCF_900083565.1 | 100 | 0.31 | *Klebsiella michiganensis* 98.5755 % | Yes | No | 294 | 3 | 8 | 24 | 33 | 20 | 6 | 25 |
| GCF_017639895.1 | 100 | 0.72 | *Klebsiella michiganensis* 98.576 % | No | No | 43 | 3 | 5 | 21 | 20 | 11 | 6 | 20 |
| GCF_001066805.1 | 100 | 1.25 | *Klebsiella michiganensis* 98.5764 % | No | No | 11 | 3 | 5 | 7 | 8 | 11 | 6 | 5 |
| GCF_019803065.1 | 100 | 0.61 | *Klebsiella michiganensis* 98.58 % | Yes | No | 535 | 3 | 5 | 7 | 13 | 24 | 6 | 5 |
| GCF_015721335.1 | 100 | 0.9 | *Klebsiella michiganensis* 98.5806 % | Yes | No | 43 | 3 | 5 | 21 | 20 | 11 | 6 | 20 |
| GCF_020120585.1 | 100 | 0.3 | *Klebsiella michiganensis* 98.5808 % | Yes | No | 85 | 3 | 5 | 21 | 13 | 24 | 6 | 19 |
| GCF_015721425.1 | 100 | 0.27 | *Klebsiella michiganensis* 98.5813 % | Yes | No | 85 | 3 | 5 | 21 | 13 | 24 | 6 | 19 |
| GCF_020118555.1 | 100 | 0.48 | *Klebsiella michiganensis* 98.5824 % | No | No | 11 | 3 | 5 | 7 | 8 | 11 | 6 | 5 |
| GCF_018443125.1 | 100 | 0.3 | *Klebsiella michiganensis* 98.5832 % | Yes | No | 213 | 3 | 5 | 21 | 13 | 24 | 6 | 61 |
| GCF_902159795.1 | 100 | 0.9 | *Klebsiella michiganensis* 98.5861 % | Yes | No | 88 | 3 | 8 | 24 | 33 | 20 | 6 | 23 |
| GCF_020121115.1 | 100 | 0.32 | *Klebsiella michiganensis* 98.5876 % | Yes | No | 85 | 3 | 5 | 21 | 13 | 24 | 6 | 19 |
| GCF_016652995.1 | 100 | 0.78 | *Klebsiella michiganensis* 98.588 % | Yes | No | 27 | 3 | 5 | 7 | 13 | 17 | 4 | 5 |
| GCF_013266825.1 | 100 | 2.13 | *Klebsiella michiganensis* 98.5892 % | Yes | No | 515 | 3 | 5 | 60 | 58 | 24 | 4 | 12 |
| GCF_002072655.1 | 100 | 0.86 | *Klebsiella michiganensis* 98.5914 % | Yes | No | 180 | 3 | 8 | 22 | 42 | 20 | 33 | 23 |
| GCF_001052785.1 | 100 | 0.35 | *Klebsiella michiganensis* 98.5915 % | Yes | No | 581 | 49 | 8 | 73 | 29 | 103 | 19 | 30 |
| GCF_020118415.1 | 100 | 0.69 | *Klebsiella michiganensis* 98.5916 % | Yes | No | 151 | 3 | 5 | 21 | 13 | 20 | 6 | 12 |
| GCF_002265195.1 | 100 | 0.82 | *Klebsiella michiganensis* 98.593 % | Yes | No | 27 | 3 | 5 | 7 | 13 | 17 | 4 | 5 |
| GCF_902162745.1 | 100 | 0.4 | *Klebsiella michiganensis* 98.5969 % | Yes | Yes | 88 | 3 | 8 | 24 | 33 | 20 | 6 | 23 |
| GCF_001052035.1 | 100 | 0.18 | *Klebsiella michiganensis* 98.5983 % | Yes | No | 581 | 49 | 8 | 73 | 29 | 103 | 19 | 30 |
| GCF_002111445.1 | 100 | 0.88 | *Klebsiella michiganensis* 98.6052 % | No | No | - | 3 | 5 | ~40 | 13 | 24 | 6 | 50 |
| GCF_901553745.1 | 100 | 0.46 | *Klebsiella michiganensis* 98.6093 % | Yes | No | - | 3 | 8 | 8 | 7 | 10 | 6 | 61 |
| GCF_013821765.1 | 100 | 0.53 | *Klebsiella michiganensis* 98.6139 % | Yes | No | 510 | 3 | 8 | 24 | 33 | 20 | 6 | 20 |
| GCF_000632415.1 | 100 | 0.52 | *Klebsiella michiganensis* 98.6228 % | No | No | - | 3 | 5 | 60? | 8 | 11 | 6 | 5 |
| GCF_922826575.1 | 100 | 0.56 | *Klebsiella michiganensis* 98.6252 % | Yes | No | 88 | 3 | 8 | 24 | 33 | 20 | 6 | 23 |
| GCF_001317245.2 | 100 | 1.1 | *Klebsiella michiganensis* 98.6268 % | Yes | No | 170 | 3 | 8 | 17 | 42 | 40 | 6 | 23 |
| GCF_022501085.1 | 100 | 0.45 | *Klebsiella michiganensis* 98.6323 % | No | No | 11 | 3 | 5 | 7 | 8 | 11 | 6 | 5 |
| GCF_020117815.1 | 100 | 0.24 | *Klebsiella michiganensis* 98.6351 % | No | No | 540 | 3 | 5 | 40 | 58 | 24 | 4 | 12 |
| GCF_022543625.1 | 100 | 0.91 | *Klebsiella michiganensis* 98.6405 % | Yes | No | 170 | 3 | 8 | 17 | 42 | 40 | 6 | 23 |
| GCF_001938605.1 | 100 | 1.55 | *Klebsiella michiganensis* 98.6416 % | Yes | Yes | 317 | 3 | 8 | 22 | 9 | 20 | 33 | 23 |
| GCF_902159075.1 | 100 | 0.44 | *Klebsiella michiganensis* 98.6479 % | Yes | No | 88 | 3 | 8 | 24 | 33 | 20 | 6 | 23 |
| GCF_019678495.1 | 100 | 0.47 | *Klebsiella michiganensis* 98.6501 % | No | No | 580 | 32 | 5 | 7 | 13 | 24 | 6 | 12 |
| GCF_022359805.1 | 100 | 0.42 | *Klebsiella michiganensis* 98.652 % | Yes | No | 28 | 3 | 5 | 7 | 8 | 11 | 12 | 5 |
| GCF_001053845.1 | 100 | 0.3 | *Klebsiella michiganensis* 98.6526 % | Yes | No | 581 | 49 | 8 | 73 | 29 | 103 | 19 | 30 |
| GCF_022569835.1 | 100 | 0.26 | *Klebsiella michiganensis* 98.653 % | No | No | 85 | 3 | 5 | 21 | 13 | 24 | 6 | 19 |
| GCF_001052045.1 | 100 | 0.2 | *Klebsiella michiganensis* 98.6548 % | Yes | No | 581 | 49 | 8 | 73 | 29 | 103 | 19 | 30 |
| GCF_018140945.1 | 100 | 0.38 | *Klebsiella michiganensis* 98.6558 % | Yes | No | 180 | 3 | 8 | 22 | 42 | 20 | 33 | 23 |
| GCF_900084035.1 | 100 | 0.67 | *Klebsiella michiganensis* 98.6577 % | Yes | No | 88 | 3 | 8 | 24 | 33 | 20 | 6 | 23 |
| GCF_004024205.1 | 100 | 1.26 | *Klebsiella michiganensis* 98.6591 % | Yes | No | 88 | 3 | 8 | 24 | 33 | 20 | 6 | 23 |
| GCF_014103175.1 | 100 | 0.77 | *Klebsiella michiganensis* 98.6598 % | Yes | No | 135 | 3 | 8 | 17 | 45 | 20 | 6 | 48 |
| GCF_001945455.1 | 100 | 1.19 | *Klebsiella michiganensis* 98.6623 % | Yes | No | 50 | 3 | 8 | 8 | 7 | 10 | 6 | 7 |
| GCF_022540105.1 | 100 | 0.93 | *Klebsiella michiganensis* 98.6635 % | Yes | No | 170 | 3 | 8 | 17 | 42 | 40 | 6 | 23 |
| GCF_902166425.1 | 100 | 0.45 | *Klebsiella michiganensis* 98.6661 % | No | No | 88 | 3 | 8 | 24 | 33 | 20 | 6 | 23 |
| GCF_016652915.1 | 100 | 1.38 | *Klebsiella michiganensis* 98.6758 % | Yes | No | 306 | 3 | 8 | 22 | 52 | 20 | 33 | 23 |
| GCF_022543615.1 | 100 | 0.82 | *Klebsiella michiganensis* 98.6816 % | Yes | No | 170 | 3 | 8 | 17 | 42 | 40 | 6 | 23 |
| GCF_022543785.1 | 100 | 0.99 | *Klebsiella michiganensis* 98.682 % | Yes | No | 170 | 3 | 8 | 17 | 42 | 40 | 6 | 23 |
| GCF_014050515.2 | 100 | 0.48 | *Klebsiella michiganensis* 98.686 % | Yes | Yes | 138 | 3 | 8 | 24 | 46 | 53 | 32 | 48 |
| GCF_002852935.1 | 100 | 0.57 | *Klebsiella michiganensis* 98.6893 % | Yes | No | 50 | 3 | 8 | 8 | 7 | 10 | 6 | 7 |
| GCF_000240325.1 | 100 | 0.47 | *Klebsiella michiganensis* 98.6915 % | No | No | 28 | 3 | 5 | 7 | 8 | 11 | 12 | 5 |
| GCF_014104095.1 | 100 | 0.62 | *Klebsiella michiganensis* 98.6921 % | Yes | No | 135 | 3 | 8 | 17 | 45 | 20 | 6 | 48 |
| GCF_902162505.1 | 100 | 1.99 | *Klebsiella michiganensis* 98.6921 % | Yes | No | 381 | 26 | 41 | 40 | 21 | 20 | 6 | 25 |
| GCF_003693455.1 | 100 | 0.81 | *Klebsiella michiganensis* 98.694 % | Yes | No | 98 | 3 | 27 | 32 | 21 | 44 | 19 | 38 |
| GCF_022543675.1 | 100 | 0.96 | *Klebsiella michiganensis* 98.7018 % | Yes | No | 170 | 3 | 8 | 17 | 42 | 40 | 6 | 23 |
| GCF_014050535.2 | 100 | 0.44 | *Klebsiella michiganensis* 98.7021 % | Yes | Yes | 138 | 3 | 8 | 24 | 46 | 53 | 32 | 48 |
| GCF_022543555.1 | 100 | 1.34 | *Klebsiella michiganensis* 98.7034 % | Yes | No | 98 | 3 | 27 | 32 | 21 | 44 | 19 | 38 |
| GCF_022540045.1 | 100 | 1.33 | *Klebsiella michiganensis* 98.7039 % | Yes | No | 98 | 3 | 27 | 32 | 21 | 44 | 19 | 38 |
| GCF_019730535.1 | 100 | 0.27 | *Klebsiella michiganensis* 98.7085 % | Yes | No | 35 | 3 | 8 | 17 | 7 | 20 | 6 | 13 |
| GCF_016734965.1 | 100 | 0.34 | *Klebsiella michiganensis* 98.715 % | Yes | No | 135 | 3 | 8 | 17 | 45 | 20 | 6 | 48 |
| GCF_022539945.1 | 100 | 1.42 | *Klebsiella michiganensis* 98.7159 % | Yes | No | 98 | 3 | 27 | 32 | 21 | 44 | 19 | 38 |
| GCF_902162395.1 | 100 | 1.1 | *Klebsiella michiganensis* 98.7159 % | Yes | No | 381 | 26 | 41 | 40 | 21 | 20 | 6 | 25 |
| GCF_902159515.1 | 100 | 0.75 | *Klebsiella michiganensis* 98.7165 % | Yes | No | 13 | 3 | 9 | 8 | 9 | 13 | 6 | 8 |
| GCF_016653035.1 | 100 | 0.98 | *Klebsiella michiganensis* 98.7171 % | Yes | No | 40 | 3 | 8 | 20 | 7 | 23 | 14 | 18 |
| GCF_014855615.1 | 100 | 2.36 | *Klebsiella michiganensis* 98.7201 % | Yes | No | 381 | 26 | 41 | 40 | 21 | 20 | 6 | 25 |
| GCF_020118815.1 | 100 | 0.31 | *Klebsiella michiganensis* 98.7217 % | Yes | No | 410 | 3 | 55 | 28 | 21 | 103 | 48 | 30 |
| GCF_019730455.1 | 100 | 0.27 | *Klebsiella michiganensis* 98.7251 % | Yes | No | 35 | 3 | 8 | 17 | 7 | 20 | 6 | 13 |
| GCF_020116935.1 | 100 | 0.9 | *Klebsiella michiganensis* 98.7313 % | Yes | No | 547 | 3 | 8 | 24 | 46 | 106 | 6 | 23 |
| GCF_020118495.1 | 100 | 0.28 | *Klebsiella michiganensis* 98.7318 % | Yes | No | 135 | 3 | 8 | 17 | 45 | 20 | 6 | 48 |
| GCF_019678825.1 | 100 | 0.61 | *Klebsiella michiganensis* 98.7353 % | Yes | No | 516 | 44 | 8 | 22 | 7 | 106 | 33 | 23 |
| GCF_007910085.1 | 99.52 | 4.66 | *Klebsiella michiganensis* 98.7357 % | Yes | No | - | 3 | 5 | 7 | 8 | 11 | 12? | 5 |
| GCF_022539965.1 | 100 | 1.55 | *Klebsiella michiganensis* 98.7399 % | Yes | No | 98 | 3 | 27 | 32 | 21 | 44 | 19 | 38 |
| GCF_022543575.1 | 100 | 1.36 | *Klebsiella michiganensis* 98.742 % | Yes | No | 98 | 3 | 27 | 32 | 21 | 44 | 19 | 38 |
| GCF_019730515.1 | 100 | 0.27 | *Klebsiella michiganensis* 98.7424 % | Yes | No | 35 | 3 | 8 | 17 | 7 | 20 | 6 | 13 |
| GCF_902162525.1 | 100 | 1.13 | *Klebsiella michiganensis* 98.7438 % | Yes | No | 381 | 26 | 41 | 40 | 21 | 20 | 6 | 25 |
| GCF_001056405.1 | 100 | 0.53 | *Klebsiella michiganensis* 98.7464 % | Yes | No | - | 12 | 8 | 17 | 9 | 106 | 32 | ~41 |
| GCF_022543495.1 | 100 | 1.54 | *Klebsiella michiganensis* 98.7465 % | Yes | No | 98 | 3 | 27 | 32 | 21 | 44 | 19 | 38 |
| GCF_022543535.1 | 100 | 1.39 | *Klebsiella michiganensis* 98.7509 % | Yes | No | 98 | 3 | 27 | 32 | 21 | 44 | 19 | 38 |
| GCF_014050555.2 | 100 | 0.6 | *Klebsiella michiganensis* 98.751 % | Yes | Yes | 138 | 3 | 8 | 24 | 46 | 53 | 32 | 48 |
| GCF_022543825.1 | 100 | 1.44 | *Klebsiella michiganensis* 98.7525 % | Yes | No | 98 | 3 | 27 | 32 | 21 | 44 | 19 | 38 |
| GCF_022539925.1 | 100 | 0.85 | *Klebsiella michiganensis* 98.7553 % | Yes | No | 170 | 3 | 8 | 17 | 42 | 40 | 6 | 23 |
| GCF_016679895.1 | 100 | 0.59 | *Klebsiella michiganensis* 98.7573 % | Yes | No | 306 | 3 | 8 | 22 | 52 | 20 | 33 | 23 |
| GCF_900083885.1 | 100 | 0.34 | *Klebsiella michiganensis* 98.7578 % | Yes | No | 50 | 3 | 8 | 8 | 7 | 10 | 6 | 7 |
| GCF_018422065.1 | 100 | 0.73 | *Klebsiella michiganensis* 98.7617 % | Yes | No | 180 | 3 | 8 | 22 | 42 | 20 | 33 | 23 |
| GCF_022539905.1 | 100 | 1.47 | *Klebsiella michiganensis* 98.7628 % | Yes | No | 98 | 3 | 27 | 32 | 21 | 44 | 19 | 38 |
| GCF_902164965.1 | 100 | 2.04 | *Klebsiella michiganensis* 98.7716 % | Yes | No | 50 | 3 | 8 | 8 | 7 | 10 | 6 | 7 |
| GCF_017348855.1 | 100 | 0.43 | *Klebsiella michiganensis* 98.7754 % | Yes | No | 370 | 3 | 29 | 74 | 21 | 20 | 31 | 12 |
| GCF_008120305.1 | 100 | 0.53 | *Klebsiella michiganensis* 98.7758 % | Yes | No | - | 3 | 9 | 40 | 21 | 20 | 6 | 46? |
| GCF_001054755.1 | 100 | 0.7 | *Klebsiella michiganensis* 98.7759 % | Yes | No | - | 12 | 8 | 17 | 9 | 106 | 32 | ~41 |
| GCF_902162325.1 | 100 | 1.25 | *Klebsiella michiganensis* 98.7813 % | Yes | No | 393 | 3 | 8 | 20 | 7 | 20 | 14 | 18 |
| GCF_001038305.1 | 100 | 1.11 | *Klebsiella michiganensis* 98.7856 % | Yes | Yes | - | 12 | 8 | 22 | 21 | 40 | ~31 | 27 |
| GCF_022540065.1 | 100 | 1.37 | *Klebsiella michiganensis* 98.7859 % | Yes | No | 98 | 3 | 27 | 32 | 21 | 44 | 19 | 38 |
| GCF_003402095.1 | 100 | 2.87 | *Klebsiella michiganensis* 98.7872 % | Yes | Yes | 226 | 3 | 9 | 55 | 21 | 81 | 6 | 64 |
| GCF_004156315.1 | 100 | 0.27 | *Klebsiella michiganensis* 98.7892 % | Yes | No | 52 | 8 | 8 | 16 | 21 | 20 | 19 | 12 |
| GCF_022539865.1 | 100 | 1.41 | *Klebsiella michiganensis* 98.7896 % | Yes | No | 98 | 3 | 27 | 32 | 21 | 44 | 19 | 38 |
| GCF_013636415.1 | 100 | 0.39 | *Klebsiella michiganensis* 98.7927 % | Yes | No | 545 | 3 | 8 | 24 | 42 | 40 | 6 | 25 |
| GCF_019428105.1 | 100 | 0.53 | *Klebsiella michiganensis* 98.7936 % | Yes | No | 88 | 3 | 8 | 24 | 33 | 20 | 6 | 23 |
| GCF_001051455.1 | 100 | 0.34 | *Klebsiella michiganensis* 98.7956 % | No | No | - | 3 | 8 | 28 | 29 | 103 | 19 | ~30 |
| GCF_015773165.1 | 100 | 0.74 | *Klebsiella michiganensis* 98.7974 % | Yes | No | 98 | 3 | 27 | 32 | 21 | 44 | 19 | 38 |
| GCF_020121435.1 | 100 | 0.48 | *Klebsiella michiganensis* 98.7981 % | Yes | No | 353 | 3 | 14 | 22 | 21 | 20 | 17 | 21 |
| GCF_014105195.1 | 100 | 0.47 | *Klebsiella michiganensis* 98.8023 % | Yes | No | 545 | 3 | 8 | 24 | 42 | 40 | 6 | 25 |
| GCF_900451945.1 | 99.99 | 1.49 | *Klebsiella michiganensis* 98.8049 % | Yes | No | - | 3 | 8 | 24 | 9 | 40 | ~6 | 41? |
| GCF_022543815.1 | 100 | 1.17 | *Klebsiella michiganensis* 98.8074 % | Yes | No | 98 | 3 | 27 | 32 | 21 | 44 | 19 | 38 |
| GCF_023502385.1 | 93.03 | 2.36 | *Klebsiella michiganensis* 98.8085 % | Yes | No | - | - | 8 | 20 | 7 | - | 14 | 18 |
| GCF_922832915.1 | 100 | 0.53 | *Klebsiella michiganensis* 98.8124 % | Yes | No | - | 3 | 41 | 8 | 29 | ~107 | 6 | 45 |
| GCF_020695625.1 | 100 | 0.82 | *Klebsiella michiganensis* 98.8129 % | Yes | No | 226 | 3 | 9 | 55 | 21 | 81 | 6 | 64 |
| GCF_022539885.1 | 100 | 1.27 | *Klebsiella michiganensis* 98.8169 % | Yes | No | 98 | 3 | 27 | 32 | 21 | 44 | 19 | 38 |
| GCF_021460075.1 | 100 | 0.31 | *Klebsiella michiganensis* 98.8212 % | No | No | 180 | 3 | 8 | 22 | 42 | 20 | 33 | 23 |
| GCF_018440985.1 | 100 | 2.66 | *Klebsiella michiganensis* 98.8246 % | Yes | No | 98 | 3 | 27 | 32 | 21 | 44 | 19 | 38 |
| GCF_020116315.1 | 100 | 0.47 | *Klebsiella michiganensis* 98.8276 % | Yes | No | 128 | 3 | 31 | 40 | 21 | 20 | 6 | 46 |
| GCF_003074075.1 | 100 | 0.65 | *Klebsiella michiganensis* 98.8294 % | Yes | No | 52 | 8 | 8 | 16 | 21 | 20 | 19 | 12 |
| GCF_007097185.1 | 100 | 0.19 | *Klebsiella michiganensis* 98.8323 % | No | No | 541 | 3 | 8 | 17 | 7 | 20 | 6 | 23 |
| GCF_010598605.1 | 100 | 0.34 | *Klebsiella michiganensis* 98.8323 % | Yes | No | 50 | 3 | 8 | 8 | 7 | 10 | 6 | 7 |
| GCF_019426465.1 | 100 | 0.45 | *Klebsiella michiganensis* 98.835 % | Yes | No | 516 | 44 | 8 | 22 | 7 | 106 | 33 | 23 |
| GCF_018092635.1 | 100 | 0.47 | *Klebsiella michiganensis* 98.8354 % | Yes | Yes | 226 | 3 | 9 | 55 | 21 | 81 | 6 | 64 |
| GCF_017798285.1 | 100 | 0.36 | *Klebsiella michiganensis* 98.8357 % | No | No | - | 3 | 8 | 24 | 21 | 40 | ~15 | 88 |
| GCF_001006545.1 | 100 | 0.34 | *Klebsiella michiganensis* 98.8364 % | No | No | - | 3 | 8 | 28 | 29 | 103 | 19 | ~30 |
| GCF_902164595.1 | 100 | 0.55 | *Klebsiella michiganensis* 98.8366 % | Yes | No | 40 | 3 | 8 | 20 | 7 | 23 | 14 | 18 |
| GCF_020117545.1 | 99.99 | 1.98 | *Klebsiella michiganensis* 98.8385 % | Yes | No | - | 3 | 62? | 7 | ~20 | 23 | ~16 | 18 |
| GCF_900083915.1 | 100 | 1.84 | *Klebsiella michiganensis* 98.8393 % | Yes | Yes | 50 | 3 | 8 | 8 | 7 | 10 | 6 | 7 |
| GCF_001583485.1 | 100 | 0.21 | *Klebsiella michiganensis* 98.8396 % | Yes | No | - | 51? | 14 | 22 | 21 | 25 | 17 | 21 |
| GCF_019731925.1 | 100 | 1.77 | *Klebsiella michiganensis* 98.8411 % | Yes | No | - | ~3 | 8 | 17 | 52 | 20 | 6 | 30? |
| GCF_001053875.1 | 100 | 0.96 | *Klebsiella michiganensis* 98.8457 % | No | No | 546 | 3 | 8 | 24 | 42 | 106 | 6 | 41 |
| GCF_007097115.1 | 100 | 0.19 | *Klebsiella michiganensis* 98.8485 % | No | No | 541 | 3 | 8 | 17 | 7 | 20 | 6 | 23 |
| GCF_020479805.1 | 100 | 0.86 | *Klebsiella michiganensis* 98.8573 % | Yes | No | 50 | 3 | 8 | 8 | 7 | 10 | 6 | 7 |
| GCF_902158915.1 | 100 | 0.56 | *Klebsiella michiganensis* 98.8575 % | Yes | No | 40 | 3 | 8 | 20 | 7 | 23 | 14 | 18 |
| GCF_007106885.1 | 100 | 1.28 | *Klebsiella michiganensis* 98.8579 % | Yes | No | 44 | 3 | 14 | 22 | 21 | 25 | 17 | 21 |
| GCF_021440665.1 | 100 | 0.84 | *Klebsiella michiganensis* 98.8581 % | Yes | No | 353 | 3 | 14 | 22 | 21 | 20 | 17 | 21 |
| GCF_019730985.1 | 100 | 1.15 | *Klebsiella michiganensis* 98.8585 % | Yes | No | 157 | 3 | 34 | 45 | 48 | 20 | 17 | 47 |
| GCF_020117655.1 | 100 | 0.74 | *Klebsiella michiganensis* 98.8599 % | Yes | No | - | 12 | 8 | 20 | 21 | 10 | 32 | 53? |
| GCF_019048985.1 | 100 | 0.29 | *Klebsiella michiganensis* 98.8633 % | Yes | No | - | ~3 | 8 | 16 | 16 | 20 | 6 | 12 |
| GCF_003011775.1 | 100 | 0.36 | *Klebsiella michiganensis* 98.864 % | Yes | No | 40 | 3 | 8 | 20 | 7 | 23 | 14 | 18 |
| GCF_022540025.1 | 99.81 | 1.41 | *Klebsiella michiganensis* 98.8662 % | Yes | No | 98 | 3 | 27 | 32 | 21 | 44 | 19 | 38 |
| GCF_900451325.1 | 99.94 | 3.45 | *Klebsiella michiganensis* 98.8753 % | Yes | No | - | 3 | 8 | 32 | 9 | ~40 | 6 | 27 |
| GCF_008121175.1 | 100 | 0.78 | *Klebsiella michiganensis* 98.8766 % | Yes | No | 366 | 3 | 8 | 17 | 21 | 51 | 6 | 23 |
| GCF_016734915.1 | 100 | 0.6 | *Klebsiella michiganensis* 98.8774 % | Yes | Yes | 64 | 10 | 8 | 27 | 29 | 32 | 6 | 27 |
| GCF_018092585.1 | 100 | 0.47 | *Klebsiella michiganensis* 98.8777 % | Yes | Yes | 226 | 3 | 9 | 55 | 21 | 81 | 6 | 64 |
| GCF_016652925.1 | 100 | 0.29 | *Klebsiella michiganensis* 98.8785 % | Yes | No | 543 | 3 | 8 | 17 | 42 | 106 | 6 | 23 |
| GCF_018604105.1 | 100 | 1.57 | *Klebsiella michiganensis* 98.8809 % | Yes | No | 109 | 3 | 9 | 8 | 9 | 20 | 6 | 8 |
| GCF_902164905.1 | 100 | 0.37 | *Klebsiella michiganensis* 98.8813 % | Yes | No | 40 | 3 | 8 | 20 | 7 | 23 | 14 | 18 |
| GCF_021228755.1 | 100 | 0.59 | *Klebsiella michiganensis* 98.8829 % | Yes | No | 231 | 3 | 9 | 20 | 21 | 25 | 17 | 56 |
| GCF_020118955.1 | 100 | 0.54 | *Klebsiella michiganensis* 98.8852 % | Yes | No | 226 | 3 | 9 | 55 | 21 | 81 | 6 | 64 |
| GCF_900083575.1 | 100 | 0.4 | *Klebsiella michiganensis* 98.8929 % | Yes | No | - | 3 | 8 | 20 | 7 | 23 | 14 | ~18 |
| GCF_020117515.1 | 100 | 1.19 | *Klebsiella michiganensis* 98.8939 % | Yes | No | 457 | 3 | 9 | 17 | 9 | 113 | 6 | 27 |
| GCF_902160685.1 | 100 | 1.64 | *Klebsiella michiganensis* 98.8945 % | Yes | No | 109 | 3 | 9 | 8 | 9 | 20 | 6 | 8 |
| GCF_021165695.1 | 100 | 3.34 | *Klebsiella michiganensis* 98.895 % | Yes | Yes | 50 | 3 | 8 | 8 | 7 | 10 | 6 | 7 |
| GCF_016772995.1 | 100 | 2.35 | *Klebsiella michiganensis* 98.9011 % | Yes | Yes | 50 | 3 | 8 | 8 | 7 | 10 | 6 | 7 |
| GCF_016653085.1 | 100 | 1.11 | *Klebsiella michiganensis* 98.9012 % | Yes | No | 196 | 3 | 14 | 50 | 21 | 25 | 17 | 21 |
| GCF_002887165.1 | 100 | 2.69 | *Klebsiella michiganensis* 98.9022 % | Yes | No | 50 | 3 | 8 | 8 | 7 | 10 | 6 | 7 |
| GCF_902162475.1 | 100 | 1.66 | *Klebsiella michiganensis* 98.904 % | Yes | No | 157 | 3 | 34 | 45 | 48 | 20 | 17 | 47 |
| GCF_902158845.1 | 100 | 1 | *Klebsiella michiganensis* 98.9061 % | Yes | No | 109 | 3 | 9 | 8 | 9 | 20 | 6 | 8 |
| GCF_902162385.1 | 100 | 1.71 | *Klebsiella michiganensis* 98.9075 % | Yes | No | 157 | 3 | 34 | 45 | 48 | 20 | 17 | 47 |
| GCF_015721285.1 | 100 | 0.31 | *Klebsiella michiganensis* 98.9119 % | Yes | No | 32 | 3 | 8 | 16 | 16 | 20 | 6 | 12 |
| GCF_002119875.1 | 100 | 0.56 | *Klebsiella michiganensis* 98.9146 % | Yes | No | - | 3 | 8 | 8 | 7 | 10 | 6 | ~7 |
| GCF_020116815.1 | 100 | 0.53 | *Klebsiella michiganensis* 98.9157 % | Yes | No | 50 | 3 | 8 | 8 | 7 | 10 | 6 | 7 |
| GCF_902162405.1 | 100 | 1.64 | *Klebsiella michiganensis* 98.9175 % | Yes | No | 157 | 3 | 34 | 45 | 48 | 20 | 17 | 47 |
| GCF_020116955.1 | 100 | 0.39 | *Klebsiella michiganensis* 98.9201 % | Yes | No | 544 | 3 | 8 | 24 | 21 | 20 | 6 | 88 |
| GCF_020117695.1 | 100 | 0.69 | *Klebsiella michiganensis* 98.9209 % | Yes | No | 13 | 3 | 9 | 8 | 9 | 13 | 6 | 8 |
| GCF_023093655.1 | 100 | 2 | *Klebsiella michiganensis* 98.9222 % | Yes | No | 50 | 3 | 8 | 8 | 7 | 10 | 6 | 7 |
| GCF_020120895.1 | 100 | 0.93 | *Klebsiella michiganensis* 98.928 % | Yes | No | 44 | 3 | 14 | 22 | 21 | 25 | 17 | 21 |
| GCF_020118915.1 | 100 | 0.36 | *Klebsiella michiganensis* 98.9307 % | Yes | No | 40 | 3 | 8 | 20 | 7 | 23 | 14 | 18 |
| GCF_902159865.1 | 100 | 1.15 | *Klebsiella michiganensis* 98.9321 % | Yes | No | 50 | 3 | 8 | 8 | 7 | 10 | 6 | 7 |
| GCF_010588125.1 | 100 | 0.24 | *Klebsiella michiganensis* 98.9363 % | Yes | No | - | 3 | ~8 | 8 | 7 | 10 | 6 | 7 |
| GCF_008120085.1 | 100 | 0.9 | *Klebsiella michiganensis* 98.9377 % | Yes | No | 157 | 3 | 34 | 45 | 48 | 20 | 17 | 47 |
| GCF_015721325.1 | 100 | 0.31 | *Klebsiella michiganensis* 98.9409 % | Yes | No | 32 | 3 | 8 | 16 | 16 | 20 | 6 | 12 |
| GCF_019378535.1 | 100 | 0.11 | *Klebsiella michiganensis* 98.9428 % | No | No | 572 | 12 | 8 | 20 | 21 | 10 | 32 | 12 |
| GCF_014330695.1 | 100 | 0.39 | *Klebsiella michiganensis* 98.9433 % | No | No | 575 | 12 | 29 | 20 | 21 | 40 | 32 | 54 |
| GCF_902159905.1 | 100 | 1.01 | *Klebsiella michiganensis* 98.9444 % | Yes | No | 50 | 3 | 8 | 8 | 7 | 10 | 6 | 7 |
| GCF_010590665.1 | 100 | 1.19 | *Klebsiella michiganensis* 98.9453 % | Yes | No | 32 | 3 | 8 | 16 | 16 | 20 | 6 | 12 |
| GCF_019754135.1 | 100 | 0.42 | *Klebsiella michiganensis* 98.9621 % | Yes | No | 203 | 3 | 39 | 8 | 7 | 20 | 6 | 38 |
| GCF_020118055.1 | 100 | 0.57 | *Klebsiella michiganensis* 98.9639 % | Yes | No | 231 | 3 | 9 | 20 | 21 | 25 | 17 | 56 |
| GCF_020117095.1 | 100 | 0.25 | *Klebsiella michiganensis* 98.9751 % | Yes | No | 40 | 3 | 8 | 20 | 7 | 23 | 14 | 18 |
| GCF_902159815.1 | 100 | 0.84 | *Klebsiella michiganensis* 98.9778 % | Yes | No | - | 3 | 14 | ~22 | 21 | 25 | 17 | 21 |
| GCF_902159775.1 | 100 | 0.84 | *Klebsiella michiganensis* 98.9791 % | Yes | No | - | 3 | 14 | ~22 | 21 | 25 | 17 | 21 |
| GCF_022501105.1 | 100 | 0.25 | *Klebsiella michiganensis* 98.9811 % | Yes | No | 194 | 12 | 29 | 20 | 51 | 40 | 6 | 46 |
| GCF_020117955.1 | 100 | 0.78 | *Klebsiella michiganensis* 98.9814 % | Yes | No | 44 | 3 | 14 | 22 | 21 | 25 | 17 | 21 |
| GCF_021228995.1 | 100 | 0.31 | *Klebsiella michiganensis* 98.9881 % | Yes | No | 196 | 3 | 14 | 50 | 21 | 25 | 17 | 21 |
| GCF_922831305.1 | 100 | 0.31 | *Klebsiella michiganensis* 99.0009 % | Yes | No | 194 | 12 | 29 | 20 | 51 | 40 | 6 | 46 |
| GCF_016734985.1 | 100 | 1.88 | *Klebsiella michiganensis* 99.0014 % | Yes | No | 231 | 3 | 9 | 20 | 21 | 25 | 17 | 56 |
| GCF_015679365.1 | 100 | 0.71 | *Klebsiella michiganensis* 99.0017 % | Yes | No | 303 | 12 | 29 | 20 | 21 | 40 | 32 | 46 |
| GCF_020121285.1 | 100 | 0.71 | *Klebsiella michiganensis* 99.0045 % | Yes | No | 303 | 12 | 29 | 20 | 21 | 40 | 32 | 46 |
| GCF_902162445.1 | 100 | 0.37 | *Klebsiella michiganensis* 99.0063 % | Yes | No | 44 | 3 | 14 | 22 | 21 | 25 | 17 | 21 |
| GCF_013074375.1 | 100 | 0.45 | *Klebsiella michiganensis* 99.0067 % | Yes | No | 573 | 12 | 8 | 22 | 21 | 40 | 31 | 27 |
| GCF_020115525.1 | 100 | 0.43 | *Klebsiella michiganensis* 99.0069 % | No | No | 52 | 8 | 8 | 16 | 21 | 20 | 19 | 12 |
| GCF_019754075.1 | 100 | 0.42 | *Klebsiella michiganensis* 99.0133 % | Yes | No | 203 | 3 | 39 | 8 | 7 | 20 | 6 | 38 |
| GCF_001056445.1 | 100 | 0.79 | *Klebsiella michiganensis* 99.0164 % | Yes | No | 41 | 3 | 9 | 8 | 9 | 20 | 15 | 8 |
| GCF_020117535.1 | 100 | 0.32 | *Klebsiella michiganensis* 99.0223 % | Yes | No | 417 | 12 | 9 | 17 | 9 | 20 | 6 | 30 |
| GCF_902166555.1 | 99.93 | 0.2 | *Klebsiella michiganensis* 99.0236 % | No | No | 44 | 3 | 14 | 22 | 21 | 25 | 17 | 21 |
| GCF_902166795.1 | 100 | 0.45 | *Klebsiella michiganensis* 99.0242 % | No | No | 40 | 3 | 8 | 20 | 7 | 23 | 14 | 18 |
| GCF_000633235.1 | 100 | 0.42 | *Klebsiella michiganensis* 99.0247 % | No | No | 52 | 8 | 8 | 16 | 21 | 20 | 19 | 12 |
| GCF_020118365.1 | 100 | 0.23 | *Klebsiella michiganensis* 99.0346 % | Yes | No | 231 | 3 | 9 | 20 | 21 | 25 | 17 | 56 |
| GCF_021394645.1 | 100 | 0.27 | *Klebsiella michiganensis* 99.0436 % | No | No | 50 | 3 | 8 | 8 | 7 | 10 | 6 | 7 |
| GCF_902164605.1 | 100 | 0.4 | *Klebsiella michiganensis* 99.0443 % | Yes | No | 40 | 3 | 8 | 20 | 7 | 23 | 14 | 18 |
| GCF_020117075.1 | 100 | 0.31 | *Klebsiella michiganensis* 99.0501 % | Yes | No | - | 12 | 29 | 55 | 12 | 20 | 31 | ~114 |
| GCF_900083825.1 | 100 | 0.43 | *Klebsiella michiganensis* 99.0504 % | No | No | - | 3 | 8 | 32 | 21 | ~103 | 6 | 47 |
| GCF_022859475.1 | 100 | 0.24 | *Klebsiella michiganensis* 99.0552 % | No | No | 194 | 12 | 29 | 20 | 51 | 40 | 6 | 46 |
| GCF_015139575.1 | 100 | 0.33 | *Klebsiella michiganensis* 99.0561 % | Yes | No | - | 12 | 29 | 17 | 9 | 20 | 57 | ~114 |
| GCF_020117465.1 | 100 | 0.49 | *Klebsiella michiganensis* 99.0629 % | Yes | No | 552 | 3 | 29 | 8 | 42 | 108 | 6 | 45 |
| GCF_014217275.1 | 100 | 0.29 | *Klebsiella michiganensis* 99.0689 % | No | No | 456 | 3 | 9 | 40 | 21 | 13 | 6 | 46 |
| GCF_900407255.1 | 100 | 1.44 | *Klebsiella michiganensis* 99.0708 % | Yes | No | - | 3 | ~8 | ~74 | 12 | 20 | 17 | 38 |
| GCF_020118235.1 | 100 | 0.27 | *Klebsiella michiganensis* 99.0724 % | Yes | No | 231 | 3 | 9 | 20 | 21 | 25 | 17 | 56 |
| GCF_020115135.1 | 100 | 0.58 | *Klebsiella michiganensis* 99.0757 % | No | No | 585 | 12 | 59 | 17 | 21 | 20 | 19 | 7 |
| GCF_902166625.1 | 99.98 | 0.28 | *Klebsiella michiganensis* 99.0798 % | No | No | 360 | 3 | 29 | 8 | 21 | 20 | 6 | 48 |
| GCF_902164365.1 | 100 | 0.22 | *Klebsiella michiganensis* 99.1088 % | Yes | No | 50 | 3 | 8 | 8 | 7 | 10 | 6 | 7 |
| GCF_002887605.1 | 100 | 0.4 | *Klebsiella michiganensis* 99.3221 % | Yes | No | - | 12 | 9 | 17 | 9 | 20 | ~17 | 29 |
| GCF_015721485.1 | 100 | 0.28 | *Klebsiella michiganensis* 99.3724 % | Yes | No | 205 | 12 | 9 | 17 | 9 | 20 | 17 | 29 |
| GCF_019679175.1 | 100 | 0.37 | *Klebsiella michiganensis* 99.3999 % | Yes | No | 205 | 12 | 9 | 17 | 9 | 20 | 17 | 29 |
| GCF_018439355.1 | 100 | 0.41 | *Klebsiella michiganensis* 99.4002 % | Yes | No | 205 | 12 | 9 | 17 | 9 | 20 | 17 | 29 |
| GCF_018438865.1 | 100 | 0.25 | *Klebsiella michiganensis* 99.4317 % | Yes | No | 205 | 12 | 9 | 17 | 9 | 20 | 17 | 29 |
| GCF_018439405.1 | 100 | 0.24 | *Klebsiella michiganensis* 99.4505 % | Yes | No | 205 | 12 | 9 | 17 | 9 | 20 | 17 | 29 |
| GCF_019378655.1 | 100 | 0.23 | *Klebsiella michiganensis* 99.4544 % | No | No | 68 | 12 | 8 | 17 | 21 | 20 | 17 | 29 |
| GCF_902163395.1 | 100 | 4.86 | *Klebsiella michiganensis* 99.4817 % | Yes | No | 177 | 3 | 8 | 17 | 21 | 40 | 17 | 54 |
| GCF_017114595.1 | 100 | 2.94 | *Klebsiella michiganensis* 99.5719 % | Yes | No | 84 | 3 | 8 | 17 | 21 | 40 | 17 | 29 |
| GCF_002918665.1 | 100 | 0.79 | *Klebsiella michiganensis* 99.6072 % | Yes | Yes | 84 | 3 | 8 | 17 | 21 | 40 | 17 | 29 |
| GCF_015721625.1 | 100 | 0.44 | *Klebsiella michiganensis* 99.6563 % | Yes | No | 84 | 3 | 8 | 17 | 21 | 40 | 17 | 29 |
| GCF_019678415.1 | 100 | 0.27 | *Klebsiella michiganensis* 99.6679 % | Yes | No | 84 | 3 | 8 | 17 | 21 | 40 | 17 | 29 |
| GCF_015721645.1 | 100 | 0.49 | *Klebsiella michiganensis* 99.6805 % | Yes | No | 84 | 3 | 8 | 17 | 21 | 40 | 17 | 29 |
| GCF_902166585.1 | 100 | 0.18 | *Klebsiella michiganensis* 99.6913 % | No | No | 177 | 3 | 8 | 17 | 21 | 40 | 17 | 54 |
| GCF_015721585.1 | 100 | 0.5 | *Klebsiella michiganensis* 99.7082 % | Yes | No | 84 | 3 | 8 | 17 | 21 | 40 | 17 | 29 |
| GCF_009173485.1 | 100 | 0.75 | *Klebsiella michiganensis* 99.9865 % | Yes | No | 84 | 3 | 8 | 17 | 21 | 40 | 17 | 29 |
| GCF_900977765.1 | 100 | 0.51 | *Klebsiella oxytoca* 100 % | Yes | No | 199 | 2 | 2 | 2 | 3 | 19 | 2 | 2 |
| GCF_013826985.1 | 100 | 0.35 | *Klebsiella oxytoca* 98.8583 % | Yes | Yes | 145 | 1 | 3 | 2 | 34 | 16 | 34 | 1 |
| GCF_002265085.1 | 100 | 1.25 | *Klebsiella oxytoca* 98.8742 % | Yes | No | 145 | 1 | 3 | 2 | 34 | 16 | 34 | 1 |
| GCF_009648375.1 | 100 | 1.94 | *Klebsiella oxytoca* 98.8955 % | Yes | No | 145 | 1 | 3 | 2 | 34 | 16 | 34 | 1 |
| GCF_002853215.1 | 100 | 1.36 | *Klebsiella oxytoca* 98.8984 % | Yes | No | 145 | 1 | 3 | 2 | 34 | 16 | 34 | 1 |
| GCF_900478285.1 | 94.91 | 2.53 | *Klebsiella oxytoca* 98.9231 % | No | No | 101 | 2 | 3 | 2 | 2 | 45 | 10 | 1 |
| GCF_905322525.2 | 100 | 0.46 | *Klebsiella oxytoca* 98.927 % | Yes | No | 145 | 1 | 3 | 2 | 34 | 16 | 34 | 1 |
| GCF_009648435.1 | 100 | 1.91 | *Klebsiella oxytoca* 98.9295 % | Yes | No | 145 | 1 | 3 | 2 | 34 | 16 | 34 | 1 |
| GCF_001594375.1 | 100 | 0.4 | *Klebsiella oxytoca* 98.9353 % | No | No | 145 | 1 | 3 | 2 | 34 | 16 | 34 | 1 |
| GCF_018441525.1 | 100 | 0.3 | *Klebsiella oxytoca* 98.9406 % | Yes | No | 222 | 23 | 42 | 2 | 14 | 34 | 41 | 62 |
| GCF_002853195.1 | 100 | 1.36 | *Klebsiella oxytoca* 98.9411 % | Yes | No | 145 | 1 | 3 | 2 | 34 | 16 | 34 | 1 |
| GCF_003073975.1 | 100 | 0.76 | *Klebsiella oxytoca* 98.9418 % | Yes | No | 600 | 1 | 3 | 13 | 24 | 127 | 10 | 10 |
| GCF_002984395.1 | 100 | 0.35 | *Klebsiella oxytoca* 98.9486 % | Yes | No | 222 | 23 | 42 | 2 | 14 | 34 | 41 | 62 |
| GCF_900451165.1 | 100 | 0.5 | *Klebsiella oxytoca* 98.9776 % | No | No | 257 | 1 | 3 | 13 | 24 | 2 | 10 | 10 |
| GCF_020120515.1 | 100 | 0.27 | *Klebsiella oxytoca* 98.9825 % | Yes | No | - | 1 | 3 | 13 | 1 | 127 | 10 | 10 |
| GCF_010365605.1 | 100 | 0.35 | *Klebsiella oxytoca* 98.9852 % | Yes | No | 222 | 23 | 42 | 2 | 14 | 34 | 41 | 62 |
| GCF_018444115.1 | 100 | 0.28 | *Klebsiella oxytoca* 98.9913 % | Yes | No | - | 1 | 3 | 13 | 24 | 2 | 10 | ~1 |
| GCF_905330835.2 | 100 | 0.24 | *Klebsiella oxytoca* 98.9943 % | Yes | No | 101 | 2 | 3 | 2 | 2 | 45 | 10 | 1 |
| GCF_900635105.1 | 100 | 0.32 | *Klebsiella oxytoca* 98.9945 % | No | No | 101 | 2 | 3 | 2 | 2 | 45 | 10 | 1 |
| GCF_905330865.2 | 100 | 0.24 | *Klebsiella oxytoca* 98.996 % | Yes | No | 101 | 2 | 3 | 2 | 2 | 45 | 10 | 1 |
| GCF_021373415.1 | 100 | 0.75 | *Klebsiella oxytoca* 99.008 % | Yes | No | - | 1 | ~3 | 2 | 2 | 85 | 10 | 2 |
| GCF_902159785.1 | 100 | 0.5 | *Klebsiella oxytoca* 99.0143 % | Yes | No | 397 | 1 | 3 | 13 | 1 | 2 | 10 | 1 |
| GCF_020118135.1 | 100 | 0.21 | *Klebsiella oxytoca* 99.0167 % | Yes | No | - | 1 | 3 | 13 | 24 | 127 | 10 | 1 |
| GCF_016516145.1 | 100 | 1.61 | *Klebsiella oxytoca* 99.0187 % | Yes | No | 592 | 1 | 3 | 13 | 86 | 127 | 10 | 1 |
| GCF_017968865.1 | 100 | 0.98 | *Klebsiella oxytoca* 99.0206 % | No | No | 48 | 1 | 3 | 13 | 24 | 27 | 10 | 1 |
| GCF_015721495.1 | 99.78 | 0.38 | *Klebsiella oxytoca* 99.0221 % | Yes | No | 285 | 1 | 3 | 13 | 24 | 2 | 10 | 1 |
| GCF_004785705.1 | 100 | 0.73 | *Klebsiella oxytoca* 99.0416 % | Yes | No | 48 | 1 | 3 | 13 | 24 | 27 | 10 | 1 |
| GCF_901212425.1 | 100 | 1.72 | *Klebsiella oxytoca* 99.0433 % | Yes | No | - | 1 | 3 | 13 | 1 | 127 | 10 | 1 |
| GCF_902163545.1 | 100 | 0.6 | *Klebsiella oxytoca* 99.0496 % | Yes | No | 397 | 1 | 3 | 13 | 1 | 2 | 10 | 1 |
| GCF_900451235.1 | 99.99 | 0.88 | *Klebsiella oxytoca* 99.0513 % | Yes | No | 287 | 1 | 2 | 2 | 2 | 94 | 1 | 1 |
| GCF_900083975.1 | 100 | 1.44 | *Klebsiella oxytoca* 99.0559 % | No | No | 153 | 1 | 2 | 1 | 1 | 1 | 1 | 1 |
| GCF_020118195.1 | 100 | 0.3 | *Klebsiella oxytoca* 99.0571 % | Yes | No | 529 | 2 | 3 | 2 | 2 | 85 | 10 | 1 |
| GCF_905338025.2 | 100 | 0.19 | *Klebsiella oxytoca* 99.0614 % | Yes | No | 328 | 1 | 3 | 13 | 24 | 97 | 10 | 80 |
| GCF_020116885.1 | 100 | 0.82 | *Klebsiella oxytoca* 99.065 % | Yes | No | 532 | 2 | 19 | 2 | 14 | 29 | 10 | 1 |
| GCF_018068785.1 | 100 | 2.63 | *Klebsiella oxytoca* 99.0654 % | Yes | No | 36 | 1 | 7 | 2 | 15 | 21 | 1 | 14 |
| GCF_009832375.1 | 100 | 1.98 | *Klebsiella oxytoca* 99.0659 % | Yes | No | 528 | 2 | 2 | 1 | 1 | 16 | 1 | 2 |
| GCF_020116675.1 | 100 | 1.01 | *Klebsiella oxytoca* 99.0663 % | Yes | No | 464 | 1 | 3 | 12 | 53 | 16 | 53 | 1 |
| GCF_016903415.1 | 100 | 3.85 | *Klebsiella oxytoca* 99.0689 % | Yes | No | 201 | 1 | 3 | 12 | 53 | 16 | 1 | 1 |
| GCF_900084025.1 | 100 | 1.4 | *Klebsiella oxytoca* 99.0739 % | Yes | No | 36 | 1 | 7 | 2 | 15 | 21 | 1 | 14 |
| GCF_902363365.1 | 100 | 0.25 | *Klebsiella oxytoca* 99.0743 % | Yes | No | 287 | 1 | 2 | 2 | 2 | 94 | 1 | 1 |
| GCF_900083875.1 | 100 | 1.42 | *Klebsiella oxytoca* 99.0745 % | Yes | No | 36 | 1 | 7 | 2 | 15 | 21 | 1 | 14 |
| GCF_900083945.1 | 100 | 1.36 | *Klebsiella oxytoca* 99.0869 % | Yes | No | 36 | 1 | 7 | 2 | 15 | 21 | 1 | 14 |
| GCF_905232115.1 | 100 | 0.13 | *Klebsiella oxytoca* 99.0929 % | Yes | No | 325 | 1 | 7 | 67 | 1 | 4 | 1 | 2 |
| GCF_900083905.1 | 100 | 0.48 | *Klebsiella oxytoca* 99.0949 % | Yes | No | 36 | 1 | 7 | 2 | 15 | 21 | 1 | 14 |
| GCF_902161925.1 | 100 | 1.1 | *Klebsiella oxytoca* 99.0982 % | Yes | No | 36 | 1 | 7 | 2 | 15 | 21 | 1 | 14 |
| GCF_016636165.1 | 100 | 0.29 | *Klebsiella oxytoca* 99.0995 % | Yes | No | 282 | 2 | 19 | 2 | 14 | 29 | 1 | 52 |
| GCF_010590325.1 | 100 | 0.34 | *Klebsiella oxytoca* 99.1014 % | Yes | No | 282 | 2 | 19 | 2 | 14 | 29 | 1 | 52 |
| GCF_016734995.1 | 99.96 | 0.21 | *Klebsiella oxytoca* 99.1021 % | No | No | 257 | 1 | 3 | 13 | 24 | 2 | 10 | 10 |
| GCF_902166465.1 | 100 | 0.45 | *Klebsiella oxytoca* 99.1033 % | No | No | 36 | 1 | 7 | 2 | 15 | 21 | 1 | 14 |
| GCF_905331245.1 | 100 | 0.99 | *Klebsiella oxytoca* 99.108 % | Yes | No | 36 | 1 | 7 | 2 | 15 | 21 | 1 | 14 |
| GCF_902705735.1 | 100 | 2.12 | *Klebsiella oxytoca* 99.115 % | Yes | No | 1 | 1 | 1 | 1 | 1 | 1 | 1 | 1 |
| GCF_000247875.1 | 100 | 0.97 | *Klebsiella oxytoca* 99.1254 % | Yes | No | 31 | 1 | 13 | 1 | 15 | 4 | 1 | 1 |
| GCF_020116575.1 | 100 | 0.32 | *Klebsiella oxytoca* 99.1324 % | Yes | No | 101 | 2 | 3 | 2 | 2 | 45 | 10 | 1 |
| GCF_020118215.1 | 100 | 1.93 | *Klebsiella oxytoca* 99.1333 % | Yes | No | 1 | 1 | 1 | 1 | 1 | 1 | 1 | 1 |
| GCF_020116275.1 | 100 | 0.25 | *Klebsiella oxytoca* 99.1386 % | Yes | No | 46 | 1 | 7 | 2 | 15 | 2 | 1 | 1 |
| GCF_018447075.1 | 100 | 1.2 | *Klebsiella oxytoca* 99.14 % | Yes | No | - | 1 | ~7 | 2 | 15 | 21 | 1 | ~14 |
| GCF_020119125.1 | 100 | 0.28 | *Klebsiella oxytoca* 99.1453 % | Yes | No | 46 | 1 | 7 | 2 | 15 | 2 | 1 | 1 |
| GCF_000252915.2 | 100 | 0.33 | *Klebsiella oxytoca* 99.1501 % | Yes | No | 36 | 1 | 7 | 2 | 15 | 21 | 1 | 14 |
| GCF_010590585.1 | 100 | 0.3 | *Klebsiella oxytoca* 99.155 % | Yes | No | 237 | 1 | 13 | 1 | 56 | 4 | 1 | 1 |
| GCF_905333885.2 | 100 | 1.59 | *Klebsiella oxytoca* 99.157 % | Yes | No | 36 | 1 | 7 | 2 | 15 | 21 | 1 | 14 |
| GCF_020116835.1 | 100 | 0.25 | *Klebsiella oxytoca* 99.1579 % | Yes | No | 46 | 1 | 7 | 2 | 15 | 2 | 1 | 1 |
| GCF_929608445.1 | 100 | 2.06 | *Klebsiella oxytoca* 99.1591 % | Yes | No | 530 | 2 | 7 | 2 | 15 | 19 | 1 | 1 |
| GCF_013750595.1 | 100 | 0.57 | *Klebsiella oxytoca* 99.1598 % | Yes | No | 37 | 1 | 7 | 1 | 18 | 4 | 13 | 15 |
| GCF_015679305.1 | 100 | 1.36 | *Klebsiella oxytoca* 99.1598 % | Yes | No | 527 | 1 | 17 | 1 | 15 | 91 | 1 | 2 |
| GCF_008082015.1 | 100 | 0.26 | *Klebsiella oxytoca* 99.1647 % | Yes | No | 500 | 2 | 7 | 2 | 15 | 2 | 56 | 1 |
| GCF_900083625.1 | 100 | 0.39 | *Klebsiella oxytoca* 99.166 % | Yes | No | 141 | 1 | 7 | 2 | 1 | 4 | 1 | 2 |
| GCF_016636055.1 | 100 | 0.67 | *Klebsiella oxytoca* 99.1665 % | Yes | No | 65 | 1 | 7 | 2 | 30 | 4 | 21 | 2 |
| GCF_016774035.1 | 100 | 2.37 | *Klebsiella oxytoca* 99.1688 % | Yes | Yes | 278 | 1 | 2 | 2 | 60 | 91 | 1 | 76 |
| GCF_902158835.1 | 100 | 0.65 | *Klebsiella oxytoca* 99.1812 % | Yes | No | 37 | 1 | 7 | 1 | 18 | 4 | 13 | 15 |
| GCF_016774075.1 | 100 | 2.38 | *Klebsiella oxytoca* 99.1849 % | Yes | Yes | 278 | 1 | 2 | 2 | 60 | 91 | 1 | 76 |
| GCF_900083585.1 | 100 | 1.19 | *Klebsiella oxytoca* 99.1869 % | Yes | No | 2 | 1 | 2 | 2 | 1 | 2 | 1 | 2 |
| GCF_902705745.1 | 100 | 1.23 | *Klebsiella oxytoca* 99.1869 % | Yes | No | 258 | 2 | 7 | 2 | 15 | 2 | 1 | 1 |
| GCF_015721705.1 | 100 | 0.35 | *Klebsiella oxytoca* 99.187 % | Yes | No | 200 | 2 | 7 | 2 | 15 | 2 | 1 | 57 |
| GCF_902160885.1 | 100 | 1.09 | *Klebsiella oxytoca* 99.1874 % | Yes | No | - | 1 | ~11 | 12 | 1 | 4 | 9 | 1 |
| GCF_001078195.1 | 100 | 0.25 | *Klebsiella oxytoca* 99.1887 % | Yes | No | 258 | 2 | 7 | 2 | 15 | 2 | 1 | 1 |
| GCF_016773035.1 | 100 | 2.37 | *Klebsiella oxytoca* 99.1901 % | Yes | Yes | 278 | 1 | 2 | 2 | 60 | 91 | 1 | 76 |
| GCF_020118785.1 | 100 | 0.41 | *Klebsiella oxytoca* 99.1925 % | Yes | No | 258 | 2 | 7 | 2 | 15 | 2 | 1 | 1 |
| GCF_902159605.1 | 100 | 0.6 | *Klebsiella oxytoca* 99.1979 % | Yes | No | 37 | 1 | 7 | 1 | 18 | 4 | 13 | 15 |
| GCF_001065715.1 | 100 | 2.26 | *Klebsiella oxytoca* 99.1989 % | Yes | No | 58 | 1 | 2 | 2 | 1 | 4 | 1 | 2 |
| GCF_922832405.1 | 100 | 0.56 | *Klebsiella oxytoca* 99.2003 % | Yes | No | 2 | 1 | 2 | 2 | 1 | 2 | 1 | 2 |
| GCF_020119115.1 | 100 | 0.28 | *Klebsiella oxytoca* 99.2017 % | Yes | No | 46 | 1 | 7 | 2 | 15 | 2 | 1 | 1 |
| GCF_020116615.1 | 100 | 1.15 | *Klebsiella oxytoca* 99.203 % | Yes | No | 527 | 1 | 17 | 1 | 15 | 91 | 1 | 2 |
| GCF_020118315.1 | 100 | 0.58 | *Klebsiella oxytoca* 99.2035 % | Yes | No | 37 | 1 | 7 | 1 | 18 | 4 | 13 | 15 |
| GCF_902160925.1 | 100 | 1.09 | *Klebsiella oxytoca* 99.2047 % | Yes | No | - | 1 | ~11 | 12 | 1 | 4 | 9 | 1 |
| GCF_020117035.1 | 100 | 0.45 | *Klebsiella oxytoca* 99.2062 % | Yes | No | 46 | 1 | 7 | 2 | 15 | 2 | 1 | 1 |
| GCF_020116545.1 | 100 | 0.61 | *Klebsiella oxytoca* 99.2064 % | Yes | No | 37 | 1 | 7 | 1 | 18 | 4 | 13 | 15 |
| GCF_023572405.1 | 100 | 0.91 | *Klebsiella oxytoca* 99.2074 % | Yes | Yes | 500 | 2 | 7 | 2 | 15 | 2 | 56 | 1 |
| GCF_020117275.1 | 100 | 0.35 | *Klebsiella oxytoca* 99.2075 % | Yes | No | 58 | 1 | 2 | 2 | 1 | 4 | 1 | 2 |
| GCF_021228775.1 | 100 | 1.12 | *Klebsiella oxytoca* 99.2108 % | Yes | Yes | 278 | 1 | 2 | 2 | 60 | 91 | 1 | 76 |
| GCF_905329555.2 | 100 | 1.19 | *Klebsiella oxytoca* 99.2114 % | Yes | No | 2 | 1 | 2 | 2 | 1 | 2 | 1 | 2 |
| GCF_902164285.1 | 100 | 0.36 | *Klebsiella oxytoca* 99.2115 % | Yes | No | 258 | 2 | 7 | 2 | 15 | 2 | 1 | 1 |
| GCF_012395905.1 | 100 | 0.49 | *Klebsiella oxytoca* 99.2117 % | Yes | Yes | 58 | 1 | 2 | 2 | 1 | 4 | 1 | 2 |
| GCF_902162875.1 | 100 | 0.56 | *Klebsiella oxytoca* 99.2117 % | Yes | No | 37 | 1 | 7 | 1 | 18 | 4 | 13 | 15 |
| GCF_902162905.1 | 100 | 0.37 | *Klebsiella oxytoca* 99.2138 % | Yes | No | 258 | 2 | 7 | 2 | 15 | 2 | 1 | 1 |
| GCF_902164825.1 | 100 | 0.59 | *Klebsiella oxytoca* 99.2177 % | Yes | No | 37 | 1 | 7 | 1 | 18 | 4 | 13 | 15 |
| GCF_900083955.1 | 100 | 0.71 | *Klebsiella oxytoca* 99.2186 % | Yes | No | - | 1 | 2 | 2 | 55 | ~19 | 1 | 2 |
| GCF_900083785.1 | 100 | 0.85 | *Klebsiella oxytoca* 99.2204 % | Yes | No | 2 | 1 | 2 | 2 | 1 | 2 | 1 | 2 |
| GCF_010598965.1 | 100 | 0.72 | *Klebsiella oxytoca* 99.2208 % | Yes | No | 58 | 1 | 2 | 2 | 1 | 4 | 1 | 2 |
| GCF_900083615.1 | 100 | 1.06 | *Klebsiella oxytoca* 99.2226 % | Yes | No | 2 | 1 | 2 | 2 | 1 | 2 | 1 | 2 |
| GCF_905331355.1 | 100 | 0.57 | *Klebsiella oxytoca* 99.2232 % | Yes | No | 2 | 1 | 2 | 2 | 1 | 2 | 1 | 2 |
| GCF_900084045.1 | 100 | 1.65 | *Klebsiella oxytoca* 99.2245 % | Yes | No | 58 | 1 | 2 | 2 | 1 | 4 | 1 | 2 |
| GCF_016774315.1 | 100 | 0.84 | *Klebsiella oxytoca* 99.2252 % | Yes | Yes | 178 | 1 | 11 | 2 | 1 | 4 | 1 | 2 |
| GCF_018443915.1 | 100 | 0.69 | *Klebsiella oxytoca* 99.2327 % | Yes | Yes | 2 | 1 | 2 | 2 | 1 | 2 | 1 | 2 |
| GCF_902161405.1 | 100 | 1.63 | *Klebsiella oxytoca* 99.2327 % | Yes | No | 58 | 1 | 2 | 2 | 1 | 4 | 1 | 2 |
| GCF_019677765.1 | 100 | 0.42 | *Klebsiella oxytoca* 99.2354 % | Yes | No | 21 | 1 | 12 | 2 | 1 | 2 | 1 | 1 |
| GCF_021228655.1 | 100 | 0.4 | *Klebsiella oxytoca* 99.2354 % | Yes | Yes | 178 | 1 | 11 | 2 | 1 | 4 | 1 | 2 |
| GCF_900083655.1 | 100 | 1.27 | *Klebsiella oxytoca* 99.2374 % | Yes | No | 2 | 1 | 2 | 2 | 1 | 2 | 1 | 2 |
| GCF_004005605.1 | 100 | 0.61 | *Klebsiella oxytoca* 99.2375 % | Yes | No | 221 | 1 | 11 | 2 | 1 | 2 | 40 | 2 |
| GCF_900083985.1 | 100 | 1.14 | *Klebsiella oxytoca* 99.2378 % | Yes | No | 2 | 1 | 2 | 2 | 1 | 2 | 1 | 2 |
| GCF_018443185.1 | 100 | 0.67 | *Klebsiella oxytoca* 99.241 % | Yes | Yes | 2 | 1 | 2 | 2 | 1 | 2 | 1 | 2 |
| GCF_016636085.1 | 100 | 0.67 | *Klebsiella oxytoca* 99.2411 % | Yes | No | 58 | 1 | 2 | 2 | 1 | 4 | 1 | 2 |
| GCF_015721405.1 | 100 | 1.66 | *Klebsiella oxytoca* 99.242 % | Yes | No | 240 | 1 | 18 | 2 | 14 | 2 | 1 | 1 |
| GCF_020117055.1 | 100 | 0.31 | *Klebsiella oxytoca* 99.242 % | Yes | No | 252 | 1 | 7 | 2 | 15 | 89 | 1 | 1 |
| GCF_021228715.1 | 100 | 0.78 | *Klebsiella oxytoca* 99.2423 % | Yes | Yes | 178 | 1 | 11 | 2 | 1 | 4 | 1 | 2 |
| GCF_900083815.1 | 100 | 1.25 | *Klebsiella oxytoca* 99.2426 % | Yes | No | 2 | 1 | 2 | 2 | 1 | 2 | 1 | 2 |
| GCF_900084015.1 | 100 | 1.34 | *Klebsiella oxytoca* 99.2426 % | Yes | No | 2 | 1 | 2 | 2 | 1 | 2 | 1 | 2 |
| GCF_018443995.1 | 100 | 0.63 | *Klebsiella oxytoca* 99.2434 % | Yes | Yes | 2 | 1 | 2 | 2 | 1 | 2 | 1 | 2 |
| GCF_018443495.1 | 100 | 0.62 | *Klebsiella oxytoca* 99.2448 % | Yes | Yes | 2 | 1 | 2 | 2 | 1 | 2 | 1 | 2 |
| GCF_000269585.1 | 100 | 0.27 | *Klebsiella oxytoca* 99.2457 % | Yes | No | 53 | 1 | 7 | 2 | 14 | 19 | 9 | 2 |
| GCF_016773175.1 | 100 | 0.43 | *Klebsiella oxytoca* 99.2463 % | Yes | Yes | 178 | 1 | 11 | 2 | 1 | 4 | 1 | 2 |
| GCF_000492815.1 | 100 | 0.47 | *Klebsiella oxytoca* 99.2467 % | Yes | No | 2 | 1 | 2 | 2 | 1 | 2 | 1 | 2 |
| GCF_900083995.1 | 100 | 1.66 | *Klebsiella oxytoca* 99.2476 % | Yes | No | 2 | 1 | 2 | 2 | 1 | 2 | 1 | 2 |
| GCF_902160765.1 | 100 | 0.27 | *Klebsiella oxytoca* 99.2487 % | Yes | No | 20 | 1 | 11 | 12 | 1 | 4 | 9 | 1 |
| GCF_015559095.1 | 100 | 0.32 | *Klebsiella oxytoca* 99.2503 % | No | No | 441 | 35 | 7 | 12 | 18 | 89 | 1 | 1 |
| GCF_020116875.1 | 100 | 0.43 | *Klebsiella oxytoca* 99.2529 % | Yes | No | 525 | 1 | 7 | 1 | 55 | 4 | 1 | 2 |
| GCF_900083965.1 | 100 | 1.53 | *Klebsiella oxytoca* 99.2536 % | Yes | No | 2 | 1 | 2 | 2 | 1 | 2 | 1 | 2 |
| GCF_001078235.1 | 100 | 0.27 | *Klebsiella oxytoca* 99.2544 % | Yes | No | 53 | 1 | 7 | 2 | 14 | 19 | 9 | 2 |
| GCF_900083725.1 | 100 | 1.22 | *Klebsiella oxytoca* 99.2554 % | Yes | No | 2 | 1 | 2 | 2 | 1 | 2 | 1 | 2 |
| GCF_000492955.1 | 100 | 0.45 | *Klebsiella oxytoca* 99.2563 % | No | No | 18 | 1 | 2 | 2 | 3 | 2 | 1 | 2 |
| GCF_008082295.1 | 100 | 1.02 | *Klebsiella oxytoca* 99.2579 % | Yes | No | - | 1 | 7 | 2 | 14 | 19 | 9 | ~2 |
| GCF_020116975.1 | 100 | 0.44 | *Klebsiella oxytoca* 99.2579 % | Yes | No | 223 | 2 | 18 | 2 | 55 | 79 | 1 | 2 |
| GCF_015721725.1 | 99.95 | 0.36 | *Klebsiella oxytoca* 99.2585 % | Yes | Yes | 2 | 1 | 2 | 2 | 1 | 2 | 1 | 2 |
| GCF_018443705.1 | 100 | 0.68 | *Klebsiella oxytoca* 99.2589 % | Yes | Yes | 2 | 1 | 2 | 2 | 1 | 2 | 1 | 2 |
| GCF_003991285.1 | 100 | 0.41 | *Klebsiella oxytoca* 99.2593 % | Yes | No | 181 | 1 | 7 | 2 | 1 | 67 | 1 | 2 |
| GCF_900083855.1 | 100 | 1.26 | *Klebsiella oxytoca* 99.2603 % | Yes | No | 2 | 1 | 2 | 2 | 1 | 2 | 1 | 2 |
| GCF_900083765.1 | 100 | 1.25 | *Klebsiella oxytoca* 99.2614 % | Yes | No | 2 | 1 | 2 | 2 | 1 | 2 | 1 | 2 |
| GCF_021264605.1 | 100 | 0.45 | *Klebsiella oxytoca* 99.2623 % | Yes | No | 18 | 1 | 2 | 2 | 3 | 2 | 1 | 2 |
| GCF_900083705.1 | 100 | 1.26 | *Klebsiella oxytoca* 99.2635 % | Yes | No | 2 | 1 | 2 | 2 | 1 | 2 | 1 | 2 |
| GCF_018444055.1 | 100 | 0.62 | *Klebsiella oxytoca* 99.2638 % | Yes | Yes | 2 | 1 | 2 | 2 | 1 | 2 | 1 | 2 |
| GCF_900407115.1 | 100 | 0.28 | *Klebsiella oxytoca* 99.2643 % | Yes | No | 520 | 1 | 2 | 2 | 1 | 2 | 9 | 2 |
| GCF_018444465.1 | 100 | 0.62 | *Klebsiella oxytoca* 99.2648 % | Yes | Yes | 2 | 1 | 2 | 2 | 1 | 2 | 1 | 2 |
| GCF_015721655.1 | 100 | 0.24 | *Klebsiella oxytoca* 99.266 % | Yes | No | 2 | 1 | 2 | 2 | 1 | 2 | 1 | 2 |
| GCF_015721685.1 | 100 | 0.24 | *Klebsiella oxytoca* 99.2668 % | Yes | No | 2 | 1 | 2 | 2 | 1 | 2 | 1 | 2 |
| GCF_020120535.1 | 100 | 0.41 | *Klebsiella oxytoca* 99.2686 % | No | No | 2 | 1 | 2 | 2 | 1 | 2 | 1 | 2 |
| GCF_018443785.1 | 100 | 0.67 | *Klebsiella oxytoca* 99.2687 % | Yes | Yes | 2 | 1 | 2 | 2 | 1 | 2 | 1 | 2 |
| GCF_020117455.1 | 100 | 0.34 | *Klebsiella oxytoca* 99.27 % | No | No | 2 | 1 | 2 | 2 | 1 | 2 | 1 | 2 |
| GCF_015721345.1 | 100 | 0.44 | *Klebsiella oxytoca* 99.2702 % | Yes | No | 223 | 2 | 18 | 2 | 55 | 79 | 1 | 2 |
| GCF_002508265.1 | 100 | 0.46 | *Klebsiella oxytoca* 99.2705 % | Yes | No | 415 | 1 | 2 | 2 | 1 | 110 | 1 | 2 |
| GCF_015551825.1 | 100 | 0.28 | *Klebsiella oxytoca* 99.2706 % | Yes | No | - | 1 | ~7 | 2 | 1 | 2 | 2 | 1 |
| GCF_018443665.1 | 100 | 0.67 | *Klebsiella oxytoca* 99.2715 % | Yes | Yes | 2 | 1 | 2 | 2 | 1 | 2 | 1 | 2 |
| GCF_016643945.1 | 100 | 0.41 | *Klebsiella oxytoca* 99.2724 % | Yes | No | 19 | 1 | 2 | 2 | 1 | 16 | 1 | 2 |
| GCF_922832385.1 | 100 | 0.9 | *Klebsiella oxytoca* 99.2737 % | Yes | No | 20 | 1 | 11 | 12 | 1 | 4 | 9 | 1 |
| GCF_015721295.1 | 100 | 1.36 | *Klebsiella oxytoca* 99.2738 % | Yes | No | 2 | 1 | 2 | 2 | 1 | 2 | 1 | 2 |
| GCF_017310465.1 | 100 | 1.47 | *Klebsiella oxytoca* 99.2742 % | Yes | No | 34 | 2 | 2 | 2 | 17 | 2 | 1 | 2 |
| GCF_016903975.1 | 100 | 0.43 | *Klebsiella oxytoca* 99.2766 % | Yes | No | 302 | 1 | 18 | 2 | 1 | 84 | 1 | 1 |
| GCF_010590025.1 | 100 | 0.4 | *Klebsiella oxytoca* 99.2774 % | Yes | No | 223 | 2 | 18 | 2 | 55 | 79 | 1 | 2 |
| GCF_020116915.1 | 100 | 0.6 | *Klebsiella oxytoca* 99.2793 % | Yes | No | 176 | 1 | 7 | 2 | 1 | 65 | 1 | 2 |
| GCF_021398915.1 | 100 | 0.4 | *Klebsiella oxytoca* 99.2817 % | No | No | - | 2 | 2 | 2 | 14 | ~19 | 1 | 1 |
| GCF_900083735.1 | 100 | 0.57 | *Klebsiella oxytoca* 99.2844 % | Yes | No | 176 | 1 | 7 | 2 | 1 | 65 | 1 | 2 |
| GCF_019771925.1 | 100 | 0.88 | *Klebsiella oxytoca* 99.285 % | Yes | Yes | 34 | 2 | 2 | 2 | 17 | 2 | 1 | 2 |
| GCF_900083675.1 | 100 | 0.32 | *Klebsiella oxytoca* 99.2856 % | Yes | No | 375 | 37 | 18 | 2 | 17 | 30 | 1 | 2 |
| GCF_003937225.1 | 100 | 0.96 | *Klebsiella oxytoca* 99.2862 % | Yes | No | 522 | 1 | 2 | 2 | 1 | 89 | 1 | 2 |
| GCF_000527235.1 | 100 | 0.33 | *Klebsiella oxytoca* 99.2883 % | Yes | No | 2 | 1 | 2 | 2 | 1 | 2 | 1 | 2 |
| GCF_001078255.1 | 100 | 1.17 | *Klebsiella oxytoca* 99.2883 % | Yes | No | 2 | 1 | 2 | 2 | 1 | 2 | 1 | 2 |
| GCF_020116635.1 | 100 | 0.21 | *Klebsiella oxytoca* 99.2902 % | No | No | 21 | 1 | 12 | 2 | 1 | 2 | 1 | 1 |
| GCF_016636135.1 | 100 | 1.02 | *Klebsiella oxytoca* 99.2915 % | Yes | No | 266 | 1 | 2 | 2 | 1 | 65 | 1 | 2 |
| GCF_016904515.1 | 100 | 0.7 | *Klebsiella oxytoca* 99.2931 % | Yes | No | 176 | 1 | 7 | 2 | 1 | 65 | 1 | 2 |
| GCF_904863345.1 | 100 | 1.39 | *Klebsiella oxytoca* 99.2933 % | Yes | No | 266 | 1 | 2 | 2 | 1 | 65 | 1 | 2 |
| GCF_902163375.1 | 100 | 0.27 | *Klebsiella oxytoca* 99.2942 % | No | No | 523 | 1 | 2 | 12 | 14 | 19 | 9 | 2 |
| GCF_016636105.1 | 100 | 0.27 | *Klebsiella oxytoca* 99.295 % | Yes | No | 266 | 1 | 2 | 2 | 1 | 65 | 1 | 2 |
| GCF_020118775.1 | 100 | 0.26 | *Klebsiella oxytoca* 99.2996 % | Yes | No | 223 | 2 | 18 | 2 | 55 | 79 | 1 | 2 |
| GCF_900083865.1 | 100 | 0.44 | *Klebsiella oxytoca* 99.3013 % | No | No | 375 | 37 | 18 | 2 | 17 | 30 | 1 | 2 |
| GCF_021228185.1 | 100 | 0.8 | *Klebsiella oxytoca* 99.3015 % | Yes | No | 2 | 1 | 2 | 2 | 1 | 2 | 1 | 2 |
| GCF_018138905.1 | 100 | 1.2 | *Klebsiella oxytoca* 99.3031 % | Yes | Yes | 176 | 1 | 7 | 2 | 1 | 65 | 1 | 2 |
| GCF_016903715.1 | 100 | 0.37 | *Klebsiella oxytoca* 99.3049 % | Yes | No | 176 | 1 | 7 | 2 | 1 | 65 | 1 | 2 |
| GCF_022605305.1 | 100 | 0.63 | *Klebsiella oxytoca* 99.3074 % | Yes | No | 176 | 1 | 7 | 2 | 1 | 65 | 1 | 2 |
| GCF_015721385.1 | 100 | 0.26 | *Klebsiella oxytoca* 99.3079 % | Yes | No | 375 | 37 | 18 | 2 | 17 | 30 | 1 | 2 |
| GCF_020116585.1 | 100 | 0.28 | *Klebsiella oxytoca* 99.3089 % | Yes | Yes | 34 | 2 | 2 | 2 | 17 | 2 | 1 | 2 |
| GCF_000247855.1 | 100 | 1.43 | *Klebsiella oxytoca* 99.3092 % | Yes | No | 30 | 1 | 2 | 2 | 14 | 19 | 9 | 2 |
| GCF_001808475.1 | 100 | 0.48 | *Klebsiella oxytoca* 99.3115 % | Yes | No | 53 | 1 | 7 | 2 | 14 | 19 | 9 | 2 |
| GCF_020117225.1 | 100 | 0.55 | *Klebsiella oxytoca* 99.3125 % | Yes | No | 176 | 1 | 7 | 2 | 1 | 65 | 1 | 2 |
| GCF_900083895.1 | 100 | 3.53 | *Klebsiella oxytoca* 99.3133 % | Yes | Yes | 375 | 37 | 18 | 2 | 17 | 30 | 1 | 2 |
| GCF_016636025.1 | 100 | 0.98 | *Klebsiella oxytoca* 99.3157 % | Yes | No | 266 | 1 | 2 | 2 | 1 | 65 | 1 | 2 |
| GCF_020118275.1 | 100 | 0.53 | *Klebsiella oxytoca* 99.3176 % | Yes | No | 176 | 1 | 7 | 2 | 1 | 65 | 1 | 2 |
| GCF_015721265.1 | 100 | 0.73 | *Klebsiella oxytoca* 99.3217 % | Yes | Yes | 223 | 2 | 18 | 2 | 55 | 79 | 1 | 2 |
| GCF_015554545.1 | 100 | 0.69 | *Klebsiella oxytoca* 99.3238 % | Yes | No | 176 | 1 | 7 | 2 | 1 | 65 | 1 | 2 |
| GCF_014103895.1 | 100 | 0.58 | *Klebsiella oxytoca* 99.3298 % | Yes | No | 176 | 1 | 7 | 2 | 1 | 65 | 1 | 2 |
| GCF_000527215.1 | 100 | 0.53 | *Klebsiella oxytoca* 99.33 % | Yes | No | 30 | 1 | 2 | 2 | 14 | 19 | 9 | 2 |
| GCF_015721195.1 | 100 | 0.38 | *Klebsiella oxytoca* 99.3327 % | No | No | 302 | 1 | 18 | 2 | 1 | 84 | 1 | 1 |
| GCF_900083795.1 | 100 | 0.5 | *Klebsiella oxytoca* 99.3366 % | Yes | No | 176 | 1 | 7 | 2 | 1 | 65 | 1 | 2 |
| GCF_900451255.1 | 100 | 1.83 | *Klebsiella oxytoca* 99.3374 % | Yes | No | 19 | 1 | 2 | 2 | 1 | 16 | 1 | 2 |
| GCF_020116855.1 | 100 | 0.49 | *Klebsiella oxytoca* 99.3379 % | Yes | No | 176 | 1 | 7 | 2 | 1 | 65 | 1 | 2 |
| GCF_001078175.1 | 100 | 0.49 | *Klebsiella oxytoca* 99.3386 % | Yes | No | 176 | 1 | 7 | 2 | 1 | 65 | 1 | 2 |
| GCF_002588345.1 | 100 | 3.21 | *Klebsiella oxytoca* 99.3386 % | Yes | No | 34 | 2 | 2 | 2 | 17 | 2 | 1 | 2 |
| GCF_001030775.1 | 100 | 0.5 | *Klebsiella oxytoca* 99.3493 % | Yes | No | 323 | 1 | 3 | 2 | 14 | 4 | 1 | 2 |
| GCF_900083715.1 | 100 | 0.73 | *Klebsiella oxytoca* 99.3497 % | Yes | No | 176 | 1 | 7 | 2 | 1 | 65 | 1 | 2 |
| GCF_900083745.1 | 100 | 0.49 | *Klebsiella oxytoca* 99.3559 % | Yes | No | 176 | 1 | 7 | 2 | 1 | 65 | 1 | 2 |
| GCF_900083645.1 | 100 | 0.42 | *Klebsiella oxytoca* 99.3564 % | Yes | No | 176 | 1 | 7 | 2 | 1 | 65 | 1 | 2 |
| GCF_003812925.1 | 100 | 0.31 | *Klebsiella oxytoca* 99.3567 % | Yes | No | 59 | 2 | 2 | 2 | 27 | 2 | 1 | 2 |
| GCF_020118355.1 | 100 | 0.33 | *Klebsiella oxytoca* 99.362 % | Yes | No | 323 | 1 | 3 | 2 | 14 | 4 | 1 | 2 |
| GCF_020116385.1 | 100 | 0.5 | *Klebsiella oxytoca* 99.3752 % | Yes | No | - | 1 | 2 | 2 | 55 | 85 | 1 | ~2 |
| GCF_021373355.1 | 100 | 0.55 | *Klebsiella oxytoca* 99.3962 % | No | No | 364 | 1 | 7 | 2 | 2 | 65 | 1 | 2 |
| GCF_017974245.1 | 96.53 | 0.42 | *Klebsiella oxytoca* 99.4127 % | Yes | No | - | 1 | 11 | 2 | 14 | - | 1 | 2 |
| GCF_020118595.1 | 100 | 0.43 | *Klebsiella oxytoca* 99.4165 % | Yes | No | 58 | 1 | 2 | 2 | 1 | 4 | 1 | 2 |
| GCF_016636045.1 | 100 | 0.88 | *Klebsiella oxytoca* 99.4208 % | Yes | No | 267 | 1 | 7 | 2 | 1 | 66 | 1 | 2 |
| GCF_004360035.1 | 100 | 0.89 | *Klebsiella oxytoca* 99.4228 % | No | No | 179 | 1 | 36 | 2 | 1 | 66 | 2 | 2 |
| GCF_001063775.1 | 100 | 1.75 | *Klebsiella oxytoca* 99.4424 % | Yes | No | 58 | 1 | 2 | 2 | 1 | 4 | 1 | 2 |
| GCF_000607265.1 | 100 | 0.61 | *Klebsiella oxytoca* 99.4522 % | No | No | 19 | 1 | 2 | 2 | 1 | 16 | 1 | 2 |
| GCF_019677525.1 | 100 | 0.54 | *Klebsiella oxytoca* 99.4538 % | Yes | No | 19 | 1 | 2 | 2 | 1 | 16 | 1 | 2 |
| GCF_001057405.1 | 100 | 0.45 | *Klebsiella oxytoca* 99.4599 % | Yes | No | 58 | 1 | 2 | 2 | 1 | 4 | 1 | 2 |
| GCF_001053715.1 | 100 | 0.68 | *Klebsiella oxytoca* 99.4696 % | Yes | No | 19 | 1 | 2 | 2 | 1 | 16 | 1 | 2 |
| GCF_001022115.1 | 100 | 1.11 | *Klebsiella oxytoca* 99.4788 % | Yes | No | 199 | 2 | 2 | 2 | 3 | 19 | 2 | 2 |
| GCF_015265865.1 | 100 | 1.11 | *Klebsiella oxytoca* 99.4902 % | Yes | No | 450 | 1 | 11 | 2 | 1 | 65 | 1 | 2 |
| GCF_001022295.1 | 100 | 1.1 | *Klebsiella oxytoca* 99.4903 % | Yes | No | 199 | 2 | 2 | 2 | 3 | 19 | 2 | 2 |
| GCF_015265825.1 | 100 | 1.1 | *Klebsiella oxytoca* 99.4941 % | Yes | No | 450 | 1 | 11 | 2 | 1 | 65 | 1 | 2 |
| GCF_902705645.1 | 100 | 0.49 | *Klebsiella oxytoca* 99.4964 % | Yes | No | 521 | 1 | 2 | 2 | 1 | 16 | 1 | 62 |
| GCF_020116375.1 | 100 | 0.5 | *Klebsiella oxytoca* 99.4976 % | Yes | No | 58 | 1 | 2 | 2 | 1 | 4 | 1 | 2 |
| GCF_001030705.1 | 100 | 0.43 | *Klebsiella oxytoca* 99.5046 % | No | No | 199 | 2 | 2 | 2 | 3 | 19 | 2 | 2 |
| GCF_001870185.1 | 100 | 1.04 | *Klebsiella oxytoca* 99.511 % | Yes | No | 199 | 2 | 2 | 2 | 3 | 19 | 2 | 2 |
| GCF_015721555.1 | 100 | 0.54 | *Klebsiella oxytoca* 99.5136 % | No | No | 19 | 1 | 2 | 2 | 1 | 16 | 1 | 2 |
| GCF_015694145.1 | 97.49 | 0.31 | *Klebsiella oxytoca* 99.514 % | Yes | No | 450 | 1 | 11 | 2 | 1 | 65 | 1 | 2 |
| GCF_015721755.1 | 100 | 1.71 | *Klebsiella oxytoca* 99.5669 % | Yes | No | 199 | 2 | 2 | 2 | 3 | 19 | 2 | 2 |
| GCF_022685985.1 | 100 | 1.05 | *Klebsiella oxytoca* 99.5698 % | Yes | No | 199 | 2 | 2 | 2 | 3 | 19 | 2 | 2 |
| GCF_010598865.1 | 100 | 1.26 | *Klebsiella oxytoca* 99.5836 % | Yes | No | 199 | 2 | 2 | 2 | 3 | 19 | 2 | 2 |
| GCF_902166285.1 | 100 | 0.72 | *Klebsiella oxytoca* 99.596 % | Yes | No | 199 | 2 | 2 | 2 | 3 | 19 | 2 | 2 |
| GCF_001055635.1 | 100 | 1.44 | *Klebsiella oxytoca* 99.6159 % | Yes | No | 199 | 2 | 2 | 2 | 3 | 19 | 2 | 2 |
| GCF_000507385.1 | 100 | 0.49 | *Klebsiella oxytoca* 99.6239 % | No | No | 199 | 2 | 2 | 2 | 3 | 19 | 2 | 2 |
| GCF_001054935.1 | 100 | 0.95 | *Klebsiella oxytoca* 99.6268 % | Yes | No | 199 | 2 | 2 | 2 | 3 | 19 | 2 | 2 |
| GCF_001054575.1 | 100 | 0.8 | *Klebsiella oxytoca* 99.635 % | Yes | No | 199 | 2 | 2 | 2 | 3 | 19 | 2 | 2 |
| GCF_020540705.1 | 100 | 0.34 | *Klebsiella oxytoca* 99.6698 % | Yes | No | 199 | 2 | 2 | 2 | 3 | 19 | 2 | 2 |
| GCF_016529765.1 | 100 | 0.64 | *Klebsiella oxytoca* 99.6914 % | Yes | No | 199 | 2 | 2 | 2 | 3 | 19 | 2 | 2 |
| GCF_020117355.1 | 100 | 0.51 | *Klebsiella oxytoca* 99.8621 % | Yes | No | 199 | 2 | 2 | 2 | 3 | 19 | 2 | 2 |
| GCF_900636985.1 | 100 | 0.52 | *Klebsiella oxytoca* 99.9451 % | No | No | 199 | 2 | 2 | 2 | 3 | 19 | 2 | 2 |
| GCF_001598695.1 | 100 | 0.46 | *Klebsiella oxytoca* 99.9525 % | No | No | 199 | 2 | 2 | 2 | 3 | 19 | 2 | 2 |
| GCF_020115535.1 | 100 | 0.45 | *Klebsiella oxytoca* 99.9645 % | No | No | 199 | 2 | 2 | 2 | 3 | 19 | 2 | 2 |
| GCF_902158725.1 | 100 | 0.68 | *Klebsiella pasteurii* 100 % | Yes | No | 321 | 7 | 16 | 38 | 31 | 26 | 25 | 65 |
| GCF_015721765.1 | 99.98 | 0.6 | *Klebsiella pasteurii* 99.0798 % | Yes | No | 193 | 7 | 23 | 29 | 31 | 72 | 25 | 34 |
| GCF_901563825.1 | 100 | 0.32 | *Klebsiella pasteurii* 99.2094 % | Yes | No | 416 | 13 | 23 | 29 | 31 | 92 | 25 | 34 |
| GCF_019890895.1 | 100 | 0.39 | *Klebsiella pasteurii* 99.2691 % | Yes | No | - | 17 | 16 | 58 | 31 | 92 | 25 | 22 |
| GCF_902158675.1 | 100 | 0.98 | *Klebsiella pasteurii* 99.2796 % | Yes | No | - | ~25 | 23 | 58 | 31 | 82 | 25 | 43 |
| GCF_018423175.1 | 100 | 0.72 | *Klebsiella pasteurii* 99.2966 % | Yes | No | - | 7 | 23 | 58 | 31 | ~92 | 25 | 43 |
| GCF_902158645.1 | 100 | 0.26 | *Klebsiella pasteurii* 99.3145 % | No | No | - | 7 | 23 | ~29 | 31 | 6 | 25 | 34 |
| GCF_015550565.1 | 100 | 0.2 | *Klebsiella pasteurii* 99.3272 % | Yes | No | 185 | 7 | 23 | 23 | 31 | 68 | 25 | 43 |
| GCF_012843205.1 | 100 | 0.48 | *Klebsiella pasteurii* 99.331 % | Yes | No | - | 7 | 23 | 29 | 31 | 69 | 30 | 22 |
| GCF_902158575.1 | 100 | 0.4 | *Klebsiella pasteurii* 99.3335 % | No | No | - | 7 | 23 | 29 | 31 | ~6 | 25 | 43 |
| GCF_902158545.1 | 100 | 0.27 | *Klebsiella pasteurii* 99.3349 % | No | No | - | 7 | 23 | 29 | 31 | ~6 | 25 | 43 |
| GCF_020116495.1 | 100 | 0.51 | *Klebsiella pasteurii* 99.3504 % | Yes | No | - | 7 | 23 | 38 | 39 | 82 | 25 | 115 |
| GCF_002186735.1 | 100 | 0.6 | *Klebsiella pasteurii* 99.3516 % | Yes | Yes | 311 | 17 | 32 | 29 | 44 | 92 | 25 | 77 |
| GCF_016616645.1 | 100 | 0.78 | *Klebsiella pasteurii* 99.3558 % | Yes | No | 351 | 7 | 23 | 23 | 39 | 69 | 25 | 43 |
| GCF_902163315.1 | 100 | 1.07 | *Klebsiella pasteurii* 99.3617 % | Yes | No | 351 | 7 | 23 | 23 | 39 | 69 | 25 | 43 |
| GCF_902163335.1 | 100 | 1.03 | *Klebsiella pasteurii* 99.3715 % | Yes | No | 351 | 7 | 23 | 23 | 39 | 69 | 25 | 43 |
| GCF_902158685.1 | 100 | 0.24 | *Klebsiella pasteurii* 99.3757 % | Yes | No | - | 17 | 16 | 23 | ~31 | 69 | 25 | 43 |
| GCF_015601345.1 | 100 | 0.58 | *Klebsiella pasteurii* 99.3796 % | Yes | No | - | 7 | 32 | 38 | 39 | 82 | 30 | 22 |
| GCF_902163325.1 | 100 | 1.03 | *Klebsiella pasteurii* 99.3812 % | Yes | No | 351 | 7 | 23 | 23 | 39 | 69 | 25 | 43 |
| GCF_003812845.1 | 100 | 0.52 | *Klebsiella pasteurii* 99.3818 % | Yes | No | 351 | 7 | 23 | 23 | 39 | 69 | 25 | 43 |
| GCF_902158665.1 | 100 | 0.19 | *Klebsiella pasteurii* 99.385 % | Yes | No | - | 7 | 23 | 23 | 31 | 69 | 25 | ~115 |
| GCF_902163345.1 | 100 | 1.09 | *Klebsiella pasteurii* 99.3884 % | Yes | No | 351 | 7 | 23 | 23 | 39 | 69 | 25 | 43 |
| GCF_902158705.1 | 100 | 0.63 | *Klebsiella pasteurii* 99.4068 % | Yes | No | 571 | 7 | 23 | 38 | 44 | 69 | 25 | 91 |
| GCF_018139045.1 | 100 | 0.22 | *Klebsiella pasteurii* 99.4105 % | Yes | No | 322 | 17 | 23 | 38 | 39 | 26 | 25 | 22 |
| GCF_902158715.1 | 100 | 0.27 | *Klebsiella pasteurii* 99.4155 % | Yes | No | 311 | 17 | 32 | 29 | 44 | 92 | 25 | 77 |
| GCF_902158585.1 | 100 | 0.24 | *Klebsiella pasteurii* 99.419 % | No | No | 311 | 17 | 32 | 29 | 44 | 92 | 25 | 77 |
| GCF_021228735.1 | 100 | 0.33 | *Klebsiella pasteurii* 99.4268 % | Yes | No | 300 | 13 | 23 | 38 | 39 | 69 | 25 | 22 |
| GCF_013266985.1 | 100 | 1.17 | *Klebsiella pasteurii* 99.4279 % | Yes | No | - | 7 | 23 | ~38 | 39 | 82 | 25 | 22 |
| GCF_009757395.1 | 100 | 0.25 | *Klebsiella pasteurii* 99.4291 % | Yes | No | 571 | 7 | 23 | 38 | 44 | 69 | 25 | 91 |
| GCF_000247915.1 | 100 | 0.37 | *Klebsiella pasteurii* 99.4303 % | Yes | No | 47 | 7 | 16 | 23 | 23 | 26 | 18 | 22 |
| GCF_019661035.1 | 100 | 0.85 | *Klebsiella pasteurii* 99.4304 % | Yes | No | - | 7 | 23 | 38 | 39 | 26 | 25 | ~22 |
| GCF_003261535.1 | 100 | 0.19 | *Klebsiella pasteurii* 99.4343 % | Yes | No | - | 7 | 23 | 58 | 39 | 64 | 25 | 43 |
| GCF_902158695.1 | 100 | 0.99 | *Klebsiella pasteurii* 99.4638 % | Yes | No | 569 | 7 | 23 | 38 | 31 | 82 | 25 | 43 |
| GCF_001065705.1 | 100 | 1.76 | *Klebsiella pasteurii* 99.4698 % | Yes | No | - | 13 | 23 | 38 | 39 | 69 | 25 | 115? |
| GCF_015679345.1 | 100 | 0.23 | *Klebsiella pasteurii* 99.4719 % | Yes | No | 47 | 7 | 16 | 23 | 23 | 26 | 18 | 22 |
| GCF_902158635.1 | 100 | 0.29 | *Klebsiella pasteurii* 99.4815 % | No | No | 320 | 17 | 16 | 38 | 39 | 82 | 25 | 43 |
| GCF_902158655.1 | 100 | 0.32 | *Klebsiella pasteurii* 99.488 % | Yes | No | 322 | 17 | 23 | 38 | 39 | 26 | 25 | 22 |
| GCF_016652955.1 | 100 | 3.06 | *Klebsiella pasteurii* 99.4983 % | Yes | No | 320 | 17 | 16 | 38 | 39 | 82 | 25 | 43 |
| GCF_001057685.1 | 100 | 0.52 | *Klebsiella pasteurii* 99.5026 % | Yes | No | 300 | 13 | 23 | 38 | 39 | 69 | 25 | 22 |
