## Supplementary Table 4 for "The plasmid-borne *hipBA* operon of *Klebsiella michiganensis* encodes a potent plasmid stabilization system"

**Supplementary Table 4. Summary of genomes predicted to encode chromosomal HipBAB**

| **Genome** | **HipBAB variant** | **Species** |
| --- | --- | --- |
| GCF_902162565 | 1 | *K. grimontii* |
| GCF_902162595 | 1 | *K. grimontii* |
| GCF_002192755 | 2 | *K. michiganensis* |
| GCF_016653015 | 2 | *K. michiganensis* |
| GCF_018439305 | 2 | *K. michiganensis* |
| GCF_018439315 | 2 | *K. michiganensis* |
| GCF_018439335 | 2 | *K. michiganensis* |
| GCF_018442305 | 2 | *K. michiganensis* |
| GCF_018442325 | 2 | *K. michiganensis* |
| GCF_018443125 | 2 | *K. michiganensis* |
| GCF_020117735 | 2 | *K. michiganensis* |
| GCF_902163235 | 2 | *K. michiganensis* |
| GCF_010093005 | 3 | *K. michiganensis* |
| GCF_009825595 | 4 | *K. michiganensis* |
| GCF_015721205 | 5 | *K. michiganensis* |
| GCF_902162465 | 6 | *K. michiganensis* |
| GCF_902162635 | 6 | *K. michiganensis* |
| GCF_902162685 | 6 | *K. michiganensis* |
| GCF_902162705 | 6 | *K. michiganensis* |
| GCF_000524315 | 7 | *K. michiganensis* |
| GCF_001753185 | 7 | *K. michiganensis* |
| GCF_002906435 | 7 | *K. michiganensis* |
| GCF_002918655 | 7 | *K. michiganensis* |
| GCF_002918695 | 7 | *K. michiganensis* |
| GCF_002919625 | 7 | *K. michiganensis* |
| GCF_014654995 | 7 | *K. michiganensis* |
| GCF_015721185 | 7 | *K. michiganensis* |
| GCF_015721425 | 7 | *K. michiganensis* |
| GCF_015721745 | 7 | *K. michiganensis* |
| GCF_020117595 | 7 | *K. michiganensis* |
| GCF_020117665 | 7 | *K. michiganensis* |
| GCF_020117715 | 7 | *K. michiganensis* |
| GCF_020118455 | 7 | *K. michiganensis* |
| GCF_020120585 | 7 | *K. michiganensis* |
| GCF_020121115 | 7 | *K. michiganensis* |
| GCF_022569835 | 7 | *K. michiganensis* |
| GCF_900083635 | 7 | *K. michiganensis* |
| GCF_902161885 | 7 | *K. michiganensis* |
| GCF_902161955 | 7 | *K. michiganensis* |
| GCF_902162365 | 7 | *K. michiganensis* |
| GCF_000240325 | 8 | *K. michiganensis* |
| GCF_000632415 | 8 | *K. michiganensis* |
| GCF_001066805 | 8 | *K. michiganensis* |
| GCF_001077175 | 8 | *K. michiganensis* |
| GCF_007910085 | 8 | *K. michiganensis* |
| GCF_010590505 | 8 | *K. michiganensis* |
| GCF_020118555 | 8 | *K. michiganensis* |
| GCF_020118655 | 8 | *K. michiganensis* |
| GCF_022359805 | 8 | *K. michiganensis* |
| GCF_022501085 | 8 | *K. michiganensis* |
| GCF_900083805 | 8 | *K. michiganensis* |
| GCF_900083835 | 8 | *K. michiganensis* |
| GCF_902166295 | 9 | *K. michiganensis* |
| GCF_902166305 | 9 | *K. michiganensis* |
