## Supplementary Table 5 for "The plasmid-borne *hipBA* operon of *Klebsiella michiganensis* encodes a potent plasmid stabilization system"

**Supplementary Table 5. HipA distance matrix (% identity)**

|  |  |  |  |  |  |  |  |  |  |  | **Cluster** | | | | |
| --- | --- | --- | --- | --- | --- | --- | --- | --- | --- | --- | --- | --- | --- | --- | --- |
|  | **Q63JY0** | **D3VDL4** | **P23874** | **S1ELM3** | **WP_038427096** | **A0A0H3C5Z1** | **A0A0H3CA27** | **Q8EIX3** | **A0A0H3CA56** | **ABP76633** | **1** | **2** | **3** | **4** | **5** |
| **Q63JY0 (*Burkholderia pseudomallei* K96243)** | 100.00 |  |  |  |  |  |  |  |  |  |  |  |  |  |  |
| **D3VDL4 (*Xenorhabdus nematophila* ATCC 19061)** | 43.15 | 100.00 |  |  |  |  |  |  |  |  |  |  |  |  |  |
| **P23874 (*Escherichia coli* K-12 (MG1655))** | 39.33 | 42.79 | 100.00 |  |  |  |  |  |  |  |  |  |  |  |  |
| **S1ELM3 (*Escherichia coli* BL21 (TaKaRa))** | 39.33 | 42.79 | 100.00 | 100.00 |  |  |  |  |  |  |  |  |  |  |  |
| **WP_038427096 (*Escherichia coli* Nissle 1917)** | 38.96 | 43.02 | 97.04 | 97.04 | 100.00 |  |  |  |  |  |  |  |  |  |  |
| **A0A0H3C5Z1 (*Caulobacter crescentus* NA1000)** | 28.41 | 27.59 | 23.87 | 23.87 | 23.93 | 100.00 |  |  |  |  |  |  |  |  |  |
| **A0A0H3CA27 (*Caulobacter crescentus* NA1000)** | 27.06 | 23.25 | 25.00 | 25.00 | 25.06 | 28.89 | 100.00 |  |  |  |  |  |  |  |  |
| **Q8EIX3 (*Shewanella oneidensis* MR-1)** | 22.46 | 21.62 | 25.83 | 25.83 | 25.22 | 26.17 | 22.98 | 100.00 |  |  |  |  |  |  |  |
| **A0A0H3CA56 (*****Caulobacter crescentus* NA1000)** | 15.19 | 15.45 | 15.35 | 15.35 | 15.78 | 16.77 | 17.66 | 14.92 | 100.00 |  |  |  |  |  |  |
| **ABP76633 (*Shewanella putrefaciens* CN-32)** | 13.14 | 15.11 | 13.15 | 13.15 | 12.97 | 15.42 | 13.58 | 14.91 | 16.67 | 100.00 |  |  |  |  |  |
| **Cluster 1** | 57.53 - 57.75 | 43.84 - 44.06 | 38.69 - 39.37 | 38.69 - 39.37 | 38.46 - 39.14 | 26.62 - 27.07 | 25.27 - 25.71 | 23.46 - 23.68 | 13.38 | 13.36 | 97.74 - 100.00 |  |  |  |  |
| **Cluster 2** | 56.63 | 43.84 | 39.59 | 39.59 | 39.37 | 27.74 | 24.84 | 23.03 | 14.01 | 13.36 | 83.94 - 84.62 | 100.00 |  |  |  |
| **Cluster 3** | 35.81 - 43.65 | 53.96 - 67.82 | 35.55 - 35.55 | 35.55 - 42.18 | 35.55 - 42.27 | 24.46 - 27.6 | 21.56 - 23.33 | 20.09 - 23.18 | 13.83 - 15.17 | 11.74 - 14.50 | 35.83 - 42.89 | 35.83 - 43.57 | 75.66 - 100.00 |  |  |
| **Cluster 4** | 15.35 - 15.57 | 16.16 - 16.38 | 16.41 - 16.41 | 16.41 - 16.41 | 16.63 - 16.63 | 16.89 - 16.89 | 14.29 - 14.50 | 14.69 - 14.90 | 28.19 - 28.63 | 13.68 - 14.32 | 16.20 - 17.27 | 15.57 - 15.99 | 12.59 - 14.59 | 95.25 - 100.00 |  |
| **Cluster 5** | 16.53 | 16.49 | 15.47 | 15.47 | 15.47 | 17.32 | 17.81 | 15.01 | 20.72 | 16.56 | 17.15 -17.78 | 16.95 - 16.95 | 15.70 - 17.51 | 18.87 - 19.31 | 100.00 |
