## Supplementary Table 6 for "The plasmid-borne *hipBA* operon of *Klebsiella michiganensis* encodes a potent plasmid stabilization system"

**Supplementary Table 6. HipB distance matrix (% identity)**

|  |  |  |  |  |  |  |  |  |  |  | **Cluster** | | | | |
| --- | --- | --- | --- | --- | --- | --- | --- | --- | --- | --- | --- | --- | --- | --- | --- |
|  | **Q63JY1** | **N1NPE2** | **P23873** | **WP_001301023** | **V0VAH2** | **A0A0H3C5I2** | **A0A0H3CAT0** | **Q8EIX4** | **A0A0H3CAW1** | **ABP76632** | **1** | **2** | **3** | **4** | **5** |
| **Q63JY1 (*Burkholderia pseudomallei* K96243)** | 100 |  |  |  |  |  |  |  |  |  |  |  |  |  |  |
| **N1NPE2 (*Xenorhabdus nematophila* ATCC 19061)** | 20.15 | 100 |  |  |  |  |  |  |  |  |  |  |  |  |  |
| **P23873 (*Escherichia coli* K-12 (MG1655))** | 17.91 | 36.36 | 100 |  |  |  |  |  |  |  |  |  |  |  |  |
| **WP_001301023 (*Escherichia coli* BL21 (TaKaRa))** | 17.91 | 36.36 | 100 | 100 |  |  |  |  |  |  |  |  |  |  |  |
| **V0VAH2 (*Escherichia coli* Nissle 1917)** | 19.4 | 36.36 | 95.45 | 95.45 | 100 |  |  |  |  |  |  |  |  |  |  |
| **A0A0H3C5I2 (*Caulobacter crescentus* NA1000)** | 19.32 | 20.73 | 17.44 | 17.44 | 17.44 | 100 |  |  |  |  |  |  |  |  |  |
| **A0A0H3CAT0 (*Caulobacter crescentus* NA1000)** | 6.34 | 7.75 | 5.99 | 5.99 | 5.99 | 14.29 | 100 |  |  |  |  |  |  |  |  |
| **Q8EIX4 (*Shewanella oneidensis* MR-1)** | 16.28 | 28.57 | 21.18 | 21.18 | 21.18 | 23.08 | 25.71 | 100 |  |  |  |  |  |  |  |
| **A0A0H3CAW1 (*Caulobacter crescentus* NA1000)** | 20.83 | 15.97 | 13.93 | 13.93 | 13.93 | 27.27 | 7.88 | 14.91 | 100 |  |  |  |  |  |  |
| **ABP76632 (*Shewanella putrefaciens* CN-32)** | 9.21 | 13.16 | 11.18 | 11.18 | 11.18 | 9.87 | 11.6 | 21.43 | 13.46 | 100 |  |  |  |  |  |
| **Cluster 1** | 35.82 - 36.57 | 25.51 - 26.53 | 21.43 - 22.45 | 21.43 - 22.45 | 22.45 - 23.47 | 19.54 - 20.69 | 5.28 - 5.63 | 21.05 - 22.11 | 16.94 - 18.55 | 10.53 - 11.18 | 90.91 - 100 |  |  |  |  |
| **Cluster 2** | 38.06 | 26.26 | 22.22 | 22.22 | 23.23 | 18.39 | 6.69 | 20.83 | 16.13 | 10.53 | 71 | 100 |  |  |  |
| **Cluster 3** | 20.15 | 63.1 | 37.5 | 37.5 | 38.64 | 24.39 | 7.04 | 23.26 | 19.17 | 13.16 | 23.23 | 23 | 100 |  |  |
| **Cluster 4** | 11.97 - 12.68 | 20.37 - 21.3 | 20.18 - 20.18 | 20.18 - 20.18 | 20.18 - 20.18 | 21.11 - 22.22 | 6.85 - 6.85 | 13.73 - 14.71 | 21.49 - 22.31 | 10.26 - 10.26 | 18.18 - 20 | 18.02 - 18.92 | 24.07 - 25 | 97.2 - 100 |  |
| **Cluster 5** | 11.69 | 13.11 | 13.11 | 13.11 | 13.11 | 19.42 | 7.17 | 16.35 | 22.22 | 10.26 | 16.26 - 17.07 | 16.26 | 11.38 | 20.16 | 100 |
